# Identification of *Chlamydia pneumoniae* and NLRP3 inflammasome activation in Alzheimer’s disease retina

**DOI:** 10.1101/2025.06.19.660619

**Authors:** Bhakta Prasad Gaire, Yosef Koronyo, Jean-Philippe Vit, Alexandre Hutton, Natalie Swerdlow, Dieu-Trang Fuchs, Altan Rentsendorj, Saba Shahin, Lalita Subedi, Edward Robinson, Alexander V. Ljubimov, Lon S. Schneider, Debra Hawes, Stuart L. Graham, Vivek K. Gupta, Mehdi Mirzaei, Keith L. Black, Jesse G. Meyer, Moshe Arditi, Timothy R. Crother, Maya Koronyo-Hamaoui

## Abstract

*Chlamydia pneumoniae* (Cp), an obligate intracellular bacterium, has been implicated in Alzheimer’s disease (AD), yet its role in retinal pathology remains unexplored. We analyzed postmortem tissues from 95 human donors and found 2.9-4.1-fold increases in Cp inclusions in AD retinas and brains, with no significant elevation in mild cognitive impairment (MCI). Proteomics revealed dysregulation of retinal and brain bacterial infection-related proteins and NLRP3 inflammasome pathways. NLRP3 expression was markedly elevated in MCI and AD retinas, and its activation was evident by increased N-terminal gasdermin D (NGSDMD) and mature interleukin-1β. Retinal Cp strongly correlated with Aβ_42_ and NLRP3 inflammasome components, which tightly linked to cleaved caspase-3-apoptotic and NGSDMD-pyroptotic cell death. Although retinal microgliosis was elevated in AD, Cp-associated microglia were reduced by 62%, suggesting impaired Cp phagocytosis. Higher retinal Cp burden correlated with APOEε4, Braak stage, and cognitive deficit. Machine learning identified retinal Cp or NLRP3 combined with Aβ_42_ as strong predictors of AD diagnosis, staging, and cognitive impairment. Our findings suggest that Cp infection contributes to AD dementia but not initiating pathology, whereas early NLRP3 activation may promote disease development, warranting studies on Cp’s role in AD pathogenesis and early antibiotic or inflammasome-targeted therapies.

## Introduction

Alzheimer’s disease (AD) is a debilitating neurodegenerative condition and the leading cause of dementia in the elderly, currently ranked as the seventh most common cause of death worldwide^1^. Affecting over 55 million individuals globally, with projections indicating a near threefold increase in cases by 2050, AD represents a major health crisis with profound social and economic implications^2^. The potential role of infectious agents in AD pathogenesis has gained increasing attention^3–9^, with *Chlamydia pneumoniae* (Cp), an obligate gram-negative bacterium primarily responsible for community-acquired pneumonia, emerging as a significant pathogen^10–13^. Genome-wide association studies (GWAS) in AD have identified genes associated with immune responses to pathogens, including the *Chlamydia* interactome, which overlaps with the AD hippocampal transcriptome and proteins involved in amyloid β-protein (Aβ) plaques and neurofibrillary tangles (NFTs)^14^. The GWAS findings show significant enrichment of AD-associated genes in pathways relevant to pathogen diversity, implicating their involvement in immune defense and pathogen resistance mechanisms^14^.

Cp inclusions have been detected in the postmortem brains of AD patients^15–19^. Immunohistochemical analyses revealed Cp proteins within vascular endothelial cells, microglia, astrocytes, and neurons, particularly in the frontal and temporal cortices, where Cp was localized near Aβ plaques and NFTs^15,17,18^. Additional studies identified Cp DNA in the cerebrospinal fluid (CSF), correlating with an increased risk of developing AD^20^, and Cp infection has been linked to the progression of dementia^21^. Notably, a recent nationwide cohort study in Taiwan demonstrated that antibiotic treatment targeting Cp significantly reduced the risk of AD onset^22^. These findings suggest that Cp infection may exacerbate AD pathology, and therapeutic strategies targeting Cp could potentially slow or mitigate AD progression.

Furthermore, Cp infection was shown to induce neuroinflammation and promote amyloid aggregation in murine models of AD, processes linked to AD progression^23–27^. Importantly, the nucleotide-binding oligomerization domain, leucine-rich repeat and pyrin domain-containing 3 (NLRP3) inflammasome, a key mediator of innate immunity that is activated in response to Cp infection^28,29^, has been implicated in AD pathogenesis^30–33^. NLRP3 inflammasome has been associated with Aβ-induced tauopathy in murine models^34,35^ and was upregulated in the brains of AD patients^36^. NLRP3 inflammasome and its components, NLRP3, pro-caspase-1, and adaptor protein ASC, trigger the release of pro-inflammatory cytokines, such as IL-1β, and lead to gasdermin D-mediated pyroptotic cell death^37–39^, probably contributing to the chronic inflammation and subsequent neurodegeneration that characterize AD^28,29^.

Histological, biochemical and *in vivo* imaging studies demonstrate that the pathological processes of AD manifest beyond the brain in the neurosensory retina^40^, a direct extension of the central nervous system that offers a unique opportunity for live observation^41–43^. In particular, the pathological hallmarks of AD, abnormal Aβ and tau aggregates, were identified in the retina of patients with mild cognitive impairment (MCI) and AD, along with vascular changes, inflammation, and neurodegeneration^44–76^. Hence, the unique accessibility of the retina to noninvasive imaging at high resolution and specificity may allow for early detection and monitoring of AD^40,43,77–82^. Despite emerging evidence indicating the presence of Cp and NLRP3 inflammasome activation in the AD brain^28–33^, the role of Cp infection and NLRP3-mediated inflammation in the retina remains unexplored. This knowledge gap underscored the need to investigate whether Cp infection and NLRP3 inflammasome activation occur in the AD retina, explore potential interactions during both early and advanced stages of AD, and examine their correlations with brain pathology and cognitive decline. These insights could unveil Cp and NLRP3 inflammasome activation as potential therapeutic targets and biomarkers of AD.

In this study, we investigated the presence and distribution of Cp inclusions in the retina and corresponding brain tissue from patients with AD dementia and MCI due to AD, compared with individuals with normal cognition (NC). We explored the relationship between retinal Cp burden and components of NLRP3 inflammasome activation, micro- and macrogliosis, as well as other AD-related retinal and brain pathologies, and cognitive function parameters. Additionally, we employed machine learning algorithms to predict various AD pathologies and cognitive dysfunction based on retinal markers of Cp, NLRP3, and cleaved caspase-3 (CCasp-3), either individually or in combination with retinal Aβ_42_, gliosis, and atrophy.

## Materials and methods

### Human eye and brain samples

Postmortem human eye globes and brain tissues were obtained from the Alzheimer’s Disease Research Center (ADRC) Neuropathology Core at the Department of Pathology in the University of Southern California (USC, Los Angeles, CA; IRB protocol HS-042071). In addition, eye globes were obtained from the National Disease Research Interchange (NDRI, Philadelphia, PA; under Cedars-Sinai IRB protocol Pro00019393). For a subset of patients and controls, we also obtained brain specimens from the ADRC Neuropathology Core at the University of California, Irvine (UCI IRB protocol HS#2014–1526). USC-ADRC, NDRI, and UCI ADRC maintain human tissue collection protocols that are approved by their managerial committees and subject to oversight by the National Institutes of Health and their respective institutional guidelines. All the histological procedures were conducted at Cedars-Sinai Medical Center under IRB protocols (Pro00053412 and Pro00019393). For histological examinations, 69 retinas were collected from deceased donors with confirmed AD (*n* = 34) or MCI due to AD (*n* = 14), and from age- and sex-matched deceased donors with NC (*n* = 21). In a subset of patients, paired brain tissues were also analyzed (*n* = 16). For mass spectrometry (MS) of retinal proteins, eyes were collected from another deceased donor cohort (*n* = 12) comprised of clinically and neuropathologically confirmed AD patients (*n* = 6) and matched NC controls (*n* = 6). For MS of brain proteins, fresh-frozen human brain tissue was obtained from an additional donor cohort (*n* = 18) of clinically and neuropathologically confirmed AD patients (*n* = 10) and matched NC controls (*n* = 8). The postmortem retinas and brain tissues were collected from clinically and neuropathologically confirmed MCI and AD patients, and age- and gender-matched NC individuals (**Table 1**, and **Suppl. Tables 1-5**). The human cohort used in this study exhibited no significant differences in age, sex, or post-mortem interval (PMI) hours. Patients’ confidentiality was maintained by de-identifying all tissue samples ensuring that donors could not be traced back.

**Table 1.** Demographic and neuropathological data on human donors for histological analysis.

|  | NC | MCI | AD | F | *p*-value |
| --- | --- | --- | --- | --- | --- |
| **Human donors** (n = 70) | 21 | 15 | 34 |  |  |
| female (%), male | 11F (52%), 10M | 7F (47%), 8M | 17F (50%), 17M |  |  |
| **Age at death** (years) | 84.57 ± 10.94 | 88.60 ± 6.25 | 86.18 ± 9.10 | 0.84 | 0.437 |
| **Race** (No.) | W(17); B/H(1/3) | W(13); B/H(1) | W(26); H/A(5/3) |  |  |
| **PMI** (hours) | 9.03 ± 5.88 | 12.23 ± 11.47 | 10.47 ± 11.85 | 0.39 | 0.676 |
| **MMSE Score** (n = 51) | 29.06 ± 1.78 | 22.00 ± 6.93 | 14.50 ± 7.83 | 26.29 | **<0.0001** |
| **CDR Score** (n = 57) | 0.17 ± 0.39 | 1.97 ± 1.18 | 2.32 ± 0.91 | 24.43 | **<0.0001** |
| **Sample size** | 12 | 15 | 34 |  |  |
| **Braak Stage** (%) | 0 (16.67) | 0 (6.67) | 0 (0) | 30.01 | **<0.0001** |
|  | I-II (41.67) | I-II (33.33) | I-II (0) |  |  |
|  | III-IV (33.33) | III-IV (33.33) | III-IV (23.53) |  |  |
|  | V-VI (8.33) | V-VI (26.66) | V-VI (76.47) |  |  |
| **ABC average**<br>(Amyloid, Braak,<br>CERAD) | 1.43 ± 0.79 | 1.98 ± 0.60 | 2.75 ± 0.38 | 30.72 | **<0.0001** |
| **Aβ Plaque** | 0.93 ± 1.05 | 1.64 ± 0.90 | 2.53 ± 1.00 | 12.89 | **<0.0001** |
| **NFT** | 0.54 ± 0.46 | 1.44 ± 0.97 | 2.36 ± 0.90 | 21.67 | **<0.0001** |
| **NT** | 0.77 ± 0.96 | 0.90 ± 0.78 | 2.01 ± 1.14 | 9.83 | **0.0002** |
| **Atrophy** | 0.55 ± 0.63 | 1.14 ± 0.91 | 1.99 ± 1.08 | 11.13 | **<0.0001** |

Paired brains with full neuropathological assessments were available for 61 human donors (n = 12 NC, n = 15 MCI, n = 34 AD). Values are presented as mean ± standard deviation. F and *p* values were determined by one-way ANOVA with Tukey's multiple comparisons test. *P* values presented in bold-italic type demonstrate significance. Mean ABC scores were determined as: **A**, Aβ plaque score modified from Thal; **B**, NFT severity score modified from Braak; **C**, neuritic plaque score modified from CERAD. Abbreviations: Aβ, amyloid beta; AD, Alzheimer's disease; A, Asian; B, Black; CDR, Clinical Dementia Rating; CERAD, Consortium to Establish a Registry for Alzheimer's Disease; NC, normal cognition; F, female; H, Hispanic; IHC, immunohistochemistry; M, male; MCI, mild cognitive impairment; MMSE, Mini- Mental State Examination; NFT, neurofibrillary tangle; NT, neuropil thread; PMI, post mortem interval; W, White.

### Clinical and neuropathological assessments

The detailed clinical and neuropathological assessments procedures are described in our recent publication^54,71^. In summary, clinical and neuropathological reports detailing patients’ neurological examinations, neuropsychological and cognitive assessments, were generously provided by ADRC system using the Unified Data Set^83^. The NDRI provided patients information, including gender, ethnicity, age at death, cause of death, medical background indicating AD, the presence or absence of dementia, and any accompanying medical conditions. Most cognitive assessments were conducted annually, typically within one year prior to death. In this study, we utilized cognitive scores assessed closest to the patient’s death. Three global indicators of cognitive status were used for clinical assessment: the Clinical Dementia Rating (CDR scores: 0 = normal; 0.5 = very mild impairment; 1 = mild dementia; 2 = moderate dementia; or 3 = severe dementia)^84^, Montreal Cognitive Assessment (MOCA scores: ≥26 = cognitively normal or <26 = cognitively impaired^85,86^, and the Mini-Mental State Examination (MMSE scores: normal cognition = 24-30; MCI = 20-23; moderate dementia = 10-19; or severe dementia ≤9)^87^.

The assessment of cerebral Aβ burden comprises the analysis of diffuse and neuritic plaques (including both the immature and mature forms), along with amyloid angiopathy, NFTs, neuritic threads (NTs), granulovacuolar degeneration, Lewy bodies, Hirano bodies, Pick bodies, balloon cells, neuronal loss, microvascular changes, and gliosis. These evaluations were conducted across different brain regions, notably in the hippocampus, the entorhinal cortex, the superior frontal gyrus in the frontal lobe, the superior temporal gyrus in the temporal lobe, the superior parietal lobule in the parietal lobe, the primary visual cortex, and the visual association area in the occipital lobe. All brain samples were uniformly collected by a neuropathologist.

Formalin-fixed, paraffin-embedded brain sections were used to determine the severity of amyloid plaques and NFTs in the brain using anti Aβ monoclonal antibody (mAb) clone 4G8, anti phospho-tau mAb clone AT8, Thioflavin-S (ThioS), and Gallyas silver staining. Two neuropathologists independently rated the burden of Aβ, NFTs, and NTs on a scale of 0, 1, 3, and 5 [(0 = none, 1 = sparse (0-5), 3 = moderate (6-20), 5 = abundant/frequent (21-30 or above), n.a. = not applicable)], with the final score being the average of the two readings. The final diagnosis included AD neuropathologic change. The Aβ plaque scoring system, was adapted from Thal et al., (A0 = no Aβ or amyloid plaques, A1 = Thal phase 1 or 2, A2 = Thal phase 3, and A3 = Thal phase 4 or 5)^88^. NFT staging was adjusted from Braak for silver-based histochemistry or p-tau immunohistochemistry (B0 = No NFTs, B1 = Braak stage I or II, B2 = Braak stage III or IV, B3 = Braak stage V or VI)^89^. The neuritic plaque score was adapted from CERAD (C0 = no neuritic plaques, C1 = CERAD sparse, C2 = CERAD moderate, C3 = CERAD frequent)^90^. Additional evaluations included neuronal loss, gliosis, granulovacuolar degeneration, Hirano bodies, Lewy bodies, Pick bodies, and balloon cells using hematoxylin and eosin staining, with scores of 0 for absent and 1 for present. Amyloid angiopathy was classified into 4 grades: Grade I indicates amyloid around normal/atrophic smooth muscle cells of vessels; Grade II shows media replaced by amyloid without blood leakage; Grade III involves extensive amyloid deposition with vessel wall fragmentation and perivascular leakage; Grade IV includes extensive amyloid deposition with fibrinoid necrosis, microaneurysms, mural thrombi, lumen inflammation, and perivascular neuritis.

### Collection and processing of eyes and brain cortical tissues

Donor eyes were collected and preserved within an average of 10 hours after time of death. These eyes were either preserved in Optisol-GS media (Bausch & Lomb, 50006-OPT), snap frozen upon delivery and stored at -80°C, or punctured once at the limbus and fixed in 10% neutral buffered formalin or 4% paraformaldehyde (PFA) and stored at 4°C. Brain tissues (hippocampus, Brodmann Area 9 of the prefrontal cortex and Brodmann area 17 of the primary visual cortex) were collected from the same donors, snap frozen, and stored at -80°C. For MS, fresh-frozen human brain (hippocampus, medial temporal gyrus, and cerebellum) tissues were obtained from an additional donor cohort. The same tissue collection and processing procedures were consistently applied regardless of whether the human donor eyes and brains were sourced from USC-ADRC, NDRI and UCI-ADRC.

### Preparation of retinal and brain cross-sections

Fresh eyes those preserved in Optisol-GS and fixed eyes were dissected on ice by removing the anterior segments to form eyecups. The vitreous humor was thoroughly removed manually. Retinas were then carefully dissected, detached from the choroid, and prepared as flatmounts following established procedures^53,54,71^. Geometric regions of the four retinal quadrants were defined for both left and right eyes by identifying the macula, optic disc, and blood vessels. Flatmount-strips, measuring 2-3 mm in width from fixed retina and 5 mm in width from fresh retina, spanning from ora serrata to optic disc, were dissected along the margins of these quadrants to create four strips: superior-temporal (ST), inferior-temporal (TI), inferior-nasal (IN), and superior-nasal (NS). Flatmount-strips from fixed retina were initially embedded in paraffin, then rotated 90° horizontally and re-embedded in paraffin block. Retinal strips, approximately 2-2.5 cm in length, encompassing central, mid, and far retinal subregions, were sectioned into 7-µm thick slices and mounted on microscope slides treated with 3-aminopropyltriethoxysilane (APES, Sigma A3648). Flatmount strips from fresh-frozen retinas were stored at − 80°C for MS protein analysis. Fresh-frozen paired brain tissues were fixed in 4% PFA for 16 hours at 4°C, following paraffin embedding. They were sectioned (10-μm thick) and mounted on microscope slides treated with APES. This sample preparation technique allowed for extensive and consistent access to retinal quadrants, layers, and pathological subregions.

### Immunohistochemical analysis

Before IHC procedure, paraffin-embedded cross-section slides were deparaffinized with 100% xylene twice (10 min each), rehydrated with decreasing concentrations of ethanol (100% to 70%), and washed with distilled water followed by PBS. Following deparaffinization, retinal and brain cross-sections were treated with antigen retrieval solution (pH 6.1; S1699, DAKO) at 99°C for 1 hour, washed with PBS, and then treated with 70% formic acid (ACROS) for 10 minutes at room temperature (RT).

For peroxidase-based labelling, we used a Vectastain Elite ABC HRP kit (Vector, PK-6102, Peroxidase Mouse IgG) according to the manufacturer’s instructions. In summary, following incubation with 3% H_2_O_2_ for 20 minutes, tissues were washed with PBS and incubated with blocking serum containing 0.25% Triton X-100 (Sigma, T8787) for 45 minutes at RT. Primary antibodies against Cp (**Suppl. Table 6**) were diluted in PBS containing blocking serum and incubated overnight at 4°C. The following day, the tissues were rinsed three times with PBS, incubated for 30 minutes at 37°C with secondary antibody, rinsed three times with PBS and incubated with ABC reagent for 30 minutes at RT. After washing with PBS, Cp inclusions in the brain or retinal tissue sections were visualized with 3,3′-diaminobenzidine (DAB) substrate (DAKO K3468). Hematoxylin counterstaining was performed followed by mounting with Paramount aqueous mounting medium (DAKO, S3025). Routine controls were processed using identical protocols while omitting the primary Ab to assess nonspecific labeling.

For fluorescence-based immunostaining, sections were first treated with a blocking solution (DAKO X0909) containing 0.25% Triton X-100 (Sigma, T8787), followed by an overnight incubation with primary antibodies (**Suppl. Table 6**) at 4°C. The next day, following wash with PBS sections were incubated with fluorophore conjugated secondary antibodies (**Suppl. Table 6**) for 1 hour at RT. Tissue sections were mounted using ProLong Gold Antifade Mountant with DAPI (Thermo Fisher, #P36935). Control sections processed without the primary antibodies were used to assess nonspecific labeling. To minimize background autofluorescence, brain sections were treated with 1X True Black (Biotium, #23007), diluted in 70% ethanol (v/v) for 40 seconds at RT before the application of primary antibodies.

### Microscopy and quantitative immunohistochemistry

Images were acquired using a Carl Zeiss Axio Imager Z1 fluorescence microscope with ZEN 2.6 blue edition software (Carl Zeiss MicroImaging, Inc.) equipped with ApoTome, AxioCam MRm, and AxioCam HRc cameras, at a resolution of 1388 × 1040 pixels, 6.45 µm × 6.45 µm pixel size, and dynamic range of >1:2200, which delivers low-noise images due to a Peltier-cooled sensor. Multi-channel image acquisition was used to create images with several channels. Images were repeatedly captured at the same focal planes with the same exposure time for each marker and human donor. Images were captured at 20×, 40× (at respective resolution of 0.5 and 0.25 µm), 63×, and 100× objectives for different purposes. For representative imaging, Z-stack images were repeatedly captured at same tissue thickness using a Carl Zeiss 780 Confocal microscope (Carl Zeiss MicroImaging, Inc.). We randomly acquired images, 3 from the central, 4 from the mid, and 3 from the far retinal subregions, for analytical purposes (as shown in **Fig. 1A**). Thickness measurements (µm) were manually performed using Axiovision Rel. 4.8 software. Retinal thickness assessments were taken from the ILM through the OLM. Images were exported to ImageJ2/Fiji (version 2.14.0; NIH) to analyze parameters of interest. Acquired images were converted to grayscale and standardized to baseline by using a histogram-based threshold in ImageJ2. The images were then submitted to ImageJ2 particle analysis for each biomarker to determine total and % IR area. For each biomarker, the IR area was determined using the same threshold percentage from the baseline in ImageJ2 with the same percentage threshold setting for all diagnostic groups. Throughout the analysis process, the researchers were blinded to each patient’s diagnosis.

**Figure 1.**
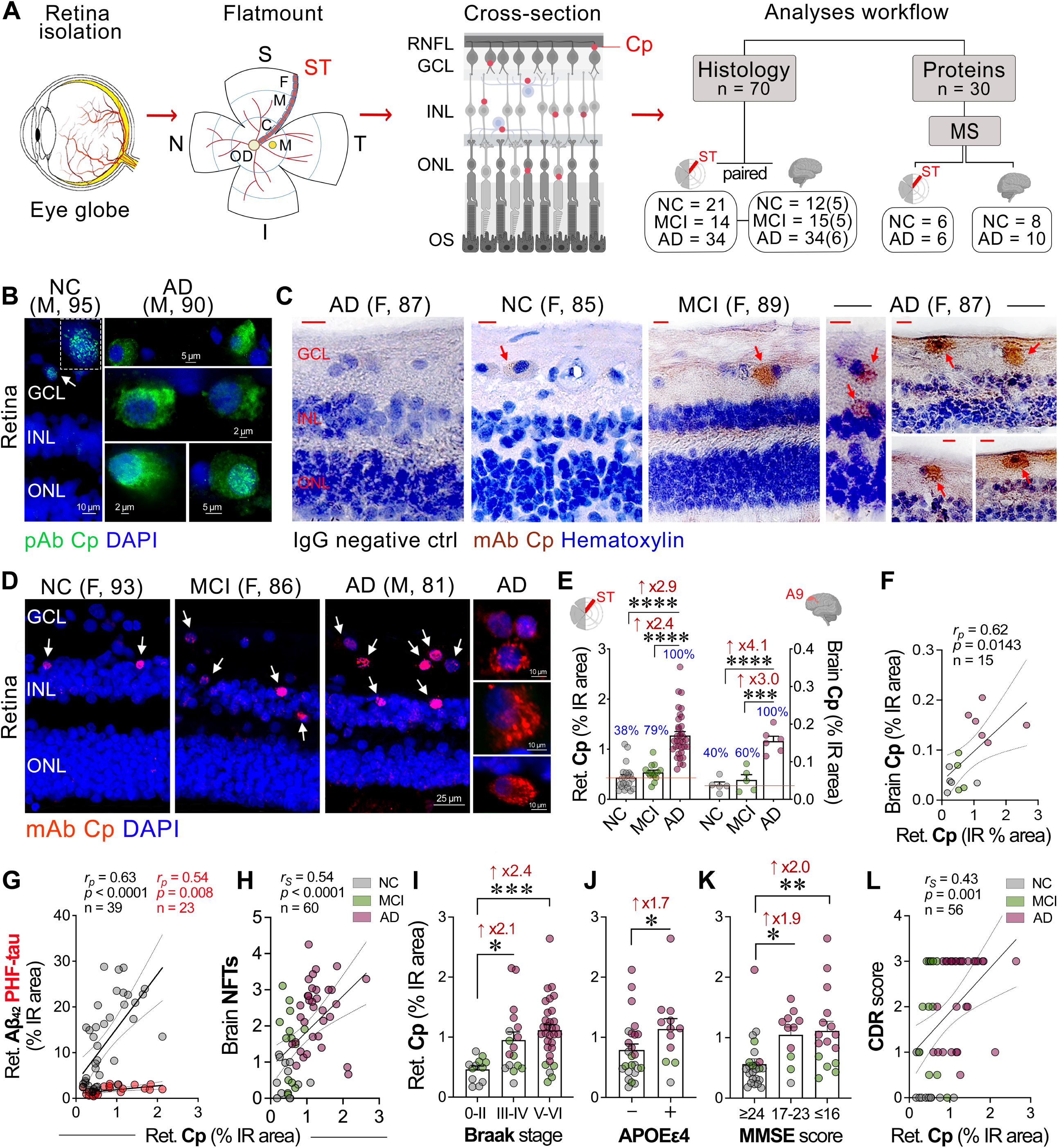
Identification of *Chlamydia pneumoniae* (Cp) inclusions in retinas and brains from MCI and AD patients: correlations with disease status. **A**. Illustration of retinal isolation and cross-section preparation from donor eyes. Retina was geometrically divided into four quadrants: superior (S), temporal (T), inferior (I), and nasal (N). A retinal strip (∼2-3 mm), from the superior-temporal (ST) region extending from the optic disc (OD) to the ora serrata, was isolated and processed into retinal cross-sections for histopathological and proteomics analyses. This strip was further divided into subregions: central (C), mid-periphery (M), and far periphery (F). Red dots in the schematic retinal cross-section represent the presence of Cp inclusions identified across retinal cell layers. Right panel depicts the analyses workflow for the cohort size in each experiment. The numbers in parenthesis for brain histology indicate the subset of subjects analyzed for Cp immunoreactivity. **B**. Representative immunofluorescence micrographs of retinal cross-sections stained with pAb against Cp, with cytosolic inclusions shown in green (arrows), in retinal GCL of a human donor with AD compared with NC controls. **C**. Representative micrographs of peroxidase-based immunohistochemistry analysis of Cp inclusions (brown) in cells (nuclei; hematoxylin) stained with mAb in retinal cross-sections from MCI and AD patients versus NC controls (Left image, IgG negative control in the retina of an AD patient). Scale bars: 10 µm. **D**. Representative immunofluorescence images of retinal cross-sections from MCI and AD patients versus NC controls, indicated the presence of specific Cp inclusions (red; white arrows), using the mAb against Cp, in cells (nuclei, DAPI) across several retinal layers (right images are of higher magnification showing cytosolic Cp inclusions in AD retinas. **E**. Bar graphs display the quantitative analyses of retinal and paired-brain Cp-immunoreactive (IR) percentage area in age- and sex-matched human donors with MCI (due to AD) and AD dementia versus NC controls [for retinal analysis (n = 21 NC, 14 MCI, and 34 AD), for paired-brain analysis (n = 5 NC, 5 MCI, and 6 AD)]. The analyzed ST retina and the Area 9 (located in the dorsolateral prefrontal cortex) from the paired brain were demarcated with red color. The percentage of Cp-positive individuals was determined by having a higher value than the mean % IR area of Cp in the NC group (red line), for each CNS tissue. **F**. Pearson correlation (*r_p_*) analysis between retinal and paired-brain Cp burdens. **G, H**. Pearson correlation (*r_p_*) analyses between retinal Cp burden and (G) retinal Aβ_42_ (grey dots) and retinal paired helical filament (PHF)-Tau (red dots) % IR areas, and (H) brain NFT severity score. **I-K**. Retinal Cp burden per (I) Braak stage stratification, 0-II (n = 12), III-IV (n = 17), and V-VI (n = 31), (J) individuals carriers (n = 12) or non-carriers (n = 25) of APOE ɛ4 allele(s), and (K) mini-mental state examination (MMSE) cognitive score categories, ≥ 24 (n = 24), ≥ 17-23 (n = 9), ≤ 16 (n = 14). **L**. Pearson’s correlation (*r_p_*) analysis between retinal Cp burden and clinical dementia rating (CDR) score. Data from individual subjects (circles) as well as group means ± SEMs are shown. Fold changes are indicated in red. \**p* < 0.05, \*\**p* < 0.01, \*\*\**p* < 0.001, and \*\*\*\**p* < 0.0001, by one-way ANOVA and Tukey’s post hoc multiple comparison test or by two-tailed paired unpaired Student’s t test for two-group comparison.

#### Quantitative analysis of Cp-associated microglia

Three distinct stages of retinal IBA1^+^ microglial involvement in the phagocytosis of Cp/Cp-infected cells were analyzed. The ‘recognition’ stage was defined as the count of IBA1^+^ microglia in direct contact with Cp/Cp-infected cells (<50% of cell circumference). The ‘engulfment’ stage was determined as the count of IBA1^+^ microglia whose processes surrounded ≥50% of the Cp/Cp-infected cells. The ‘ingestion’ stage refers to the count of IBA1^+^ microglia in which Cp/Cp-infected cells were fully internalized, evidenced by co-localization of Cp (red) and IBA1 (green) as yellow in merged images. Microglial cells participating in these three stages were classified as ‘Cp-associated microglia’ (CAM). The relative contribution of microglia to Cp/Cp-infected cell recognition and uptake was quantified by calculating the proportion of CAM cells relative to the total IBA1^+^ microglial population.

### Biochemical determination of Aβ_1–42_ levels by sandwich ELISA in human retina

Fresh-frozen human retinal strips from the temporal hemisphere (ST, IT) were homogenized (1 mg tissue/10 µl buffer) in cold homogenizing buffer (100 mM TEA Bromide [Sigma, 241059], 1% sodium deoxycholate [SDC; Sigma, D6750], and 1x Protease Inhibitor cocktail set I [Calbiochem, 539,131]). Retinal homogenates were sonicated (Qsonica sonicator with an M-Tip probe, amplitude 4, 6 W, for 90 sec, in bouts of 15 sec) while the ultrasonic probe was positioned inside the sample tube that was placed in ice water. The amount of retinal Aβ_1–42_ was determined using an anti-human Aβ_1–42_ end-specific sandwich ELISA kit (Thermo Fisher, KHB3441) and normalized to the total protein concentrations (Thermo Fisher Scientific).

### Western blot analysis

Fresh frozen postmortem human retinal tissues were homogenized in radioimmunoprecipitation assay (RIPA) buffer supplemented with protease and phosphatase inhibitors. Protein concentrations were determined using the bicinchoninic acid protein assay kit (Pierce^TM^). Equal amounts of protein (30 µg per sample) were separated on 4–20% precast polyacrylamide gels (Bio-Rad, catalog #4561094) and transferred to a polyvinylidene difluoride membrane. The membrane was then blocked with 2.5% bovine serum albumin in 1X TBS-T (Tris-buffered saline with 0.1% Tween-20) for 1 hour at room temperature, followed by overnight incubation at 4 °C with primary antibodies: anti-IL-1β (1:1000, Abcam, catalog #ab9722) and anti-GAPDH (1:1000, Millipore Sigma, catalog #G8795). After washing with 1X TBS-T, the membrane was incubated with fluorescently labeled secondary antibodies, and bands were detected using the LI-COR Odyssey imaging system. Band intensities were quantified using Image Studio software (LI-COR), and relative protein expression levels were calculated by normalizing target protein signals to GAPDH.

### Proteome analysis by mass spectrometry (MS)

#### Preparation of retinal and brain samples from NC and AD donors

Frozen brain and retina tissue samples were processed for mass spectrometry (MS) analysis by the University of Queensland, in accordance with approval from the institution’s Human Research Ethics Committee (approval number #2017000490). For the brain analysis, frozen aliquots from the hippocampus, medial temporal gyrus, and cerebellum were used. These tissues were transferred into Precellys homogenization tubes (Bertin Technologies), homogenized in liquid nitrogen, and lysed in ice-cold T-PER extraction buffer (Thermo Scientific) containing protease and phosphatase inhibitors. Tissue lysates were cleared of debris by ultracentrifugation at 100,000 g for 60 minutes at 4°C. Retinal temporal hemisphere (ST and TI) tissues were homogenized in a buffer containing 100 mM TEA Bromide (Sigma, 241059), 1% SDC (Sigma, D6750), and a protease inhibitor cocktail (Calbiochem 539131) using a Qsonica sonicator with an M-Tip probe (amplitude 4, 6 W, for 90 seconds, with the sonication pulse stopped every 15 seconds to allow cooling for 10 seconds). Insoluble materials were removed by centrifugation at 15,000 g for 10 minutes at 4°C. Protein concentrations of brain and retinal lysates were determined using the Bradford assay (Bio-Rad Laboratories). Extracted proteins were reduced using 5 mM DTT and alkylated with 10 mM iodoacetamide. Protein concentration was further verified using a BCA assay kit (Pierce). Dual digestion was performed on 150 µg of protein, initially using Lys-C (Wako, Japan) at a 1:100 enzyme ratio overnight at RT, followed by trypsin (Promega) at a 1:100 enzyme ratio overnight at 37°C. Detailed MS proteome analysis methods, including tandem mass tag (TMT) labeling and nanoflow liquid chromatography electrospray ionization tandem MS (nano LC–ESI–MS/MS), were as described in our previous publication^54^.

#### Database Searching, Peptide Quantification, and Statistical Analysis

Raw data files were processed using Proteome Discoverer V2.1 software (Thermo Scientific) and Mascot (Matrix Science, UK). Data were matched against the reviewed SwissProt *Homo sapiens* protein database. The MS1 tolerance was set to ± 10 ppm and the MS/MS tolerance to 0.02 Da. Carbamidomethyl (C) was set as a static modification, while TMT10-plex (N-term, K), oxidation (M), deamidation (N, Q), Glu->pyro-Glu (N-term E), Gln->pyro-Glu (N-term Q), and acetylation (Protein N-Terminus) were set as dynamic modifications. The percolator algorithm was used to discriminate correct from incorrect peptide-spectrum matches and to calculate statistics including adjusted *p* value with the Benjamini-Hochberg procedure for controlling the false discovery rate (FDR) when conducting multiple hypothesis tests and posterior error probabilities. Search results were further filtered to retain protein with an FDR of <1%, and only master proteins assigned via the protein grouping algorithm were retained. Proteins were further analyzed using the TMTPrepPro analysis pipeline^91^. TMTPrepPro scripts are implemented in the R programming language and are available as an R package, which was accessed through a graphic user interface provided by a local Gene Pattern server. In pairwise comparison tests, the relative quantitation of protein abundance was derived from the ratio of the TMT label S/N detected in each condition (AD versus NC), and differentially expressed proteins (DEPs) were identified based on Student’s t-tests between AD and NC group ratios (log-transformed). Overall fold changes were calculated as the geometric means of the respective ratios. Differential expression was defined by both a fold change (|FC| > 1.2) and *p*-value threshold (*p*<0.05), by Student t-test.

#### Functional Network and Computational Analysis

Gene Ontology (GO) analysis of differentially expressed proteins (DEPs) was performed in Metascape (https://metascape.org/; cutoffs: overlap ≥ 3, *p*-value <0.01, enrichment ≥ 1.5) and included the GO Biological Processes, Reactome, Kyoto Encyclopedia of Genes and Genomes (KEGG) and WikiPathways databases. Enrichment analysis results are reported with z-scores, unadjusted *p*-values and Benjamini-Hochberg adjusted *p*-values to control the FDR. Networks of pathways related to infection, neuroinflammation, immune response and cell death were created in Metascape, then loaded and modified in Cytoscape 3.10.2 (https://cytoscape.org/). Protein interaction networks were generated in String v12.0 and modified in Cytoscape. Volcano plots representing expression changes [log_2_(FC)] and significance level [-log_10_(*p*)] in AD versus NC were created using Prism 10.3.1 (GraphPad) and included human proteins linked to *Chlamydia* inclusion membrane. The list of human proteins interacting with *Chlamydia* inclusions (termed ‘*Chlamydia* interactome’) was determined from four original studies and a meta-analysis study^92–96^. Heatmaps corresponding to the protein expression level of select proteins in the retina of the 6 NC individuals and the 6 AD patients, standardized by unit variance scaling, were generated in ClustVis (https://biit.cs.ut.ee/clustvis/). Chord diagrams representing the association of DEPs with select functional pathways were created in Circos online (https://mk.bcgsc.ca/tableviewer/). The mass spectrometry proteomics data have been deposited to the ProteomeXchange Consortium via the PRIDE^97^ partner repository with the dataset identifier PXD040225^97^.

### Statistical analysis

GraphPad Prism 10.3.1 was used for statistical analyses. A comparison of three or more groups was performed using one-or two-way ANOVA followed by Tukey’s multiple comparison post-hoc test. Two-group comparisons were analyzed using a two-tailed unpaired Student’s t-test. The statistical association between two or more variables was determined by Pearson’s (*r_p_* for parametric) or Spearman (*r_s_* for non-parametric) correlation coefficient test (Gaussian-distributed variables). Pair-wise Pearson’s (*r_p_*) or Spearman’s (*r_s_*) coefficients with unadjusted *p* values were used to indicate the direction and strength of the linear relationship between the two variables, whereas adjusted *p* values were used for multivariate correlation analyses, as specified in the respective figure legends. The correlation strength was defined by coefficient (*r*) value as follows: very strong 0.80-1.00, strong 0.60-0.79, moderate 0.40-0.59, and weak 0.20-0.39. Required sample sizes for comparisons of two group (differential mean) were calculated using the nQUERY t-test model, assuming a two-sided α level of 0.05, 80% power, and unequal variances, with the means and common standard deviations for the different parameters. Results are expressed as mean ± standard error of the mean (SEM), with *p <* 0.05 considered significant. Fold changes (FC) and corresponding 95% confidence intervals (CI) were calculated.

### Machine learning prediction

#### Prediction of brain measures and diagnosis

Two sets of machine learning models were trained to predict brain-based measures using information from the retina: one set of regressors for predicting continuous measures, and a set of classifiers for predicting disease diagnosis. Due to the sample size and degree of missingness across the variables of interest, random forests^98^ with 80 estimators were used for both the regression and classification tasks. The data were split into two portions stratified by diagnosis: 80% for model training, and 20% for testing. The 20% test set was left untouched until after the final model was selected and was only used to evaluate model performance. The 80% training set consisted of 56 subjects (17/12/27 NC/MCI/AD), and the 20% test set consisted of 14 subjects (4/3/7 NC/MCI/AD). All models were evaluated using 5-repeated 2-fold cross-validation. The 2-fold cross-validation consists of randomly splitting the training dataset into two parts stratified by diagnosis, training the model on one and evaluating on the other, then swapping the two parts and repeating. This is repeated 5 times to obtain a distribution of predicted performance. The data were processed, and models were trained using a combination of Scikit-learn^99^, Numpy^100^, Pandas^101^, Scipy^102^, and custom Python 3.11 code (https://github.com/xomicsdatascience/Retinal_Alzheimer_Prediction).

#### Prediction of brain-based measures

Random forest models were used to predict brain Aβ plaques, brain NFTs, brain gliosis, brain atrophy, Braak stage, and ABC severity scores, as well as MMSE and MOCA cognitive scores. Different subsets of retinal features were used to train a model, combining one of Cp, NLRP3, CCasp3 with Aβ_42_, gliosis (IBA1, vimentin, GFAP), atrophy index, or by themselves, in addition to using Aβ_42_ alone as a reference. For each target, these results in 13 models were used for evaluation. Due to concerns over the variance of predictive performance due to the sample size, every model was evaluated using a 5-repeated 2-fold cross-validation to obtain the coefficient of determination (r^2^).

#### Prediction of disease diagnosis

Similar to the prediction of the brain biomarkers, random forest models were used to predict disease diagnosis from information gathered from the retina. Each subject was identified by one of three disease statuses: MCI (due to AD), AD (dementia), or NC. To obtain the receiver operating characteristic (ROC) curve for this non-binary task, we averaged the ROC curves across disease status. For each of the reported models, we included the area under the ROC curve (AUC) as an overall summary of the model performance. Model performance was compared with the null distribution by obtaining the AUC distribution generated from training dummy classifiers and comparing the model’s AUC value to the distribution. Given that the model performance distributions were sufficiently broad to make it unclear whether the reported models were meaningfully different, we compared the AUC distributions from the 5-repeated 2-fold cross-validation across different models using a Wilcoxon signed-rank test^103^, with the obtained *p* values adjusted for multiple comparisons with Benjamini-Hochberg procedure^104^.

## Results

To investigate Cp infection and the association with neuroinflammation and neurodegeneration in the AD retina, we analyzed retinal and brain tissues from a cohort of 95 patients: 46 with AD dementia (mean age ± SD: 85.98 ± 10.33 years, 25 females/21 males), 15 with MCI due to AD (88.60 ± 6.25 years, 7 females/8 males), and 34 NC individuals (84.88 ± 10.09 years, 19 females/15 males). There were no significant differences in age, sex, or postmortem interval across groups. Demographics, clinical, and neuropathological data are detailed in **Table 1** and **Suppl. Tables 1-5**. To study the protein expression profile related to Cp infection, we conducted MS-based proteome analysis on the temporal retina and cortex from human donors (n = 30) whose postmortem tissues were promptly processed and fresh frozen for protein isolation and analysis^54^. Our IHC analyses were focused on the superior-temporal (ST) retinal subregion and the dorsolateral prefrontal cerebral cortex (A9), given their strong association with AD pathology^43,53,105,106^ (**Fig. 1A**).

### 1. Identification of Cp inclusions in the AD retina, correlations with retinal Aβ_42_ burden, brain Cp, APOE genotype, and dementia status

To explore the presence and distribution of Cp inclusions in the human retina, we initially performed IHC analysis using an anti-Cp polyclonal antibody (pAb), which can also cross-react with other known *Chlamydia* species (e.g., *C. trachomatis* and *C. psittaci*). Utilizing this antibody, Cp-positive inclusions were identified predominantly in retinal ganglion cell layer (GCL) and inner nuclear layer (INL), as visualized through both fluorescence-based (**Fig. 1B**) and peroxidase-based (**Suppl. Fig. 1A, B**) immunolabeling. The retinas of AD patients compared with those of NC controls exhibited more frequent Cp-positive retinal ganglion cells (RGCs). Most of the Cp inclusions were cytosolic, whereas few inclusions were also detected in the peri-nucleus or nucleus and colocalized with DAPI. We further confirmed the presence of Cp inclusions in retinal cross-sections with a Cp-specific monoclonal antibody (mAb), which does not cross-react with other *Chlamydia* species (**Fig. 1C, D**; extended data in **Suppl. Fig. 1C, D**). With this mAb, Cp inclusions were mainly observed as cytosolic puncta aggregates in the MCI and AD retina, which resembles the patterns observed using the anti-Cp pAb. Cp inclusions were also detected in the corresponding cerebral cortices of AD patients (**Suppl. Fig. 1C, E**), identifying typical intracellular inclusions, similar to previously reported Cp inclusion patterns in the AD brain^17,18^.

Quantitative analysis of retinal and brain Cp^+^ immunoreactive (IR, mAb) area revealed significant 2.9- and 4.1-fold increases in Cp inclusions, respectively, in AD patients compared with NC controls (**Fig. 1E**; *p*<0.0001). No significant difference in Cp load was observed between NC and MCI retinas and brains, indicating that the expansion of Cp infection likely occurs later in disease progression, during the clinical dementia stages of AD. Notably, the percentage of individuals with retinal or brain Cp-positive values, relative to the average Cp IR area in the NC group, was 38-40% among NC controls, 60-79% of MCI patients, and 100% (all) of AD dementia patients (**Fig. 1E**). Gaussian distribution curves for both retinal and brain Cp levels revealed a strong overlap between the NC and MCI groups compared with the AD group (**Suppl. Fig. 1F, G**). In general, Cp patterns in the retina were comparable to those observed in the respective brains (**Fig. 1C, D** and **Suppl. Fig. 1C-E**), with a strong correlation between retinal and brain Cp loads (**Fig. 1F**; *r_p_* = 0.62, *p* = 0.0143).

We next explored the geometrical distribution of Cp infection and found that Cp was uniformly distributed across the central (C), mid-peripheral (M), and far-peripheral (F) subregions of the ST retina (**Suppl. Fig. 2A, B**; Perinuclear and nuclear Cp mAb fluorescence shows a pink signal colocalized with DAPI). Cp inclusions were predominantly detected in the inner retinal layers (GCL, IPL, and INL), with more frequent cytosolic staining of Cp inclusions observed in the MCI and AD retinas versus the NC retinas. No significant differences in the levels of retinal Cp were detected between male and female subjects for each diagnostic group (**Suppl. Fig. 2C**), suggesting no sex-specific dimorphism for retinal Cp burden.

Previous studies demonstrated that Cp infection induces amyloid accumulation in the mouse brain^24,25^, linking Cp infection to amyloid deposition and Alzheimer’s-related pathology. Consistent with this, we found a strong correlation between retinal Cp burden and retinal amyloidosis (**Fig. 1G** and **Suppl. Table 7**; Aβ_42_: *r_p_* = 0.63, *p*<0.0001, Aβ_40_: *r_p_* = 0.65, *p* = 0.0014), with no correlation with intracellular Aβ oligomers (**Suppl. Table 7**), indicating a specific association with the extracellular plaque-dominant Aβ_42_ and vascular-dominant Aβ_40_ alloforms. Our analysis also revealed moderate-to-weak significant correlations between retinal Cp burden and markers of retinal tauopathy, such as paired-helical filament (PHF-1) of tau (*r_p_* = 0.54, *p* = 0.0085, **Fig. 1G**), pS396-tau (*r_p_* = 0.38, *p* = 0.0116), T22^+^ oligo-tau (*r_p_* = 0.43, *p* = 0.0040), and citrullinated tau (CitR_209_: *r_p_* = 0.48, *p* = 0.0028; **Suppl. Table 7**). Retinal Cp burden did not correlate with retinal AT8-tau or MC-1^+^ mature tau tangles (**Suppl. Table 7**). These data indicate that Cp inclusions in the MCI and AD retina appear dominant in the inner cell layers, closely interact with amyloidogenic Aβ, and modestly associate with certain retinal tau isoforms while not with others.

Next, we determined whether retinal Cp burden associates with AD-related brain pathology, APOE ɛ4 genotype, disease staging, or the extent of cognitive deficit (**Fig. 1H-L**, **Suppl. Fig. 2**, and **Suppl. Table 7**). We found that retinal Cp significantly correlated with severity of brain NFTs (**Fig. 1H**; *r_s_* = 0.54, *p*<0.0001) and was 2.1-2.4-fold higher in patients with advanced Braak stages (**Fig. 1I**; *Stage III-IV or V-VI versus 0-II: p*<0.05-0.001) suggesting Cp’s involvement in brain tauopathy progression. Retinal Cp burden significantly and moderate-to-weakly correlated with the following brain pathologies: Aβ plaques (*r_s_* = 0.40, *p* = 0.0014), ABC severity score (*r_s_* = 0.54 *p*<0.0001), neuropil threads (NT; *r_s_* = 0.37, *p* = 0.0033), cerebral amyloid angiopathy (CAA; *r_s_* = 0.35, *p* = 0.0057), gliosis (*r_s_* = 0.40, *p*<0.0016), and brain atrophy (*r_s_* = 0.48, *p* = 0.0001; **Suppl. Table 7**), highlighting the possible links between retinal Cp infection and brain AD pathology. Notably, retinal Cp burden was higher in APOE ɛ4 allele carriers compared with non-carriers, regardless of AD diagnosis (**Fig. 1J**; *p* = 0.0373).

Common bacterial infections, such as *Helicobacter pylori*, Cp, *Borelia burgdorferi*, and spirochetal *Trepenoma* have been previously linked to cognitive decline and increased dementia risk in elderly^107^. Here, we found that individuals with elevated levels of retinal Cp inclusions exhibited lower Mini-Mental State Examination (MMSE) scores (**Suppl. Fig. 2D**; *r_s_* = -0.53, *p*<0.0001). Furthermore, retinal Cp burden significantly correlated with Clinical Dementia Rating (CDR) scores (**Fig. 1L**; *r_s_* = -0.43, *p* = 0.0010) and Montreal Cognitive Assessment (MoCA) scores (**Suppl. Fig. 2E**; *r_s_* = -0.56, *p* = 0.0334), reinforcing Cp’s link to cognitive impairment. Despite the small cohort size (n = 14-15), brain Cp burden strongly-to-very strongly correlated with increased brain AD pathology, including ABC score, Braak Stage, NFT, NT, gliosis, and atrophy scores (*r_s_* = 0.60-0.77, *p*<0.05-0.001), and reduced MMSE cognitive performance (**Suppl. Table 8**; *r_s_* = -0.73, *p* = 0.0043).

### 2. Dysregulated Cp interactome associates with activated NLRP3 inflammasome and cell death pathways in the AD brain and retina

The detection of Cp inclusions in the retinas and brains of AD patients prompted further investigation into Cp infection-induced protein dysregulation in these tissues, using a mass spectrometry-based proteomics in a separate human cohort (see **Suppl. Tables 2 - 5**), as previously described^54^. Metascape gene ontology (GO) analysis identified multiple dysregulated human proteins related to response to bacterial infection, including gram-negative intracellular bacteria, in the brains and retinas of AD patients (**Fig. 2A-B**), suggesting a significant involvement of bacterial infection in AD pathogenesis. To gain a closer look at *Chlamydia* infection, we searched for differentially expressed proteins (DEPs) in AD versus NC brains and retinas that were included in the *Chlamydia* interactome (**Fig. 2C-D**; extended data in **Suppl. Tables 9** and **10**). Out of 787 proteins in the *Chlamydia* interactome^92–96^, we identified 607 in the human brain, of which 52 DEPs were downregulated and 32 DEPs were upregulated (13.8% DEPs) in AD patients compared with NC controls (**Fig. 2C**). Importantly, despite being separate cohorts, similar bacterial infection-associated pathways (**Fig. 2A-B**) and dysregulated *Chlamydia* interactome DEPs were identified in the AD retina (**Fig. 2D; Suppl. Fig. 3A**), with 52 downregulated DEPs and 40 upregulated DEPs (13.0% DEPs) among the 710 identified (**Fig. 2D**). These data suggest shared infection-associated mechanisms in the brains and retinas of AD patients. GO network analysis further revealed the enrichment of proteins involved in immune responses to microorganisms and cell death in the AD brains and retinas (**Fig. 2E-F**; extended data in **Suppl. Fig. 3B-D** and **Suppl. Fig. 4A-C**). Inflammation-related proteins were primarily associated with cytokine signaling, toll-like receptor (TLR) pathways, interferon responses, NF-κB activation, NLRP3 inflammasome activation, and pyroptosis—pathways typically triggered by gram-negative bacteria in peripheral tissues^108,109^.

**Figure 2.**
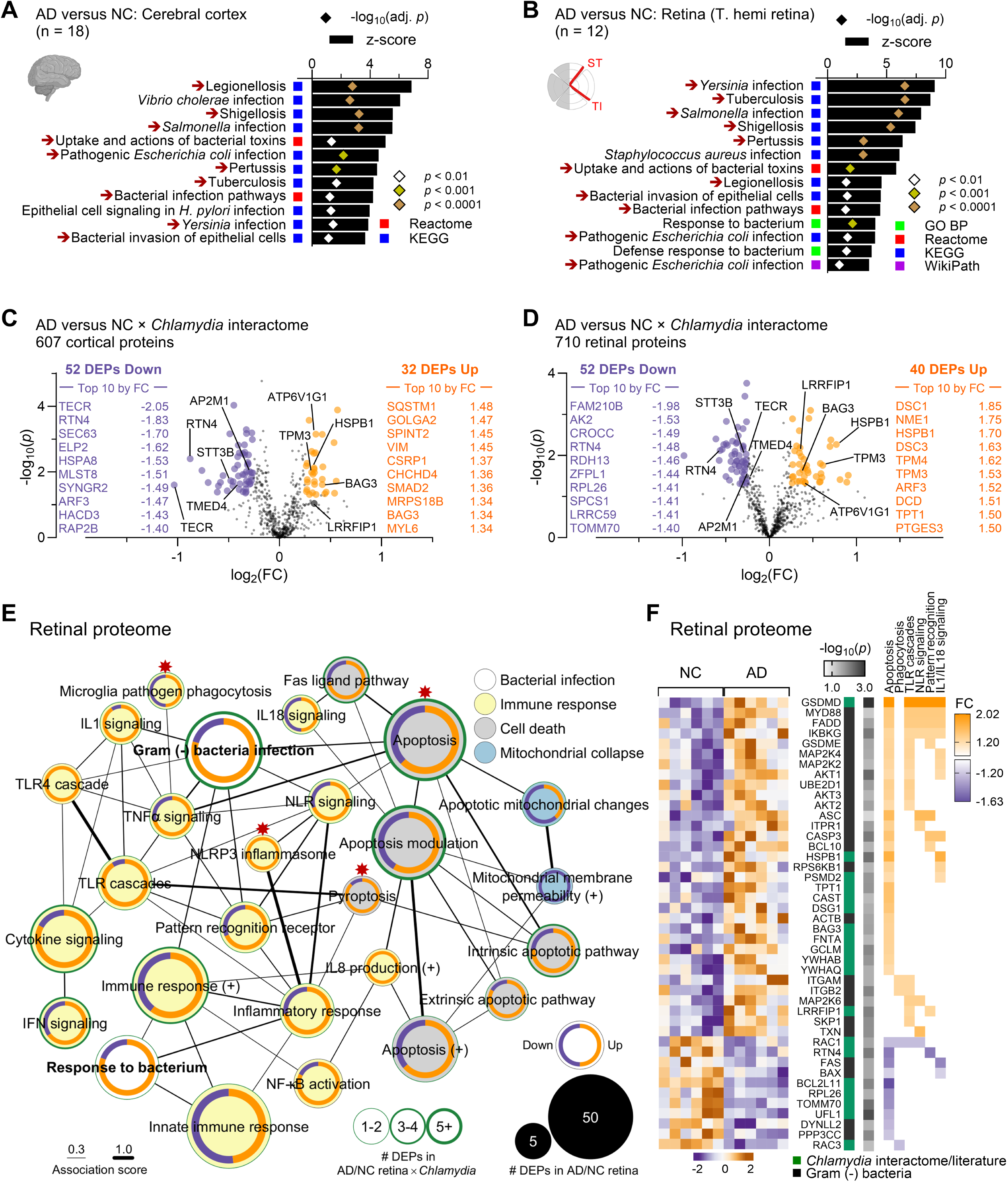
Bacterial infection-associated proteome pathways in AD retina and cerebral cortex. **A-B**. Gene ontology (GO) analysis of differentially expressed proteins (DEPs) related to bacterial infection (**A**) in the cerebral temporal cortex and (**B**) in the temporal hemi-retina from 2 separate cohorts of human donors with AD (brain: n = 10; retina: n = 6) versus NC (brain: n = 8; retina: n = 6) subjects. The analysis was carried out in Metascape and included the Reactome, Kyoto Encyclopedia of Genes and Genomes (KEGG) and WikiPathways (WikiPath) databases. Red arrows indicate the shared pathways between brain and retina. Bar and symbol graphs represent z-scores and Benjamini-Hochberg adjusted *p*-values from Metascape analysis, respectively. Range of *p*-values are presented as color-coded symbols. **C-D**. Volcano plots display the fold changes [log_2_(FC)] and significance level [-log_10_(*p*)] (**C**) in the cerebral cortex and (**D**) in the retina of AD versus NC subjects for proteins known to interact with *Chlamydia* inclusion membrane/vacuole. The list of human interacting proteins (termed ‘*Chlamydia* interactome’) was extracted from 4 original studies and a meta-analysis study and comprises 787 human proteins. Top 10 DEPs by FC upregulated (orange) and downregulated (purple) interactors are shown. The highlighted proteins (5 downregulated: AP2M1, RTN4, STT3B, TECR, TMED4; 5 upregulated: ATP6V1G1, BAG3, HSPB1, LRRFIP1, TPM3) were found in both the temporal cortex and the retina (F). **E**. GO network (Metascape) of enriched retinal pathways related to bacterial infection, immune response and cell death, including the apoptotic mitochondrial collapse. The size of the nodes represents the number of DEPs in AD versus NC retina, with the inner ring showing the proportion of these DEPs that are downregulated (purple) or upregulated (orange) in AD. The green border and its thickness represent the number of DEPs that interact with *Chlamydia* inclusion in each pathway. The thickness of edges between nodes represents the shared DEPs (association score) between pathways. Red asterisks indicate pathways that were further explored and validated. **F**. Heatmaps of upregulated (orange) and downregulated (purple) DEPs [-log_10_(*p*) and FC] in AD versus NC retina for selected pathways. Only proteins connected to gram-negative bacterial infection (Metascape analysis) and *Chlamydia* infection (*Chlamydia* interactome and literature) are shown for each pathway. The heatmap on the left corresponds to the protein expression level in the 6 NC individuals and the 6 AD patients, normalized by unit variance scaling and generated in ClustVis. Clustering of DEPs was carried out manually based on their involvement in selected pathways for visual clarity.

*Chlamydia* has been shown to trigger host’s innate immune response, requiring TLR2/MYD88 signaling and NLRP3/ASC/Caspase-1 inflammasome^28,29^. Indeed, both MYD88 innate immune signal transduction adaptor (MYD88) and PYD and CARD domain containing protein (PYCARD or ASC) were upregulated in the AD retina (**Fig. 2F**). Additionally, the DNA pathogen sensor and *Chlamydia* interactor, LRR binding FLII interacting protein 1 (LRRFIP1), which positively regulates TLR4 by competing with FLII actin remodeling protein (FLII) for interaction with MYD88^110^, was upregulated in both the AD brain and retina (**Fig. 2F; Suppl. Fig. 3A**). Importantly, the retinal AD proteome was enriched in proteins linked to pyroptosis (**Fig. 2E** and **Suppl. Fig. 3B, D**), a form of inflammatory regulated necrosis triggered by intracellular pathogens, including *Chlamydia*^111^. Notably, three members of the gasdermin (GSDM) family, GSDMD, GSDME (or DFNA5) and GSDMA, were upregulated in AD retina. Proteins involved in apoptosis, pyroptosis and inflammation were generally associated with levels of Aβ_1-42_ measured by ELISA in the retina, as well as retinal and cerebral isoforms of tau, quantified by MS (**Suppl. Fig. 4D-H** and **Suppl. Fig. 5**). Overall, these findings suggest the presence of intracellular gram-negative bacterial infection, specifically *Chlamydia*-related proteins, and associated inflammation and degeneration in the AD brains and retinas.

### 3. NLRP3 inflammasome activation in retinal cells of MCI and AD patients: links to Cp burden, cell death pathways, and brain atrophy

Cp activates the NLRP3 inflammasome in murine infection models^29,112,113^, however, its effect on NLRP3 activation in the AD retina remains unknown. To investigate the link between Cp, NLRP3 inflammasome, and neurodegeneration in the AD retina at different disease stages, we applied a quantitative immunohistochemistry analysis on retinal cross-sections from patients with MCI due to AD and AD dementia as compared with matched non-AD individuals with normal cognition (**Fig. 3**; extended data in **Suppl. Fig. 6** and **Suppl. Table 11**). Representative micrographs showed elevated retinal NLRP3 expression colocalized with caspase-1 along with more prominent Cp-associated ASC specks signal in the retina of MCI and AD patients compared with NC controls (**Fig. 3A, B** and **Suppl. Fig. 6A, B**). Retinal caspase-1 and ASC immunoreactive areas were 2.5- and 3.1-fold increased, respectively, only at the later stage of disease, in patients with AD dementia (**Fig. 3C, D**), but not in MCI patients. Notably, our quantitative analyses revealed that retinal NLRP3 expression was 2.1-fold increased early in the MCI retina, and furthermore, 3.6-folds in the AD retina, compared with NC controls (**Fig. 3E**, *p*<0.001-0.0001). The early rise in the NLRP3 immunoreactivity and later induction of caspase-1 and ASC markers may suggest that NLRP3 is activated by earlier processes such as misfolded Aβ and tau accumulation in the retina. Similarly, inflammatory cascades are initiated early in AD progression, as seen by increased micro- and macro-gliosis in the MCI retina^50,54,67^.

**Figure 3.**
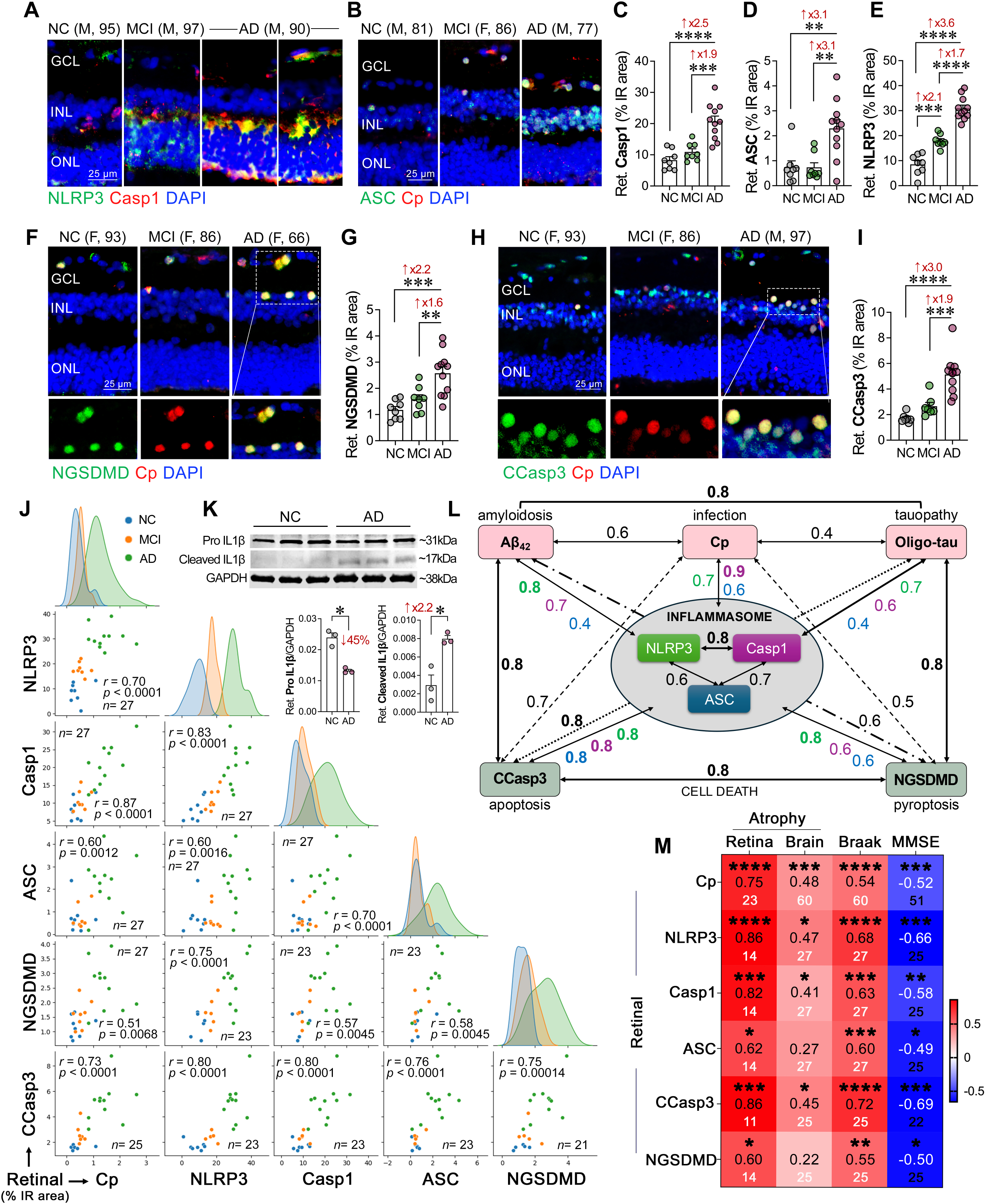
Retinal NLRP3 inflammasome, pyroptotic, and apoptotic markers and associations with Cp infection in early and advanced AD. **A-B**. Representative immunofluorescence images of retinal cross-sections from MCI and AD patients versus NC controls stained for NLRP3 inflammasome activation markers: **A**. NLRP3 (green), Caspase-1 (Casp1, red), and DAPI (nuclei, blue), and **B**. ASC (green), Cp inclusions (red), and DAPI (nuclei, blue); right panel, higher magnification images with separate channels showing ASC^+^ signals colocalized with Cp-infected cells within the INL (yellow). **C-E**. Quantitative IHC analysis of retinal, **C**. Casp1, **D**. ASC and **E**. NLRP3 percentage IR area in donors with NC (n = 8), MCI (due to AD; n = 8), and AD dementia (n = 11). **F**. Representative immunofluorescence image of retinal cross-sections from MCI and AD patients versus NC controls stained for the pyroptotic marker, N-terminal gasdermin D (NGSDMD, green), Cp inclusions (mAb; red), and DAPI (nuclei, blue). **F**. Lower image panel, separate channels showing NGSDMD^+^ cells infected with Cp located in the retinal INL and GCL (yellow). **G**. Quantitative analysis of retinal NGSDMD percentage IR area in donors with NC (n = 8), MCI (n = 8), and AD (n = 11). **H**. Representative immunofluorescence images of retinal cross-sections from MCI and AD patients versus NC controls stained for the apoptotic marker, cleaved caspase-3^+^ (CCasp3^+^, green), Cp inclusions (red), and DAPI (nuclei, blue). **H**. Lower image panel, high magnification images with separate channels showing CCasp3^+^ cells infected with Cp localized within retinal INL (yellow). **I**. Quantitative analysis of retinal CCasp3 percentage IR area in subset donors with NC (n = 7), MCI (n = 7), and AD (n = 11). Scale bars: 50 µm. **J**. Multivariable Pearson’s correlation coefficient (*r_p_*) analyses are presented by scatter plots and adjusted *p* values, to assess the relationships between the retinal markers, including Cp, NLRP3, Casp1, ASC, CCasp3—apoptosis, and NGSDMD— pyroptosis. Gaussian distribution curves for each marker are also presented. **K**. Representative Western blot analysis of retinal lysates from AD and NC subjects (n = 3) demonstrating the pro- and mature forms of IL-1β immunolabeling, with corresponding band intensity quantification normalized to GAPDH. **L**. Schematic representation depicting the strength (*r_P_*) of association between various retinal marker within three categories: 1. Inflammasome activators (Cp, Aβ_42_, and oligomeric tau), 2. NLRP3 inflammasome activation markers (NLRP3, Casp1, and ASC), and 3. Cell death (CCasp3 and NGSDMD). Pearson’s correlation *r_p_* values are highlighted in bold for very strong correlations (≥ 0.8), with associations to NLRP3, Casp1, and ASC indicated in green, purple, and blue, respectively, while other interactions are marked in black. **M**. Heatmap depicting pairwise Pearson’s correlations (*r_p_*) between retinal Cp-related markers and retinal atrophy, whereas pairwise Spearman’s rank correlations (*r_s_*) are between retinal Cp-related markers and brain atrophy, Braak stage, and MMSE score. Stars present the level of significance by unadjusted *p* values, values in middle row are *r_p_* or *r_s_*, and lower values are the sample sizes. Data from individual subjects (circles) and group means ± SEMs are shown. Fold changes are indicated in red. Statistical significance is denoted as \**p* < 0.05, \*\**p* < 0.01, \*\*\**p* < 0.001, and \*\*\*\**p* < 0.0001, determined by one-way ANOVA with Tukey’s post hoc multiple comparison test or by two-tailed paired unpaired Student’s t test for two-group comparison.

These findings together with our MS results prompted us to investigate the possible impact of retinal Cp infection and NLRP3 inflammasome activation on cellular apoptosis and pyroptosis (**Fig. 3F, H** and **Suppl. Fig. 6C, D**). We found that Cp-infected cells in the AD retina frequently coexpressed the pyroptotic marker cleaved gasdermin D at N-terminal (NGSDMD) and the apoptotic marker cleaved caspase-3 (CCasp3) (**Fig. 3F, H**; extended data in **Suppl. Fig. 6C, D**). Quantitative analyses indicated that the levels of both retinal NGSDMD pyroptotic and CCasp3 apoptotic markers were significantly 2.2- and 3.0-fold elevated, respectively, in AD retinas compared with NC retinas (**Fig. 3G, I**, *p*<0.001-0.0001), whereas the MCI retina had a trend of increase that did not reach statistical significance. Indeed, most Cp-infected cells exhibited positive staining for either pyroptotic or apoptotic cell markers, suggesting that Cp infection in the AD retina may trigger cell death via these cellular pathways. Furthermore, the increased levels of NGSDMD in the AD retina provided evidence for NLRP3 inflammasome activation.

Next, we explored the inter-relationships between retinal Cp burden, NLRP3 inflammasome components, and degeneration markers (**Fig. 3J-K**). Multivariate correlations analysis indicated that retinal Cp burden had a very strong to strong associations with retinal NLRP3 inflammasome activation components, in particular, caspase-1 (*r* = 0.87, *p*<0.0001), as well as NLRP3 (*r* = 0.70, *p*<0.0001) and ASC (*r* = 0.60, *p* = 0.0012) (**Fig. 3J**; extended data in **Suppl. Table 11**). As compared with the NC retina, the AD retina exhibited activation of NLRP3 inflammasome as further revealed by increased expression of active cleaved interleukin-1β (IL-1β) levels (**Fig. 3K**). Indeed, the pro-form of IL-1β was significantly reduced, whereas the mature form was markedly elevated in AD versus NC retinas (**Fig. 3K**). In addition, Pearson’s correlation (*r_p_*) analyses revealed a strong association between retinal Cp and retinal Aβ_42_ burden (*r* = 0.63, *p*<0.0001), and a moderate association with retinal T22^+^ oligo-tau burden (*r* = 0.43, *p*<0.0040; **Suppl. Table 7**), both of which were very strongly to strongly linked to NLRP3 inflammasome marker (*r* = 0.70-0.81, *p*<0.001-0.0001, **Fig. 3L** and **Suppl. Table 11**). In our cohort, retinal oligo-tau was equally and strongly correlated with retinal CCasp3 apoptosis (*r* = 0.80, *p*<0.0001) or NGSDMD pyroptosis (*r* = 0.77, *p*<0.0001). However, retinal Aβ_42_ had a stronger correlation with CCasp3 apoptosis (*r* = 0.77, *p* = 0.0003) compared with NGSDMD pyroptosis (*r* = 0.64, *p* = 0.0247). Notably, all three retinal NLRP3 inflammasome components were very significantly and strongly inter-correlated (**Fig. 3J, L**; *r* = 0.60-0.83, *p* = 0.0016-*p*<0.0001) with NLRP3 and Casp1 most closely linked (**Fig. 3L**). As it relates to associations between retinal NLRP3 inflammasome components with retinal cell death markers, all three components were strongly to very strongly correlated with CCasp3 apoptosis (**Fig. 3J, L**; *r* = 0.76-0.80, *p*<0.0001). At the same time, NLRP3 expression was most closely associated with NGSDMD pyroptosis (*r* = 0.74, *p*<0.0001) as compared with Casp1 and ASC that had moderate correlations with NGSDMD (*r* = 0.57-0.58, *p*<0.01). These data suggest that Aβ_42_ species, more than the tau oligomeric forms, interact closely with Cp to trigger NLRP3 and caspase-1 inflammasome activation in the retina, which may mediate cellular apoptosis and pyroptosis contributing to retinal degeneration in AD.

We found that retinal NLRP3 and Casp1 were very strongly correlated with retinal atrophy, an index of retinal thickness from inner limiting membrane to outer limiting membrane (**Fig. 3M**; *r* = 0.82-0.86, *p*<0.001-0.0001), whereas these markers had moderate correlation with brain atrophy (**Fig. 3M**; *r* = 0.41-0.47, *p*<0.05). Retinal ASC showed strong correlation with retinal atrophy (*r* = 0.62, *p* = 0.0171) but not with the brain atrophy. Retinal Cp burden has strong association with retinal atrophy, and it was moderately but most significantly correlated with brain atrophy (**Fig. 3M**; *r* = 0.48-0.75, *p*<0.001-0.0001). Retinal CCasp3 showed moderate to very strong correlation with brain and retinal atrophy (**Fig. 3M**; *r* = 0.45-0.86, *p*<0.05-0.001), whereas retinal NGSDMD had strong correlation with retinal atrophy (**Fig. 3M**; *r* = 0.60, *p* = 0.0247) but not with brain atrophy. All retinal inflammasome components, Cp, and cell death markers, moderately to strongly correlated with the Braak stages (*r* = 0.55-0.72, *p*<0.01-0.0001), and inversely correlated with the MMSE cognitive performance scores (**Fig. 3M**; *r* = -0.49-[-0.69], *p*<0.05-0.0001). These data suggest that Cp and related NLRP3 inflammasome activation can drive retinal degeneration by promoting apoptotic (CCasp3) and pyroptotic (NGSDMD) cell death pathways (**Fig. 3L, M**), and eventually lead to retinal atrophy that was parallel to the extent of brain atrophy. Furthermore, the correlations between retinal Cp burden, NLRP3 components, and cell death markers with the severity of brain AD pathology, Braak stage, and MMSE cognitive scores (**Suppl. Table 12**), suggest that Cp infection-mediated NLRP3 inflammasome activation and neurodegeneration may play a role in AD progression and associated cognitive decline.

### 4. Retinal gliosis closely localizes to Cp-infected cells, with microglia exhibiting impaired Cp phagocytosis in AD patients

Cp is known to infect and extensively proliferate in astrocytes and neurons, whereas microglia are involved in Cp phagocytosis^114^. Previous studies, including our own, have demonstrated increases in glial cell activation markers in the retinas of individuals with MCI and AD^50,54,67^. However, the potential interplay between Cp and glial cells in the MCI and AD retina remains unexplored. In the current study, we observed significant spatial interactions between Cp and glial cells, specifically microglia and astrocytes, as well as the Müller glia, in the AD retina (**Fig. 4A-K, Suppl. Fig. 7A, B**). We detected elevated levels of retinal GFAP^+^ (astrocyte and reactive Müller glia marker) and vimentin^+^ (Müller glia marker) macroglia, as well as IBA1^+^ microglia in both MCI and AD retinas compared with NC controls (**Fig. 4B, D, G, Suppl. Fig. 7C-E**). Furthermore, we identified moderate to strong associations between Cp burden and these gliosis markers (**Fig. 4C, E, H**). Strong correlations between retinal Cp and gliosis were noted for GFAP^+^ astrocytosis and IBA1^+^ microgliosis (**Fig. 4C** and **4H**; *r* = 0.65-0.70, *p*<0.0001). However, only a moderate association was noted with vimentin^+^ macroglia (**Fig. 4E**; *r* = 0.55, *p* = 0.0090) that did not appear to be expressing the activation-related marker GFAP. Similarly, brain Cp burden also showed a strong correlation with brain gliosis (**Suppl. Table 8;** *r* = 0.77, *p* = 0.0008). The strong association between Cp burden and gliosis indicates potential inflammatory cascades following Cp infection. Furthermore, we found that retinal astrocytes and microglia were involved in engulfing Cp-infected cells (**Fig. 4A** and **4I**). Whereas retinal astrocytes appeared to engulf Cp-infected cells in the innermost retinal layers, retinal microglia appeared to phagocytose the Cp/Cp-infected cells (**Fig. 4A, I**). Specifically, retinal microglia appeared to exhibit different stages or types of responses to Cp-infected cells, whereas most cells were close to and in partial contact with the Cp positive cells. Other microglia were directly involved in engulfing or ingesting Cp-infected cells (**Fig. 4I**; extended data on microglia recognizing, engulfing or phagocytosing Cp-infected cells in **Suppl. Fig. 8A-C**). The percent of these Cp-associated microglia (CAM) was increased by 60% (**Fig. 4J** and **Suppl. Fig. 8A-C**), however, 62% fewer retinal CAM relative to Cp burden were detected in the AD versus NC retinas (**Fig. 4K**). These findings show relatively lower microglial cells in proximity or engulfing Cp-infected cells in AD patients, suggesting impaired microglial responses to cells harboring bacterial inclusions. Multi-interaction analyses between retinal gliosis and various AD biomarkers in the retina and brain (**Fig. 4L**, heatmap; extended data in **Suppl. Tables 11** and **12**), revealed very strong associations between retinal GFAP^+^ or vimentin^+^ macrogliosis and NLRP3 load (*r* = 0.84-0.91, *p*<0.01-0.0001), and between retinal GFAP^+^ astrogliosis and CCasp3^+^ apoptosis (*r* = 0.85, *p*<0.0001). As it relates to amyloidosis and tauopathy, retinal Aβ_42_ and oligo-tau burdens most closely correlated with retinal IBA1^+^ microgliosis levels (**Fig. 4L**; *r* = 0.69-0.85, *p*<0.0001). In addition, retinal GFAP^+^ strongly predicted the Braak scores (*r* = 0.78, *p*<0.0001). These finding suggest close interactions between retinal glial cells and Cp-infected cells, strongly correlating with NLRP3 inflammasome components and apoptosis/pyroptosis cell death markers, and potentially impaired ability of microglia to phagocytose and clear Cp infection in the AD retina.

**Figure 4.**
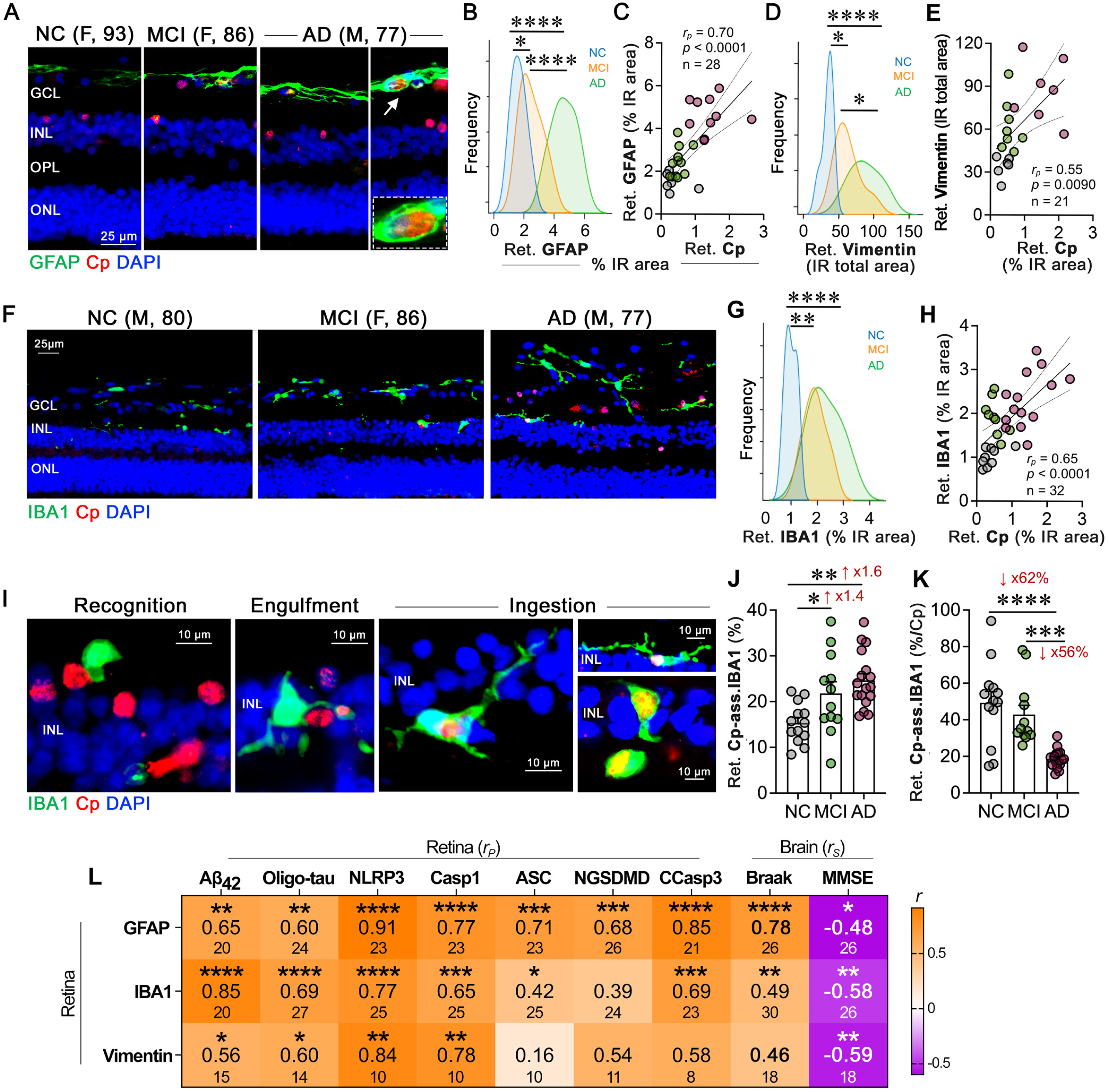
Cp-associated glial activation and phagocytosis in MCI and AD retina. **A.** Representative immunofluorescence images of retinal cross-sections from MCI and AD patients versus NC controls stained with the marker of macrogliosis (GFAP, green), Cp (mAb, red), and nuclei (DAPI, blue). High magnification images of Cp-infected retinal astrocytes at the NFL/GCL are shown in the right panel (Cp inclusions are engulfed by retinal GFAP^+^ astroglia). **B.** Gaussian distribution curves display quantitative analysis of retinal GFAP-immunoreactivity (IR) % area (frequency) in donors with NC (n = 8), MCI (due to AD; n = 10), and AD dementia (n = 10). **C**. Pearson’s correlation (*r_p_*) analysis between retinal Cp and GFAP % IR area in the same cohort. **D**. Gaussian distribution curves display the quantitative analysis of retinal vimentin (total IR area) in donors with NC (n = 6), MCI (n = 8), and AD (n = 7). **E**. Pearson’s correlation (*r_p_*) analysis between retinal Cp (% IR area) and vimentin (total IR area) in the same cohort. **F**. Representative immunofluorescence images of retinal cross-sections from MCI and AD patients versus NC controls stained with the microglial marker (IBA1, green), Cp (red), and nuclei (DAPI, blue). **G**. Gaussian distribution curves display quantitative analysis of retinal IBA1 % IR area in donors with NC (n = 9), MCI (n = 9), and AD (n = 14). **H**. Pearson correlation (*r_p_*) analysis between retinal Cp and IBA1 % IR area in the same cohort. **I**. Immunofluorescence images of retinal cross-sections show three distinct stages of microglial (green) involvement in the phagocytosis of Cp-infected cells (red): recognition (left image, microglial attached to Cp-infected cells), engulfment (second image, microglia cell processes surrounding Cp-infected cells), and ingestion (final three images, microglial internalization of Cp-infected cells). **J**. Quantitative analysis of Cp-associated IBA1-positive cells involved in recognition, engulfment, and ingestion of Cp-infected cells in donors with NC (n = 13), MCI (due to AD; n = 12), and AD dementia (n = 17). **K**. Quantitative analysis of retinal Cp-associated IBA1-positive cells per Cp burden (% IR area). **L**. Heatmap depicting Pearson’s correlations (*r_p_*) between retinal gliosis (GFAP- or vimentin-astrogliosis and IBA1-microgliosis) and retinal amyloidosis (Aβ_42_), tauopathy (oligo-tau), NLRP3 inflammasome activation (NLRP3, Casp1, ASC), and cell death (NGSDMD-pyroptosis, CCasp3-apoptosis). Spearman’s rank correlations (*r_s_*) were performed to assess associations between retinal gliosis, Braak stage (brain), and MMSE score (cognition). Stars present the level of significance by unadjusted *p* values, values in middle row are *r_p_* or *r_s_*, and lower values are the sample sizes. Data from individual subjects (circles) as well as group means ± SEMs are shown. Fold or % changes are shown in red. \**p* < 0.05, \*\**p* < 0.01, \*\*\**p* < 0.001, and \*\*\*\**p* < 0.0001, by one-way ANOVA and Tukey’s post hoc multiple comparison test.

### 5. Retinal Cp and NLRP3 prediction of disease diagnosis and status

We next aimed to assess whether retinal Cp burden could serve as a predictor of AD diagnosis, severity of brain pathology, disease stage, and/or cognitive dysfunction (**Fig. 5**; extended data in **Suppl. Figs. 9-11**). In addition to retinal Cp, we included key retinal markers that demonstrated significant correlations with brain pathologies and were associated with Cp infection, specifically NLRP3, Aβ_42_, and CCasp3-apoptosis. These markers were analyzed either in isolation or in combination with retinal gliosis (IBA1+GFAP+vimentin), atrophy, and Aβ_42_, which can be potentially imaged in living patients^46,48,49,53,60,65,115–120^. Multivariable analysis employing random forest models indicated that retinal Cp alone weakly predicted AD-related pathologies in the brain, including ABC score and Braak stage (**Fig. 5A, B**), as well as the cognitive function (MOCA, **Suppl. Fig. 9B**). Multiple models were fit using 5×2 cross validation to obtain distributions of model performance. However, the predictive power of retinal Cp was generally enhanced when combined with retinal Aβ_42_ or gliosis. We found that retinal Aβ_42_ alone was a good predictor of brain NFTs, ABC score, Braak stage, and the MMSE score (**Fig. 5A, B, D** and **Suppl. Fig. 9A**). Notably, the combined retinal Cp and gliosis index was the best predictor of ABC score (**Fig. 5A**, r^2^ = 0.34) and brain gliosis (**Fig. 5C**, r^2^ = 0.26). The best predictor of Braak stage was retinal Aβ_42_ combined with CCasp3 (r^2^ = 0.41), and also with NLRP3 (r^2^ = 0.38), and Cp (r^2^ = 0.28) (**Fig. 5B**). In addition, the combined retinal Aβ_42_ and NLRP3 index was the best predictor of MMSE score (**Fig. 5D**, r^2^ = 0.25), and retinal Aβ_42_ with either Cp or CCasp3 also predicted MMSE (**Fig. 5D**, r^2^ = 0.22-0.23). Retinal gliosis provided the most accurate prediction of brain gliosis (**Fig. 5C**), while no individual marker was predictive of brain atrophy (**Suppl. Fig. 9D**).

**Figure 5.**
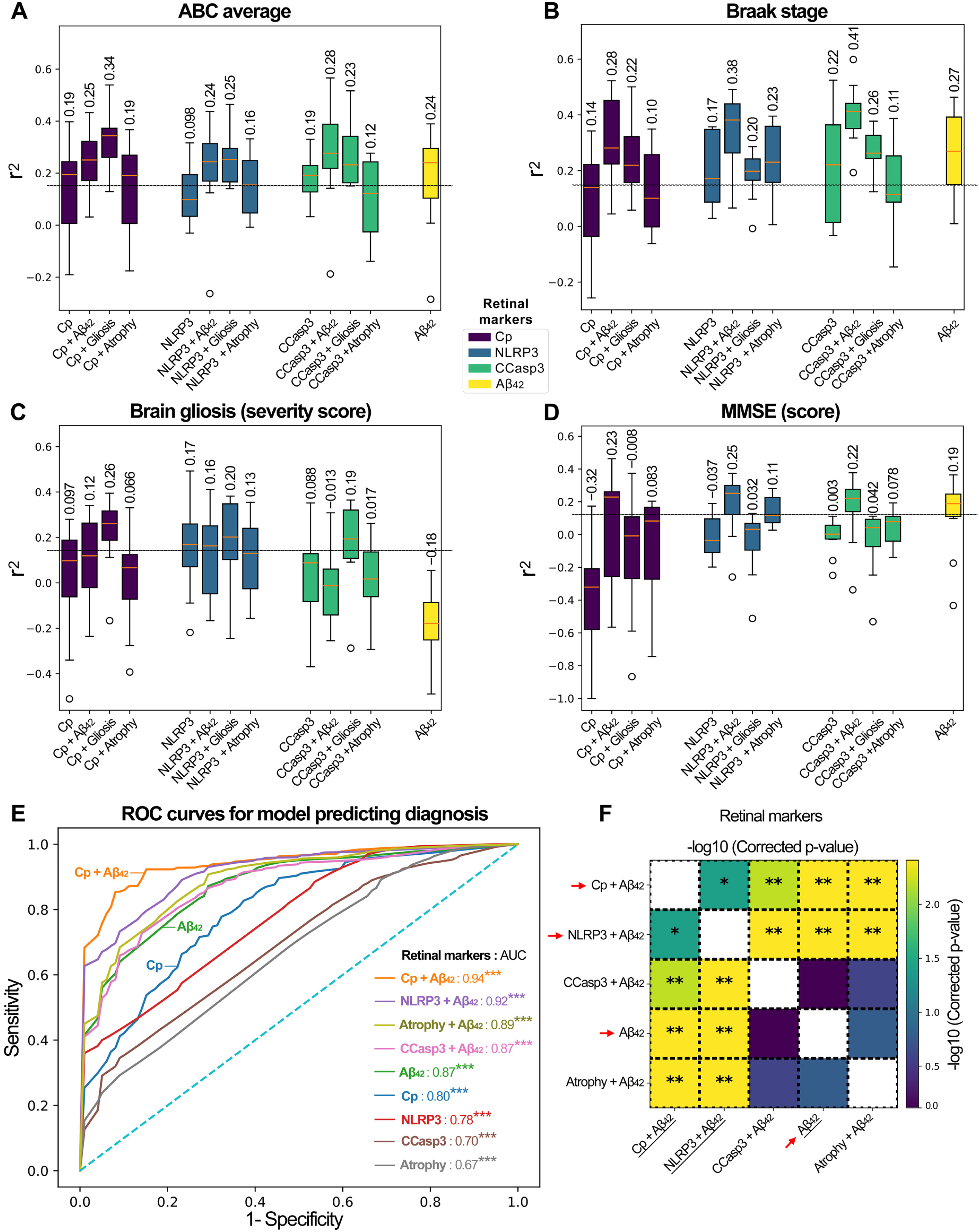
Multivariable predictions of brain AD pathology and cognitive dysfunction conferred by retinal Cp, NLRP3, CCasp-3, and Aβ_42_ markers. Random forest regressor using 80 estimators was trained on the data to predict several brain pathologies, including **A**. ABC average, **B**. Braak stage, **C**. brain gliosis, and **D**. mini-mental state examination (MMSE) score. The distributions show the spread of models trained on different folds of the 5×2 cross-validation. Only models performing with a variance coefficient of determination r^2^>0.15 (gray dotted line) were retained. **E**. The ROC curves for different retinal biomarkers, including Cp, Aβ_42_, NLRP3, CCasp3, and retinal atrophy either individual or combined with retinal Aβ_42_. Each model was obtained by averaging the curves across diagnosis separately in each cross-validation fold. In the ROC curves plot, AUC is listed for each curve and unadjusted *p* values are included (\*\*\**p* < 0.001). **F**. The models were compared by using a Wilcoxon signed-rank test and *p* values were adjusted for multiple comparisons using Benjamini-Hochberg correction. The heat map shows that among the top 5 performing models, we have 3 that are different from one another. The model trained on retinal Cp + retinal Aβ_42_ performed best and was significantly different from the second-best model (retinal NLRP3 + retinal Aβ_42_) with *p*<0.05. The other three models were different from the top 2, but not from one another. Red arrows highlight retinal markers, individually or in combination, which were significantly different among the performance models to predict disease diagnosis. Statistics: \**p* < 0.05 and \*\**p* < 0.01, adjusted for multiple comparisons with Benjamini-Hochberg procedure.

We further evaluated the performance of these variables using the area under the ROC curve (AUC) for disease diagnosis (**Fig. 5E** and **Suppl. Fig. 10**). When combined with retinal amyloidopathy (Aβ_42_), the AUC for retinal Cp increased from 0.80 to 0.94 (**Fig. 5E**), indicating that retinal Cp, in combination with retinal Aβ_42_, serves as an excellent marker for identifying disease status. Interestingly, all tested variables exhibited strong AUC values for disease diagnosis, with a notable enhancement in AUC values when combined with retinal Aβ_42_ (**Fig. 5E, F**). For the different diagnosis of the NC, MCI, and AD groups, retinal Aβ_42_ alone demonstrated superior AUC values compared with the other retinal markers (Cp, NLRP3, and CCasp3) alone, whereas retinal atrophy in combination with retinal Aβ_42_ showed the highest mean AUC for diagnosing AD status (**Suppl. Fig. 11A-D**). Based on the results presented in **Fig. 5E** and **5F**, we selected the model using Cp and retinal Aβ_42_ for evaluation on the test set; the results are reported in **Table 2**. The model performs poorly for subjects with MCI, but performs reasonably well for identifying NC and AD. We note that NC and MCI subjects had fewer subjects in the test set.

**Table 2.** Model performance on the machine learning test set.

|  | Precision | Recall | F1-Score | Support |
| --- | --- | --- | --- | --- |
| AD | 0.83 | 0.71 | 0.77 | 7 |
| MCI | 0.50 | 0.33 | 0.40 | 3 |
| NC | 0.67 | 1.00 | 0.80 | 4 |
| Accuracy |  |  | 0.71 | 14 |
| Macro average | 0.67 | 0.68 | 0.66 | 14 |
| Weighted average | 0.71 | 0.71 | 0.70 | 14 |

The final test set consisting of 14 samples with a distribution of diagnoses matching the training set was used to evaluate the random forest model using Cp and retinal A $\beta_{42}$ to predict AD diagnosis.

## Discussion

This study identifies Cp inclusions in the human retina and provides evidence linking Cp infection to retinal inflammation and neurodegeneration in AD. We found that Cp inclusions are substantially more prevalent in the retinas of AD patients than in those with MCI due-to-AD or NC controls. Additionally, a greater Cp burden was observed in APOE ɛ4 allele carriers, and this burden correlated with brain Cp load, AD pathological changes (e.g., brain NFTs, Braak, and ABC severity scores), and cognitive deficits (e.g., CDR, MMSE). Concomitant with retinal Cp burden there is an early and progressive increase in NLRP3 in the retina of MCI and AD patients, with very strong correlations with caspase-1. Retinal NLRP3 inflammasome components (NLRP3, ASC, and caspase-1) and downstream effectors of activation (mature IL-1β and N-terminal GSDMD) were strongly associated with retinal neuronal cell death via pyroptosis and apoptosis and correlated with gliosis and atrophy in both the retina and brain. Notably, we detected retinal microglial dysfunction in response to Cp infection in AD patients, which implies impaired clearance ability of Cp-infected cells. Our MS-based proteome analysis of AD brains and retinas further supports these findings, identifying dysregulated *Chlamydia* interactome proteins alongside dysregulated proteins involved in related immune response pathways, including response to gram-negative bacterial infection, NLRP3 inflammasome components, pyroptosis, apoptosis, and microglial pathogen phagocytosis. Finally, machine learning analysis demonstrated that combined indices of retinal Cp or NLRP3 with Aβ_42_ could be strong predictors of AD diagnosis (AUC = 0.92–0.94, p<0.001), as well as brain pathology and cognitive decline. These results support the development of noninvasive imaging techniques that combine retinal amyloid with infection/inflammasome markers for AD diagnosis. Moreover, the findings emphasize the role of Cp infection and its interactions with Aβ and the NLRP3 inflammasome in AD pathogenesis, reinforcing similar Cp involvement in the AD brain and retina.

This study reveals the presence and elevated levels of retinal and brain Cp inclusions in patients with AD dementia, with a 2.9- and 4.1-fold increase in Cp burden in the retinas and brains, respectively, compared with NC controls, with a strong correlation between these two CNS tissues. These findings align with several previous studies comparing Cp levels in body fluids or brains of AD patients versus healthy controls^15–18,20,121–125^. Indeed, various pathogens such as Cp infection have been implicated in the progression of neurodegenerative diseases including AD, by triggering chronic inflammation that results in neuronal damage and cognitive decline^12,15,17–19,121^. Cp infection is common in humans, with antibody prevalence in peripheral blood reaching 50% by age 20 and 80% by age 60-70 years, indicating lifelong asymptomatic exposure and reinfection^126^. Indeed, under immune and antibiotic pressure, Cp can form aberrant bodies that can persist for years, potentially driving chronic inflammation^127,128^. Cp is distinct from other *Chlamydia* species, particularly *C. trachomatis*, which is the leading cause of infectious blindness and the most common sexually transmitted urogenital infection^129^. Both species can cause chronic, low-grade infections, suggesting a broader link between *Chlamydia* and chronic inflammation, including in CNS disorders. Epidemiological studies show a strong association between Cp infection and AD, with a five-fold increased risk of AD in the presence of Cp^19^. Cp is thought to enter the CNS via the nasal and intravascular routes through monocytes, with evidence including Cp DNA in the olfactory bulb of AD patients and accelerated Aβ plaque development in Cp-inoculated mice^23,25^. Given the retina’s direct anatomical connection to the brain and the parallel development of AD pathology in both tissues^44–76^, the presence of Cp infection in the AD retina is therefore not unexpected. These findings highlight the parallel susceptibility of the AD brain and retina to Cp infection and underscore its potential role in exacerbating AD pathology and cognitive decline, which merits future investigation.

Our findings demonstrate retinal and brain Cp inclusions in 60-79% of early AD (MCI) patients and 100% of AD dementia patients as compared with 38-40% of age- and sex-matched NC individuals, while not reaching significance for prodromal AD (MCI). In fact, cerebral and retinal Cp inclusions were detected in relatively large amount of our aged population without showing any symptoms. These results suggest that Cp may not be an early driver of AD pathology nor essential for cognitive decline, but rather a consequence of AD-related processes that can further exacerbate cognitive deterioration. It is conceivable that increased Cp burden in retinas and brains of AD patients is an outcome of the blood-brain barrier and/or blood-retinal barrier breakdown in AD^40,80,130,131^, and excessive infiltration of pathogens into the CNS, particularly at the dementia phase. However, given that both cerebral and retinal vascular changes occur early in AD progression^64,76,132–138^, it remains unclear why Cp load is not significantly elevated in MCI patients. It is also possible that Cp may infect the CNS through alternative routes, such as the nasal route^23^, rather than exclusively via the vascular pathway. Furthermore, these results perhaps reflect a later dysfunction in microglial capacity to phagocytose and clear Cp/Cp-infected cells in patients with AD dementia, thereby contributing to increased Cp burden in these patients. Hence, Cp infection may contribute to neurodegeneration by exacerbating inflammatory cascades, alongside pathogenic Aβ and tau forms, through glial activation and NLRP3 inflammasome signaling. Even at the MCI stage, this could potentially drive cognitive decline, progressing from mild impairment to dementia. Future studies are warranted to investigate whether Cp is a consequence of AD pathology or a trigger that drives or contributes to dementia.

The current study demonstrates strong associations between retinal Cp burden and retinal amyloidogenic hallmarks of AD, specifically Aβ_42_ and Aβ_40_ alloforms, with moderate to lack of associations to retinal tauopathy isoforms such as PHF-tau, CitR_209_-tau, tau oligomers, pS396-tau, AT8^+^ p-tau, and MC1^+^ tangles. These findings suggest that the presence of retinal Cp may induce the secretion of Aβ, which could function as an antimicrobial peptide^139,140^, thereby enhancing the immune response against pathogens in the CNS. This interpretation aligns with a potential hypothesis that Aβ, often associated with AD, might play a protective role in the brain, especially in response to microbial threats^139–141^. Interestingly, both retinal and brain Cp presence moderately to strongly correlated with the severity of brain disease stage (e.g., NFT, ABC, and Braak stage severity score), implicating Cp infection in contributing to brain tauopathy and AD progression. Furthermore, retinal Cp burden was elevated in individuals carrying the APOE ɛ4 allele, highlighting that genetic predisposition that impacts cellular lipid content^142,143^, which is required for Cp’s intracellular growth and proliferation^144–146^, may modulate susceptibility to Cp infection in the CNS. Future studies should establish the connection between APOE allele genotype, Cp infection in the CNS and risk for AD development and progression.

Cp infection, among other gram-negative bacterial infections, activates the NLRP3 inflammasome via LPS, driving inflammatory cascades and cellular degeneration^147^. Retinal Cp burden strongly correlated with NLRP3 inflammasome components—NLRP3, caspase-1, and ASC—and inflammasome activation occurred early in disease progression, as evidenced by significantly increased NLRP3 immunoreactivity in MCI patients compared with controls. However, caspase-1 and ASC were activated only at later stages of AD, suggesting that while NLRP3 is primed early in the disease, potentially through early abnormal Aβ and tau accumulation^31,148^, full inflammasome activation, requiring a secondary signal, occurs later during AD progression. Our findings of increased mature IL-1β form in the AD retina further support the active status of retinal NLRP3 inflammasome in AD. Indeed, NLRP3 inflammasome activation requires a two-step process involving primming (signal 1) and activation (signal 2)^149,150^. Priming is triggered by inflammatory stimuli, such as those recognized by TLRs or cytokines like TNF-α, leading to NF-κB activation and subsequent upregulation of NLRP3 and mature IL-1β. Signal 2 involves various pathogen-associated molecular patterns (PAMPs) and damage-associated molecular patterns (DAMPs) that promote NLRP3 inflammasome assembly, leading to caspase-1 activation and the cleavage of pro-IL-1β and pro-IL-18 to their respective mature forms (reviewed in^149^). Cp can act as secondary signal to fully activate the NLRP3 inflammasome that further potentiate inflammation and neurodegeneration in the AD retina, a hypothesis that warrants further investigation.

Our proteomic analyses further supported our histological findings and identified dysregulated pathways associated with the immune response to intracellular obligatory, gram-negative bacterial infection, TLR and NLRP3 inflammasome activation, and cell death in both the retina and brain. Specifically in the AD retina, Cp infection triggered activation of the NLRP3 inflammasome and cell death pathways such as pyroptosis (GSDMA, GSDMD, GSDME) and apoptosis (CASP3, FADD), which are hallmarks of neurodegenerative diseases^151–153^. Elevated NLRP3 inflammasome components in AD retinas correlated with retinal Aβ_42_ and oligo-tau levels, closely linked to degeneration markers and retinal atrophy, suggesting that Cp infection induces a localized inflammatory response in the retina, mirroring inflammatory processes in the brain that contribute to neurodegeneration^13,154–158^. Our mass spectrometry analyses further revealed a shared *Chlamydia* interactome between the retina and brain, despite analyzing different cohorts. Out of the 787 proteins that constitute the *Chlamydia* interactome, ∼13% proteins were dysregulated in the AD retina and brain, with ten proteins were similarly altered in both tissues, including reticulon 4 (RTN4), TECR, STT3B, TMED4 and AP2M1, which were downregulated, and HSPB1, TPM3, BAG3, LRRFIP1 and ATP6V1G1, which were upregulated. Although the precise role of these *Chlamydia* interactome proteins in AD pathology remains unclear, accumulating evidence indicates that these proteins are involved in biological processes relevant to AD, including regulating apoptosis and mitochondrial function (e.g., RTN4, HSPB1, BAG3), modulating inflammatory and synaptic plasticity pathways, endoplasmic reticulum stress (TECR), and clearance of Aβ (LRRFIP1, HSPB1) and tau (BAG3)^110,111,159–165^. Notably, the DNA pathogen sensor and *Chlamydia* interactor LRRFIP1, which positively regulates TLR4 by competing with FLII actin remodeling protein (FLII) for interacting with MYD88^110^, was upregulated in the AD brain and retina. Despite these observations, the functional significance of these interactors in the context of infection-driven inflammation and neurodegeneration in AD progression remains largely unexplored. Therefore, comprehensive studies are warranted to elucidate the mechanistic links of these interactome proteins with infection, inflammatory cascades, and neurodegenerative processes during AD progression.

This study also highlights the complex interactions between Cp-infected retinal cells and glial cells, particularly astrocytes and microglia. Increased gliosis was observed in AD retinas, as previously reported by our group and others^50,54,67^, with strong correlations between Cp load and gliosis markers (GFAP, IBA1). Notably, we found that 62% smaller population of retinal microglia were engaged in engulfing or ingesting Cp-infected retinal cells in AD patients compared with NC individuals, suggesting impaired ability to recognize, bind, and phagocytose Cp-infected cells by retinal microglia in AD and a dysfunctional immune response that could exacerbate chronic infection and inflammasome activation, leading to neurodegeneration. These findings encourage further study of microglial phenotypes and dysfunction in AD and its potential impact on pathogen and misfolded protein clearance, chronic inflammation and progressive neurodegeneration.

Despite these comprehensive findings, several limitations should be considered. First, the smaller sample size of the brain tissues and lack of MCI group for the proteomic MS analysis, limits the ability to fully generalize the results. Larger cohorts would help validate the associations observed between Cp infection and neurodegenerative pathology. Second, the cross-sectional nature of the study precludes direct causal inference. Mechanistic studies in animal models and longitudinal studies would be valuable in confirming the causational or temporal relationship between Cp infection and AD progression. Third, the lack of information on patients’ visual function, which might be crucial for determining whether Cp infection contributes to visual abnormalities beyond cognitive dysfunction, and needs to be addressed in future studies. Additionally, while our proteomic analyses indicate a dysregulation in immune response to infection pathways, further functional studies are needed to investigate how Cp infection specifically might directly trigger these pathways and contribute to retinal and brain neurodegeneration and cognitive decline.

## Conclusions

This is the first study to document the presence, distribution, and severity of Cp infection and its association with NLRP3 inflammasome activation in the retinas and corresponding brains of MCI and AD patients compared with age- and sex-matched control individuals, offering new insights into the mechanisms linking peripheral infections to central neurodegeneration in AD. Our findings suggest that Cp infection is a significant factor in retinal degeneration and may play an important role in the pathogenesis of AD. The correlation between retinal Cp burden, NLRP3 inflammasome activation, retinal amyloidogenic peptides, brain tauopathy, and disease status underscores the potential for targeting Cp and its associated inflammatory pathways as a therapeutic strategy in AD. The interactions between Cp infection and glial cells, particularly the impaired microglial response, highlight the importance of maintaining proper immune function in preventing disease progression. Future studies should explore therapeutic interventions such as antibiotics aimed at controlling retinal and brain Cp infection, attenuating NLRP3 inflammasome activity, or enhancing microglial phagocytosis to prevent or mitigate AD-related neurodegeneration.

## Abbreviations

AD: Alzheimer’s disease
ADRC: AD research center
ANOVA: analysis of variance
APOE: apolipoprotein
ARG1: arginase-1
ASC: apoptosis-associated speck-like protein
Aβ: amyloid beta
BCA: bicinchoninic acid
C: central or Caucasian
CAA: Cerebral amyloid angiopathy
Casp-1: caspase-1
CCasp3: cleaved caspase-3
CDR: clinical dementia rating
CERAD: Consortium to Establish a Registry for Alzheimer’s Disease
Cit tau: citrullinated tau
CNS: central nervous system
Cp: Chlamydia pneumoniae
CSF: cerebrospinal fluid
DAB: 3,3’-diaminodbenzidine
DAPI: 4’,6-diamidino-2-phenylindole
DEPs: differentially expressed proteins
DMEM: Dulbecco’s modified Eagle’s medium
DNA: deoxyribonucleic acid
DTT: Dithiothreitol
EB: elementary body
F: female or far-periphery
FBS: fetal bovine serum
GCL: ganglion cell layer
gDNA: genomic DNA
GFAP: glial fibrillary acidic protein
GO BP: gene ontology biological process
HRP: horseradish peroxidase
I: inferior
IBA1: ionized calcium-binding adaptor molecule 1
IFN: interferon
IL: interleukin
INL: inner nuclear layer
IPL: inner plexiform layer
IR: immunoreactivity
IRB: institutional review board
IT: inferior-temporal
IN: inferior-nasal
KEGG: Kyoto encyclopedia of genes and genomes
LPS: lipopolysaccharides
M: male or mid-periphery or macula
mAb: monoclonal antibody
MCI: mild cognitive impairment
MHC II: major histocompatibility class 2
MOCA: Montreal cognitive assessment
MMSE: mini-mental state examination
NFTs: neurofibrillary tangles
MOMP: major outer-membrane porin
MS: mass spectrometry
N: nasal
n.a.: not available
N/A: not applicable
NC: normal cognition
NDRI: national disease research interchange
NF-κB: nuclear factor kappa-light-chain-enhancer of activated B cells
NGSDMD: N-terminus gasdermin D
NLR: nucleotide-binding domain (NOD) leucine rich containing protein
NLRP3: NLR family pyrin domain containing 3
NMDAR: N-methyl-D-aspartate receptor
ns: not significant
NS: nasal superior
APES: Aminopropyltriethoxysilane
NTs: neuropil threads
OD: optic disc
ONL: outer nuclear layer
OS: outer segment
pAb: polyclonal antibody
PBS: phosphate-buffered saline
PHF: paired helical filament
PSD95: postsynaptic density protein 95
PCR: polymerase chain reaction
qPCR: quantitative PCR
r: retinal
RB: reticulate body
Ret.: retinal
RFU: relative fluorescence unit
RGC: retina ganglion cells
RNFL: retinal nerve fiber layer
*r*_p_: Pearson’s correlation coefficient
RNA: ribonucleic acid
rRNA: ribosomal RNA
*r*_s_: Spearmen correlation coefficient
S: superior
SPG: sucrose phosphate buffer
ST: superior-temporal
SYNAP: synaptophysin
T: temporal
T22: oligomeric tau 22
TEA: Triethanolamine
TGFβ: transforming growth factor β
ThioS: Thioflavin-S
TLR: toll-like receptor
TNF: tumor necrosis factor
Tx: treatment
UCS: university of California South
VGlut: vesicular glutamate transporter

## Acknowledgements

We acknowledge Dr. Carol Miller for providing a portion of the human tissues and neuropathological reports. The article is dedicated to the memory of Dr. Salomon Moni Hamaoui and Lillian Jones Black, both of whom passed away from Alzheimer’s disease.

## Author Contribution

Study conception and design: BPG, YK, TC, MKH; data acquisition: BPG, YK, JPV, AH, NS, DTF, SS, AR, LS, ER, LSS, DH, MM, MKH; tissue isolation, processing, and histological and biochemical analyses: YK, BPG, AR, DTF, NS, SS, AVL, LSS, DH, MKH; mass spectrometry and data analysis: JPV, MM, BPG, YK, MKH; machine learning analysis: AH, JGM, BPG, MKH; statistical analysis: BPG, YK, DTF, JPV, AH, SS, JGM, MKH; interpretation of data and discussion: BPG, YK, TC, MA, SLG, VKG, KLB, MM, MA, TC, MKH; manuscript writing: MKH, BPG, YK; manuscript editing and revision: BPG, YK, JPV, AH, JGM, MM, AVL, LSS, SLG, VKG, MA, TC, MKH; study supervision: MKH. All authors read and approved the final manuscript.

## Data availability statement

Most data generated or analyzed for this study are included in this manuscript and supplementary material. Data generated for multivariable analyses are available on GitHub. All the processed proteomics data generated in this study have been included in the manuscript and the online supplementary materials. The mass spectrometry raw files and search results have been deposited to the ProteomeXchange Consortium via the PRIDE partner repository with the dataset identifier PXD040225. Additional data are available from the corresponding author upon reasonable request.

