## Supplementary materials for "Identification of *Chlamydia pneumoniae* and NLRP3 inflammasome activation in Alzheimer’s disease retina"

**Supplementary figures and legends:**
**Supplementary Table 1**. Detail neuropathological reports of human retinal and brain donors for histological study.
**Supplementary Table 2.** List of human donors whose postmortem retinas were used for mass spectrometry.
**Supplementary Table 3.** List of human donors whose postmortem brains were used for mass spectrometry.
**Supplementary Table 4**. Demographic data on human retinal donors for mass spectrometry analysis.
**Supplementary Table 5.** Demographic data on human brain donors for mass spectrometry analysis**.**
**Supplementary Table 6.** List of antibodies for immunohistochemistry.
**Supplementary Table 7**: Correlation of retinal Cp burden versus retinal and brain AD pathologies and cognition.
**Supplementary Table 8**: Correlation of brain Cp burden versus brain AD pathologies and cognition.
**Supplementary Table 9.** Upregulated and downregulated DEPs in AD versus NC brains from *Chlamydia* interactome.
**Supplementary Table 10.** Upregulated and downregulated DEPs in AD versus NC retina from *Chlamydia* interactome.
**Supplementary Table 11**: Pearson's correlation analyses of markers for retinal inflammasome, cell degeneration, and gliosis with retinal AD pathology markers.
**Supplementary Table 12**: Spearman's correlation analysis of markers for retinal inflammasome, cell degeneration, and gliosis with brain AD pathology and cognition
**Supplementary Figure 1.** Cp inclusions in retinal or brain cross-sections.
**Supplementary Figure 2.** Distribution of Cp signals across different retinal regions and sex.
**Supplementary Figure 3.** Cell death and immune response pathways, and association with Chlamydia infection in the retina.
**Supplementary Figure 4.** Cell death and immune response pathways, AD neuropathology, and association with Chlamydia infection in the retina and the brain.
**Supplementary Figure 5.** Correlation of Chlamydia inclusion interactors with amyloid plaque and neurofibrillary tangle burden in the AD retina.
**Supplementary Figure 6.** Association of retinal Cp with retinal NLRP3 inflammasome components, early apoptosis, and cellular pyroptosis.
**Supplementary Figure 7.** Association of retinal Cp with retinal gliosis.
**Supplementary Figure 8.** Association of retinal Cp with retinal microgliosis.
**Supplementary Figure 9.** Prediction of brain AD pathologies by retinal Cp, NLRP3, cleaved caspase-3, and $A\beta_{42}$.
**Supplementary Figure 10.** The AUC box plots across all diagnostic groups.
**Supplementary Figure 11.** The AUC box plots across each diagnostic groups.
**References**.

**Supplementary Table 1**. Detail neuropathological reports of human retinal and brain donors for histological study.

| Donor | Sex | Race | Age at Death | Thal (A) | Braak (B) | CERAD (C) | CAA Score | Braak Stage | CDR Score | MMSE Score | APOE status | MOCA |
|---|---|---|---|---|---|---|---|---|---|---|---|---|
| NC1 | F | W | 93 | 3 | 2 | 3 | 0 | 3.5 | 1 | 27 | e3/e2 | n.a. |
| NC2 | F | H | 85 | 2 | 1 | 2 | 0 | 1.5 | 0 | 30 | e3/e3 | n.a. |
| NC3 | M | H | 81 | 3 | 1 | 2 | 0 | 1.5 | 0 | 23 | e3/e4 | 23 |
| NC4 | F | W | 99 | 1 | 2 | 1 | 0 | 3 | 0 | 29 | e3/e3 | n.a. |
| NC5 | M | W | 95 | 1 | 1 | 1 | 0 | 1 | 0 | 30 | e3/e3 | 27 |
| NC6 | M | H | 76 | 2 | 0 | 2 | 0 | 0 | 0 | 29 | e3/e3 | 27 |
| NC7 | M | W | 69 | 0 | 0 | 1 | 0 | 0 | 1 | 28 | n.a. | n.a. |
| NC8 | F | W | 95 | 3 | 3 | 2 | 1 | 5 | 0 | 30 | e3/e3 | n.a. |
| NC9 | F | W | 95 | 1 | 0 | 0 | 0.5 | 1 | 0 | 30 | n.a. | n.a. |
| NC10 | F | W | 98 | n.a. | n.a. | n.a. | n.a. | n.a. | n.a. | n.a. | n.a. | n.a. |
| NC11 | F | n.a. | 95 | n.a. | n.a. | n.a. | n.a. | n.a. | n.a. | n.a. | n.a. | n.a. |
| NC12 | F | n.a. | 91 | n.a. | n.a. | n.a. | n.a. | n.a. | n.a. | n.a. | n.a. | n.a. |
| NC13 | M | W | 77 | n.a. | n.a. | n.a. | n.a. | n.a. | n.a. | 30 | n.a. | n.a. |
| NC14 | F | W | 58 | n.a. | n.a. | n.a. | n.a. | n.a. | n.a. | 30 | n.a. | n.a. |
| NC15 | M | W | 84 | n.a. | n.a. | n.a. | n.a. | n.a. | n.a. | 30 | n.a. | n.a. |
| NC16 | M | W | 70 | n.a. | n.a. | n.a. | n.a. | n.a. | n.a. | 30 | n.a. | n.a. |
| NC17 | F | W | 88 | n.a. | n.a. | n.a. | n.a. | n.a. | n.a. | n.a. | n.a. | n.a. |
| NC18 | M | W | 74 | n.a. | n.a. | n.a. | n.a. | n.a. | n.a. | n.a. | n.a. | n.a. |
| NC19 | M | W | 87 | n.a. | n.a. | n.a. | n.a. | n.a. | n.a. | 30 | n.a. | n.a. |
| NC20 | M | B | 80 | n.a. | n.a. | n.a. | n.a. | n.a. | n.a. | n.a. | n.a. | n.a. |
| NC21 | F | W | 86 | n.a. | n.a. | n.a. | n.a. | n.a. | n.a. | n.a. | n.a. | n.a. |
| MCI1 | F | W | 86 | 3 | 1 | 3 | 0 | 1.5 | 2 | 15 | e3/e4 | 26 |
| MCI2 | F | W | 93 | 3 | 2 | 2 | 2 | 4 | 3 | 11 | e3/e3 | n.a. |
| MCI3 | M | W | 93 | 2 | 0 | 2 | 0 | 0 | 3 | 19 | e3/e2 | n.a. |
| MCI4 | M | W | 97 | 2 | 3 | 3 | 1 | 5 | 1 | 28 | e3/e3 | n.a. |
| MCI5 | F | B | 94 | 2 | 1 | 2 | 0 | 2 | 0.5 | 29 | e3/e3 | 23 |
| MCI6 | F | W | 89 | 1 | 2 | 2 | 1 | 4 | 0.5 | 24 | e3/e3 | 21 |
| MCI7 | M | W | 83 | 2 | 3 | 2 | 1.5 | 3.5 | 0 | 26 | n.a. | n.a. |
| MCI8 | F | W | 91 | 2 | 2 | 2 | 0 | 3 | 3 | 29 | n.a. | n.a. |
| MCI9 | F | W | 98 | 3 | 3 | 2 | 2 | 5 | 2 | 15 | n.a. | n.a. |
| MCI10 | F | W | 87 | 3 | 3 | 3 | 1.5 | 5.5 | 3 | 13 | e3/e3 | n.a. |
| MCI11 | M | W | 88 | 1 | 2 | 2 | 0 | 3 | 3 | n.a. | n.a. | 24 |
| MCI12 | M | H | 80 | 3 | 3 | 2 | n.a. | 5 | 3 | 29 | e3/e3 | n.a. |
| MCI13 | M | W | 75 | n.a. | n.a. | n.a. | n.a. | n.a. | n.a. | n.a. | n.a. | n.a. |
| MCI14 | M | W | 90 | n.a. | n.a. | n.a. | n.a. | n.a. | n.a. | n.a. | n.a. | n.a. |

| Donor | Sex | Race | Age at Death | Thal (A) | Braak (B) | CERAD (C) | CAA Score | Braak Stage | CDR Score | MMSE Score | APOE status | MOCA |
|---|---|---|---|---|---|---|---|---|---|---|---|---|
| MCI15* | M | H | 85 | 0 | 1 | 3 | 1 | 1.5 | 0.5 | n.a. | n.a. | n.a. |
| AD1 | F | W | 90 | 2 | 3 | 3 | 1 | 5 | 2 | 9 | n.a. | n.a. |
| AD2 | F | W | 100 | 2 | 3 | 3 | 1 | 5.5 | 2 | 16 | n.a. | n.a. |
| AD3 | M | W | 90 | 3 | 3 | 3 | 2 | 6 | 3 | n.a. | n.a. | n.a. |
| AD4 | M | W | 88 | 2 | 3 | 2 | 1 | 5.5 | 1 | 4 | e3/e4 | 18 |
| AD5 | F | W | 87 | 2 | 3 | 3 | 1.5 | 5 | 3 | 16 | e3/e4 | n.a. |
| AD6 | F | W | 70 | 3 | 3 | 3 | 1.5 | 5 | 0.5 | 24 | n.a. | n.a. |
| AD7 | M | W | 77 | 3 | 3 | 3 | 1 | 6 | 2 | 18 | e3/e4 | n.a. |
| AD8 | F | W | 66 | 3 | 3 | 3 | 0 | 6 | 3 | 2 | e3/e3 | n.a. |
| AD9 | M | A | 81 | 3 | 3 | 3 | 1 | 5 | 3 | 12 | e4/e4 | n.a. |
| AD10 | F | H | 99 | 3 | 3 | 3 | 1.5 | 4 | 3 | 23 | e3/e3 | n.a. |
| AD11 | F | W | 85 | 3 | 3 | 3 | 1.5 | 5.5 | 3 | n.a. | e3/e3 | n.a. |
| AD12 | F | W | 81 | 3 | 3 | 3 | 2 | 6 | n.a. | 4 | n.a. | n.a. |
| AD13 | M | W | 83 | 3 | 2 | 3 | 2 | 4 | 1 | 18 | n.a. | n.a. |
| AD14 | M | H | 65 | 3 | 3 | 3 | 1.5 | 5 | 3 | n.a. | e4/e4 | n.a. |
| AD15 | F | H | 92 | 2 | 3 | 2 | 2 | 5 | 3 | 9 | n.a. | n.a. |
| AD16 | F | A | 88 | 2 | 3 | 3 | 1.5 | 5 | 3 | 4 | n.a. | n.a. |
| AD17 | M | W | 66 | 3 | 3 | 3 | 1.5 | 5 | 3 | 19 | n.a. | n.a. |
| AD18 | F | W | 86 | 3 | 3 | 2 | 3 | 5.5 | 3 | 18 | e3/e4 | n.a. |
| AD19 | M | W | 88 | 3 | 3 | 3 | 1.5 | 5.5 | 1 | 16 | e2/e3 | n.a. |
| AD20 | F | A | 93 | 2 | 2 | 2 | 1.5 | 3.5 | 3 | 17 | n.a. | n.a. |
| AD21 | F | W | 94 | 3 | 3 | 3 | 0 | 5.5 | 3 | n.a. | e3/e3 | n.a. |
| AD22 | F | H | 81 | 3 | 3 | 3 | 1.5 | 5.5 | 3 | 12 | e3/e3 | n.a. |
| AD23 | F | W | 90 | 3 | 3 | 3 | 1 | 5.5 | 3 | n.a. | e3/e4 | n.a. |
| AD24 | M | W | 90 | 3 | 2 | 3 | 1 | 4 | 3 | n.a. | e3/e4 | 1 |
| AD25 | F | W | 93 | 2 | 3 | 3 | 1 | 5 | 3 | n.a. | n.a. | n.a. |
| AD26 | M | W | 79 | 3 | 3 | 3 | 1.5 | 5 | n.a. | n.a. | n.a. | n.a. |
| AD27 | F | W | 87 | 3 | 3 | 3 | 2 | 5.5 | 3 | n.a. | e3/e4 | 9 |
| AD28 | M | W | 88 | 2 | 2 | 2 | 0 | 3.5 | n.a. | 4 | n.a. | n.a. |
| AD29 | M | H | 97 | 3 | 2 | 3 | 1 | 3 | 1 | 26 | e3/e3 | n.a. |
| AD30 | M | W | 90 | 3 | 3 | 2 | 2 | 5 | 2 | 18 | n.a. | n.a. |
| AD31 | M | W | 85 | 3 | 2 | 3 | 1 | 4 | n.a. | n.a. | n.a. | n.a. |
| AD32 | M | W | 99 | n.a. | n.a. | n.a. | n.a. | n.a. | n.a. | n.a. | n.a. | n.a. |
| AD33 | M | n.a. | 90 | n.a. | n.a. | n.a. | n.a. | n.a. | n.a. | n.a. | n.a. | n.a. |
| AD34 | M | H | 97 | 3 | 2 | 3 | 1 | 3 | 1 | 26 | e3/e3 | n.a. |

AD, Alzheimer's disease dementia; MCI, mild cognitive impairment; NC, normal cognition; F, female; M, male; A, Asian; B, Black; H, Hispanic; W, White; A, Aβ plaque score modified from Thal; B, NFT stage modified from Braak; C, neuritic plaque score modified from CERAD; CAA, cerebral amyloid angiopathy; CDR, clinical dementia rating; MMSE, mini-mental state examination; MOCA: Montreal cognitive assessment; n.a., not available; APOE, apolipoprotein alleles. *Brain tissues only.

**Supplementary Table 2.** List of human donors whose postmortem retinas were used for mass spectrometry.

| Donor | Sex | Race | Age at Death | Thal (A) | Braak (B) | CERAD (C) | CAA Score | Braak Stage | CDR Score | MMSE Score | APOE Status |
|---|---|---|---|---|---|---|---|---|---|---|---|
| AD1 | F | W | 93 | 2 | 3 | 3 | 1 | 5 | 3 | n.a. | n.a. |
| AD2 | M | W | 88 | 2 | 3 | 2 | 1 | 5.5 | 1 | 4 | e3/e4 |
| AD3 | F | B | 94 | 3 | 3 | 3 | 0 | 5.5 | 3 | n.a. | e3/e3 |
| AD4 | F | H | 48 | n.a. | n.a. | n.a. | 2 | 5 | n.a. | n.a. | n.a. |
| AD5 | M | W | 72 | n.a. | n.a. | n.a. | n.a. | n.a. | n.a. | n.a. | n.a. |
| AD6 | F | W | 100 | 2 | 3 | 3 | 1 | 5.5 | 2 | 16 | n.a. |
| NC1 | M | H | 81 | 3 | 1 | 2 | 0 | 1.5 | 0 | 23 | e3/e4 |
| NC2 | F | W | 75 | n.a. | n.a. | n.a. | n.a. | n.a. | n.a. | n.a. | n.a. |
| NC3 | F | W | 72 | n.a. | n.a. | n.a. | n.a. | n.a. | n.a. | n.a. | n.a. |
| NC4 | M | W | 69 | n.a. | n.a. | n.a. | n.a. | n.a. | n.a. | n.a. | n.a. |
| NC5 | F | W | 79 | n.a. | n.a. | n.a. | n.a. | n.a. | n.a. | n.a. | n.a. |
| NC6 | M | W | 85 | 0 | 1 | 3 | 1 | 1.5 | 0.5 | n.a. | n.a. |

*Abbreviations*: AD, Alzheimer's disease; APOE, apolipoprotein, B, Black; A (Thal), Aβ plaque score modified from Thal; B (Braak), NFT stage modified from Braak; C (CERAD), neuritic plaque score modified from CERAD; CAA, cerebral amyloid angiopathy; CDR, clinical dementia rating; CERAD, consortium to establish a registry for Alzheimer's disease; F, female; H, Hispanic; M, male; MMSE, mini-mental state examination; n.a., not available; NC, normal cognition; W, White. All human donor tissues obtained from UCI-ADRC.

**Supplementary Table 3.** List of human donors whose postmortem brains were used for mass spectrometry.

| Diagnosis | Sex | Race | Age at Death | Plaque Stage | Braak Stage | Neuritic Plaque | MMSE Score | APOE Status |
|---|---|---|---|---|---|---|---|---|
| AD1 | M | W | 86 | C | 6 | 4 | 19 | 3/4 |
| AD2 | F | W | 89 | C | 6 | 4 | 15 | 3/4 |
| AD3 | F | W | 95 | C | 6 | 3 | n.a. | 2/3 |
| AD4 | F | W | 93 | C | 6 | 4 | 13 | 3/3 |
| AD5 | F | W | 88 | C | 5 | 4 | 10 | 3/3 |
| AD6 | M | W | 92 | C | 5 | 4 | 12 | 3/3 |
| AD7 | M | H | 86 | C | 5 | 4 | n.a. | 3/3 |
| AD8 | F | W | 98 | C | 6 | 4 | 14 | 3/3 |
| AD9 | F | W | 91 | C | 6 | 4 | 17 | 3/3 |
| AD10 | F | W | 82 | C | 6 | 4 | 17 | 3/3 |
| NC1 | F | W | 87 | 0 | 2 | 1 | 30 | 2/3 |
| NC2 | F | W | 91 | 0 | 4 | 1 | 29 | 2/3 |
| NC3 | F | H/AI | 95 | 0 | 2 | 1 | n.a. | 3/3 |
| NC4 | F | W | 90 | B | 3 | 3 | 18 | 3/3 |
| NC5 | M | W | 96 | 0 | 2 | 1 | 29 | 2/3 |
| NC6 | M | W | 94 | 0 | 1 | 1 | 27 | 3/3 |
| NC7 | M | W | 86 | 0 | 2 | 2 | 27 | 3/3 |
| NC8 | F | W | 91 | A | 2 | 2 | 29 | 3/3 |

*Abbreviations*: AD, Alzheimer's disease; APOE, apolipoprotein; AI, American Indian; F, female; H, Hispanic; M, male; MMSE, Mini-Mental State Examination; n.a., not available; NC, normal cognition; Plaque Stage: 0, none; A, phase I-II; B, phase III; C, phase IV-V; Neuritic Plaque scores: 4, frequent NP; 3, moderate NP; 2, sparce NP; 1, no NPs. All human donor tissues obtained from UCI-ADRC.

**Supplementary Table 4**. Demographic data on human retinal donors for mass spectrometry analysis.

| | | NC | AD | F | *p* |
|---|---|---|---|---|---|
| **n = 12** | | 6 (3F, 3M) | 6 (4F, 2M) | - | - |
| **Age at Death (Years)** | | 76.8 ± 5.9 | 82.5 ± 19.4 | 0.68 | 0.52 |
| **Race** | | 5W, 1H | 4W, 1B, 1H | - | - |
| **PMI (Hours)** | | 9.0 ± 5.1 | 7.8 ± 2.7 | 0.62 | 0.57 |
| **MMSE Score (n = 3)** | | 23.0 ± n.a | 17.0 ± 1.4 | - | **-** |
| **CDR Score (n = 6)** | | 0.25 ± 0.35 | 2.25 ± 0.96 | 2.72 | 0.053 |
| **Brain Neuropathology Severity Score (n = 8)** | Braak stage (%) | I-II (100%) | V-VI (100%) | 31.03 | **<0.0001** |
| | ABC (amyloid, Braak, CERAD) (n = 6) | 1.83 ± 0.71 | 2.67 ± 0.27 | 1.61 | 0.33 |
| | Aβ plaque | 1.76 ± 0.68 | 3.09 ± 1.26 | 1.90 | 0.14 |
| | NFTs | 1.15 ± 0.01 | 2.50 ± 0.96 | 3.45 | **0.0182** |
| | NTs | 0.22 ± 0.11 | 1.43 ± 0.43 | 5.91 | **0.0022** |
| | Atrophy | 0.55 ± 0.07 | 1.41 ± 0.97 | 2.16 | 0.08 |

Mean ABC scores were determined as follows: A, Aβ plaque score modified from Thal; B, NFT stage modified from Braak; C, neuritic plaque score modified from CERAD.

Group values are presented as mean ± standard deviation. F and *p*-values were determined using an unpaired Student t test. *p*-values presented in bold type demonstrate significance.

*Abbreviations*: Aβ, amyloid beta; AD, Alzheimer's disease; CDR, clinical dementia rating; CERAD, consortium to establish a registry for Alzheimer's disease; NC, normal cognition; MMSE, mini-mental state examination; NFTs, neurofibrillary tangles; NTs, neuropil threads; PMI, postmortem interval.

**Supplementary Table 5.** Demographic data on human brain donors for mass spectrometry analysis**.**

| Human Donors | | NC | AD | *t* | *p* |
|---|---|---|---|---|---|
| **n = 18** | | **8** (5F, 3M) | **10** (7F, 3M) | - | - |
| **Age at Death** [Years]§ | | **91.3** ± 3.6 | **90.0** ± 4.8 | 0.61 | *0.55* |
| **Race** [No] | | W (7)<br>H/A (1) | W (9)<br>H (1) | - | - |
| **MMSE Score**§ | | 27.0 ± 4.1 | 14.6 ± 2.9 | 6.74 | ***<0.0001*** |
| **PMI** [Hours]§ | | 3.7 ± 0.8 | 5.3 ± 3.2 | 1.43 | *0.17* |
| **Brain Neuropathology Severity Score** (n = 18) | **Aβ-Plaque Stage** (No; %) | None (6; 75%)<br>Stage A (1; 12.5%)<br>Stage B (1; 12.5%) | Stage C (10; 100%) | - | - |
| | **Braak Stage** (NO; %) | I-II (6; 75%)<br>III-IV (2; 25%)<br>V-VI (0) | V-VI (10; 100%) | - | - |

*Abbreviations*: Aβ, Amyloid-β protein; Aβ-Plaque stages score: None, no Aβ plaque or amyloid plaque; A, mild, A1 Thal phases 1 or 2; B, moderate, A2 Thal phase 3; C, severe plaque pathology, A3 Thal phases 4 or 5; AD, Alzheimer's disease; Braak (NFT) stage scores; MMSE, mini-mental state examination; NC, normal cognition; NFT, neurofibrillary tangles; PMI, postmortem interval; Values are presented as mean ± SD. The *t* and *p* values were determined by Student's t-test.

**Supplementary Table 6.** List of antibodies for immunohistochemistry.

| Antibodies | Source Species | Dilution | Application | Source | Catalog. # |
|---|---|---|---|---|---|
| *Primary antibody* | | | | | |
| Cp pAb | Rabbit | 1:500 | IF/IHC (DAB) | MyBioSource | MBS534621 |
| Cp mAb | Mouse | 1:50 | IF | invitrogen | MA5-18183 |
| Cp mAb | Mouse | 1:200 | IHC/DAB | invitrogen | MA5-18183 |
| NLRP3 mAb | Rat | 1:100 | IF | R&D systems | MAB7578 |
| Caspase-1 mAb | Rabbit | 1:150 | IF | R&D systems | MAB62156 |
| ASC | Rabbit | 1:300 | IF | | |
| N-GSDMD mAb | Rabbit | 1:500 | IF | Cell Signaling | 36425 |
| CCasp3 pAb | Rabbit | 1:400 | IF | Cell Signaling | 9661 |
| GFAP pAb | Goat | 1:500 | IF | Invitrogen | 13-0300 |
| Iba-1 mAb | Rabbit | 1:400 | IF | Wako | 019-19741 |
| Vimentin | Rabbit | 1:350 | IF | abcam | Ab92547 |
| NeuN mAb | Rabbit | 1:500 | IF | abcam | Ab177487 |
| *Secondary antibody* | | | | | |
| Cy3 (anti-mouse, anti-rat,) | Donkey | 1:200 | IF | Jackson ImmunoResearch Laboratories | 715-165-150, 712-165-153 |
| Cy5 (anti-rabbit) | Donkey | 1:200 | IF | Jackson ImmunoResearch Laboratories | 711-175-152 |
| Cy2 (anti-goat) | Donkey | 1:200 | IF | Jackson ImmunoResearch Laboratories | |
| HRP (anti-mouse, anti-rabbit) | Goat | - | IHC/DAB | DAKO invision | K4001, K4003 |

*Abbreviations*: ASC - apoptosis-associated speck like protein; Cp – *Chlamydia pneumoniae*; GFAP – glial fibrillary acidic protein; Iba1 – ionized calcium binding adaptor molecule 1; CCasp3 – Cleaved Caspase-3; DAB - 3,3-Diaminobenzidine; HRP - Horseradish peroxidase; IF – immunofluorescence; IHC – immunohistochemistry; mAb – monoclonal antibody; NGSDMD – Gasdermin D with cleaved N-terminus; NLRP3 - nucleotide-binding oligomerization domain-like receptor containing pyrin domain 3; pAb – polyclonal antibody.

**Supplementary Table 7**: Correlation of retinal Cp burden versus retinal and brain AD pathologies and cognition

| **Pearson's correlation (*$r_p$*) analysis: Retinal Cp versus retinal AD pathologies** | | | | | |
|---|---|---|---|---|---|
| $A\beta_{42}$ | $A\beta_{40}$ | Aβ oligomers | pS396 | PHF-1 tau | Oligo-tau (T22) |
| **0.63****** **n = 39** | **0.65**** **n = 21** | 0.18 n = 22 | 0.38* n = 44 | 0.54** n = 23 | 0.43** n = 43 |
| Cit-tau | AT8-tau | MC-1 tau | NLRP3 | Caspase-1 | ASC |
| 0.48** n = 37 | 0.02 n = 39 | 0.04 n = 43 | **0.70****** **n = 27** | **0.87****** **n = 27** | **0.60***** **n = 27** |
| NGSDMD | CCasp3 | IBA1 | GFAP | Vimentin | S100β |
| 0.51** n = 27 | **0.73****** **n = 25** | **0.65****** **n = 32** | **0.70****** **n = 28** | 0.55** n = 21 | 0.36 n = 6 |
| Nissl loss | Atrophy | | | | |
| -0.43* n = 34 | **0.75****** **n = 23** | | | | |
| **Spearman's correlation (*$r_s$*): Retinal Cp versus brain AD pathologies** | | | | | |
| ABC | Braak Stage | Aβ plaques | NFTs | NTs | Gliosis |
| 0.54**** n = 60 | 0.54**** n = 60 | 0.40** n = 60 | 0.54**** n = 60 | 0.37** n = 60 | 0.40** n = 60 |
| Atrophy | CAA | | | | |
| 0.48*** n = 60 | 0.35** n = 60 | | | | |
| **Spearman's correlation (*$r_s$*): Retinal Cp versus cognition** | | | | | |
| CDR | MMSE | MoCA | | | |
| 0.43** n = 56 | -0.53**** n = 50 | -0.56* n = 15 | | | |

Pearson's and Spearman's rank correlation analyses: *p* and *r* values determine the statistical significance and strength of each pairwise association between retinal Cp burden versus retinal and brain AD pathologies and cognition. *p* and *r* values presented in bold type with asterisk(s) depicts strong to very strong correlation.

**Supplementary Table 8**: Correlation of brain Cp burden versus brain AD pathologies and cognition

| **Spearman's correlation ($r_s$): Brain Cp versus Brain AD pathologies** | | | | | |
|---|---|---|---|---|---|
| ABC | Braak Stage | Aβ plaques | NFTs | NTs | Gliosis |
| **0.74**** <br> **n = 16** | **0.72**** <br> **n = 16** | 0.45 <br> n = 16 | **0.73**** <br> **n = 16** | **0.75**** <br> **n = 16** | **0.77***** <br> **n = 16** |
| Atrophy | CAA | | | | |
| **0.60*** <br> **n = 16** | 0.57* <br> n = 16 | | | | |
| **Spearman's correlation ($r_s$): Brain Cp versus cognition** | | | | | |
| CDR | MMSE | MoCA | | | |
| 0.50 <br> n = 16 | **-0.73**** <br> **n = 14** | -0.52 <br> n = 6 | | | |

Spearman's rank correlation analyses: *p* and *r* values determine the statistical significance and strength of each pairwise association between brain Cp burden versus brain AD pathologies and cognition. *p* and *r* values presented in bold type with asterisk(s) depicts strong to very strong correlation.

**Supplementary Table 9.** Upregulated and downregulated DEPs in AD versus NC brains from *Chlamydia* interactome.

| Accession | Symbol | Description | FC | *p* | References |
|---|---|---|---|---|---|
| **Upregulated in AD brain (32)** | | | | | |
| Q13501 | **SQSTM1** | Sequestosome-1 | **1.48** | 0.0001 | [1,2] |
| Q08379 | **GOLGA2** | Golgin subfamily A member 2 | **1.47** | 0.0190 | [3] |
| O43291 | **SPINT2** | Kunitz-type protease inhibitor 2 | **1.45** | 0.0439 | [1] |
| P08670 | **VIM** | Vimentin | **1.45** | 0.0013 | [2] |
| E9PND2 | **CSRP1** | Cysteine and glycine-rich protein 1 | **1.37** | 0.0411 | [3] |
| Q8N4Q1 | **CHCHD4** | Mitochondrial intermembrane space import and assembly protein 40 | **1.36** | 0.0462 | [4] |
| Q15796 | **SMAD2** | Mothers against decapentaplegic homolog 2 | **1.36** | 0.0329 | [3] |
| Q9Y676 | **MRPS18B** | 28S ribosomal protein S18b, mitochondrial | **1.34** | 0.0069 | [4] |
| O95817 | **BAG3** | BAG family molecular chaperone regulator 3 | **1.34** | 0.0243 | [2] |
| G3V1V0 | **MYL6** | Myosin light polypeptide 6 | **1.34** | 0.0007 | [2] |
| P06703 | **S100A6** | Protein S100-A6 | **1.34** | 0.0167 | [1] |
| A1L188 | **NDUFAF8** | NADH dehydrogenase [ubiquinone] 1 alpha subcomplex assembly factor 8 | **1.28** | 0.0485 | [4] |
| O75348 | **ATP6V1G1** | V-type proton ATPase subunit G 1 | **1.28** | 0.0007 | [1] |
| Q14019 | **COTL1** | Coactosin-like protein | **1.28** | 0.0026 | [1] |
| P21333 | **FLNA** | Filamin-A | **1.27** | 0.0237 | [2] |
| Q9UBI6 | **GNG12** | Guanine nucleotide-binding protein G(I)/G(S)/G(O) subunit gamma-12 | **1.27** | 0.0195 | [1] |
| P56385 | **ATP5ME** | ATP synthase subunit e, mitochondrial | **1.27** | 0.0103 | [4] |
| P04792 | **HSPB1** | Heat shock protein beta-1 | **1.27** | 0.0082 | [1,2] |
| P04080 | **CSTB** | Cystatin-B | **1.26** | 0.0075 | [3] |
| P17931 | **LGALS3** | Galectin-3 | **1.26** | 0.0027 | [2] |
| Q96HC4 | **PDLIM5** | PDZ and LIM domain protein 5 | **1.25** | 0.0118 | [3] |
| O75369 | **FLNB** | Filamin-B | **1.25** | 0.0429 | [3] |
| Q6DD88 | **ATL3** | Atlastin-3 | **1.24** | 0.0024 | [1,4] |
| J3KN67 | **TPM3** | Tropomyosin alpha-3 chain | **1.24** | 0.0103 | [1-3] |
| P61254 | **RPL26** | 60S ribosomal protein L26 | **1.22** | 0.0483 | [3] |
| Q15691 | **MAPRE1** | Microtubule-associated protein RP/EB family member 1 | **1.22** | 0.0003 | [4] |
| Q12797 | **ASPH** | Aspartyl/asparaginyl beta-hydroxylase | **1.21** | 0.0205 | [1,4] |
| Q9BQT9 | **CLSTN3** | Calsyntenin-3 | **1.21** | 0.0171 | [4] |
| Q9C0C2 | **TNKS1BP1** | 182 kDa tankyrase-1-binding protein | **1.21** | 0.0054 | [2] |
| Q09666 | **AHNAK** | Neuroblast differentiation-associated protein AHNAK | **1.22** | 0.0342 | [2,3] |
| Q5TZA2 | **CROCC** | Rootletin | **1.22** | 0.0395 | [1] |
| P05556 | **ITGB1** | Integrin beta-1 | **1.20** | 0.0065 | [2] |
| **Downregulated in AD brain (52)** | | | | | |

| | | | | | |
|---|---|---|---|---|---|
| Q9NZ01 | **TECR** | Very-long-chain enoyl-CoA reductase | **-2.05** | 0.0246 | [1] |
| Q9NQC3 | **RTN4** | Reticulon-4 | **-1.83** | 0.0039 | [1,3,4] |
| Q9UGP8 | **SEC63** | Translocation protein SEC63 homolog | **-1.70** | 0.0089 | [1] |
| Q6IA86 | **ELP2** | Elongator complex protein 2 | **-1.62** | 0.0209 | [4] |
| E9PK54 | **HSPA8** | Heat shock cognate 71 kDa protein | **-1.53** | 0.0411 | [1] |
| Q9BVC4 | **MLST8** | Target of rapamycin complex subunit LST8 | **-1.51** | 0.0263 | [4] |
| O43760 | **SYNGR2** | Synaptogyrin-2 | **-1.49** | 0.0208 | [1,3] |
| P61204 | **ARF3** | ADP-ribosylation factor 3 | **-1.47** | 0.0128 | [1] |
| Q9P035 | **HACD3** | Very-long-chain (3R)-3-hydroxyacyl-CoA dehydratase 3 | **-1.43** | 0.0372 | [1] |
| P61225 | **RAP2B** | Ras-related protein Rap-2b | **-1.40** | 0.0007 | [1] |
| Q8TCJ2 | **STT3B** | Dolichyl-diphosphooligosaccharide--protein glycosyltransferase subunit | **-1.39** | 0.0185 | [1,4] |
| O15258 | **RER1** | Protein RER1 | **-1.37** | 0.0263 | [1,4] |
| O43747 | **AP1G1** | AP-1 complex subunit gamma-1 | **-1.37** | 0.0196 | [1] |
| O95716 | **RAB3D** | Ras-related protein Rab-3D | **-1.36** | 0.0000 | [1] |
| P0CG08 | **GPR89B** | Golgi pH regulator B | **-1.36** | 0.0080 | [4] |
| K7EJH8 | **ACTN4** | Alpha-actinin-4 | **-1.33** | 0.0062 | [1,3] |
| P63096 | **GNAI1** | Guanine nucleotide-binding protein G(i) subunit alpha-1 | **-1.33** | 0.0034 | [3] |
| Q92905 | **COPS5** | COP9 signalosome complex subunit 5 | **-1.32** | 0.0015 | [4] |
| O15144 | **ARPC2** | Actin-related protein 2/3 complex subunit 2 | **-1.32** | 0.0133 | [1] |
| P07099 | **EPHX1** | Epoxide hydrolase 1 | **-1.32** | 0.0340 | [1] |
| F2Z2X4 | **XPO4** | Exportin-4 | **-1.31** | 0.0026 | [4] |
| Q7Z7H5 | **TMED4** | Transmembrane emp24 domain-containing protein 4 | **-1.31** | 0.0230 | [1] |
| P15531 | **NME1** | Nucleoside diphosphate kinase A | **-1.28** | 0.0028 | [3] |
| Q01581 | **HMGCS1** | Hydroxymethylglutaryl-CoA synthase, cytoplasmic | **-1.28** | 0.0119 | [4] |
| O00410 | **IPO5** | Importin-5 | **-1.28** | 0.0401 | [4] |
| Q93050 | **ATP6V0A1** | V-type proton ATPase 116 kDa subunit a isoform 1 | **-1.27** | 0.0015 | [1,4] |
| Q9BXS5 | **AP1M1** | AP-1 complex subunit mu-1 | **-1.27** | 0.0120 | [3,4] |
| Q96F07 | **CYFIP2** | Cytoplasmic FMR1-interacting protein 2 | **-1.27** | 0.0030 | [3] |
| P42704 | **LRPPRC** | Leucine-rich PPR motif-containing protein, mitochondrial | **-1.27** | 0.0025 | [4] |
| P60228 | **EIF3E** | Eukaryotic translation initiation factor 3 subunit E | **-1.26** | 0.0438 | [3] |
| O43759 | **SYNGR1** | Synaptogyrin-1 | **-1.26** | 0.0330 | [4] |
| F8VXU5 | **VPS29** | Vacuolar protein sorting-associated protein 29 | **-1.26** | 0.0234 | [1,3] |
| P60981 | **DSTN** | Destrin | **-1.26** | 0.0406 | [3] |
| P20645 | **M6PR** | Cation-dependent mannose-6-phosphate receptor | **-1.25** | 0.0005 | [1] |

| | | | | | |
|---|---|---|---|---|---|
| E9PFW3 | **AP2M1** | AP-2 complex subunit mu | **-1.25** | 0.0097 | [4] |
| A0A0C4DGQ5 | **CAPNS1** | Calpain small subunit 1 | **-1.25** | 0.0185 | [1,4] |
| Q9BTE1 | **DCTN5** | Dynactin subunit 5 | **-1.25** | 0.0007 | [4] |
| P68104 | **EEF1A1** | Elongation factor 1-alpha | **-1.24** | 0.0061 | [1] |
| P61106 | **RAB14** | Ras-related protein Rab-14 | **-1.24** | 0.0051 | [1] |
| M0QYN0 | **MYDGF** | Myeloid-derived growth factor | **-1.24** | 0.0056 | [1] |
| P00338 | **LDHA** | L-lactate dehydrogenase A chain | **-1.23** | 0.0030 | [1] |
| A0A2U3TZU2 | **GPI** | Glucose-6-phosphate isomerase | **-1.23** | 0.0199 | [1] |
| Q9NZJ7 | **MTCH1** | Mitochondrial carrier homolog 1 | **-1.23** | 0.0097 | [4] |
| Q06136 | **KDSR** | 3-ketodihydrosphingosine reductase | **-1.23** | 0.0235 | [1,4] |
| P20340 | **RAB6A** | Ras-related protein Rab-6A | **-1.22** | 0.0222 | [1] |
| P05388 | **RPLP0** | 60S acidic ribosomal protein P0 | **-1.21** | 0.0086 | [1] |
| Q5VV89 | **MGST3** | Microsomal glutathione S-transferase 3 | **-1.21** | 0.0104 | [3,4] |
| O14617 | **AP3D1** | AP-3 complex subunit delta-1 | **-1.21** | 0.0182 | [4] |
| P61019 | **RAB2A** | Ras-related protein Rab-2A | **-1.21** | 0.0142 | [1] |
| A0A0A0MRA8 | **EPB41L3** | Band 4.1-like protein 3 | **-1.21** | 0.0006 | [4] |
| O95782 | **AP2A1** | AP-2 complex subunit alpha-1 | **-1.20** | 0.0010 | [4] |
| Q08209 | **PPP3CA** | Serine/threonine-protein phosphatase 2B catalytic subunit alpha | **-1.20** | 0.0106 | [3,4] |

FC, fold change. |FC| > 1.2, $p$<0.05.

**Supplementary Table 10.** Upregulated and downregulated DEPs in AD versus NC retinas from *Chlamydia* interactome.

| Accession | Symbol | Description | FC | *p* | References |
|---|---|---|---|---|---|
| **Upregulated in AD retina (40)** | | | | | |
| A6NFX8 | **NUDT5** | ADP-sugar pyrophosphatase | **1.87** | 0.0302 | 3 |
| Q08554 | **DSC1** | Desmocollin-1 | **1.85** | 0.0450 | 1 |
| P15531 | **NME1** | Nucleoside diphosphate kinase A | **1.75** | 0.0386 | 3 |
| P04792 | **HSPB1** | Heat shock protein beta-1 | **1.70** | 0.0054 | 1,2 |
| Q14574 | **DSC3** | Desmocollin-3 | **1.63** | 0.0042 | 4 |
| P67936 | **TPM4** | Tropomyosin alpha-4 chain | **1.62** | 0.0060 | 1-3 |
| J3KN67 | **TPM3** | Tropomyosin alpha-3 chain | **1.53** | 0.0174 | 1-3 |
| P61204 | **ARF3** | ADP-ribosylation factor 3 | **1.52** | 0.0344 | 1 |
| P81605 | **DCD** | Dermcidin | **1.51** | 0.0276 | 1 |
| A0A0B4J2C3 | **TPT1** | Translationally-controlled tumor protein | **1.50** | 0.0156 | 2 |
| A0A087WYT3 | **PTGES3** | Prostaglandin E synthase 3 | **1.50** | 0.0436 | 1 |
| P20810 | **CAST** | Calpastatin | **1.49** | 0.0449 | 1,2 |
| Q02413 | **DSG1** | Desmoglein-1 | **1.45** | 0.0327 | 1 |
| H3BUF6 | **ATXN2L** | Ataxin-2-like protein | **1.39** | 0.0008 | 2 |
| Q01581 | **HMGCS1** | Hydroxymethylglutaryl-CoA synthase, cytoplasmic | **1.37** | 0.0013 | 4 |
| Q9UBE0 | **SAE1** | SUMO-activating enzyme subunit 1 | **1.36** | 0.0261 | 4 |
| Q8TAA9 | **VANGL1** | Vang-like protein 1 | **1.36** | 0.0230 | 4 |
| E9PAV3 | **NACA** | Nascent polypeptide-associated complex subunit alpha, muscle-specific | **1.34** | 0.0175 | 1 |
| O75348 | **ATP6V1G1** | V-type proton ATPase subunit G 1 | **1.33** | 0.0451 | 1 |
| O43852 | **CALU** | Calumenin | **1.33** | 0.0459 | 1 |
| O60493 | **SNX3** | Sorting nexin-3 | **1.32** | 0.0201 | 3 |
| P13797 | **PLS3** | Plastin-3 | **1.32** | 0.0322 | 3 |
| O43399 | **TPD52L2** | Tumor protein D54 | **1.32** | 0.0124 | 1 |
| O95817 | **BAG3** | BAG family molecular chaperone regulator 3 | **1.30** | 0.0225 | 2 |
| P49354 | **FNTA** | Protein farnesyltransferase/geranylgeranyltransferase type-1 subunit alpha | **1.30** | 0.0426 | 4 |
| P48507 | **GCLM** | Glutamate--cysteine ligase regulatory subunit | **1.29** | 0.0106 | 5 |
| Q9NZ08 | **ERAP1** | Endoplasmic reticulum aminopeptidase 1 | **1.28** | 0.0065 | 1 |
| A0MZ66 | **SHTN1** | Shootin-1 | **1.27** | 0.0264 | 3 |
| P60981 | **DSTN** | Destrin | **1.27** | 0.0423 | 3 |
| Q32MZ4 | **LRRFIP1** | Leucine-rich repeat flightless-interacting protein 1 | **1.27** | 0.0087 | 1-4 |
| P31946 | **YWHAB** | 14-3-3 protein beta/alpha | **1.27** | 0.0163 | 1,4 |
| O15355 | **PPM1G** | Protein phosphatase 1G | **1.27** | 0.0025 | 4 |
| P54727 | **RAD23B** | UV excision repair protein RAD23 homolog B | **1.24** | 0.0431 | 4 |
| Q8IYD1 | **GSPT2** | Eukaryotic peptide chain release factor GTP-binding subunit ERF3B | **1.24** | 0.0058 | 3 |
| O75821 | **EIF3G** | Eukaryotic translation initiation factor 3 subunit G | **1.22** | 0.0077 | 3 |

| | | | | | |
|---|---|---|---|---|---|
| P98172 | **EFNB1** | Ephrin-B1 | **1.21** | 0.0325 | 4 |
| Q13200 | **PSMD2** | 26S proteasome non-ATPase regulatory subunit 2 | **1.20** | 0.0168 | 1 |
| O14974 | **PPP1R12A** | Protein phosphatase 1 regulatory subunit 12A | **1.20** | 0.0013 | 1 |
| P27348 | **YWHAQ** | 14-3-3 protein theta | **1.20** | 0.0333 | 1-3 |
| Q99614 | **TTC1** | Tetratricopeptide repeat protein 1 | **1.20** | 0.0464 | 2 |
| **Downregulated in AD retina (52)** | | | | | |
| Q96KR6 | **FAM210B** | Protein FAM210B, mitochondrial | **-1.98** | 0.0101 | 4 |
| P54819 | **AK2** | Adenylate kinase 2, mitochondrial | **-1.53** | 0.0040 | 4 |
| Q5TZA2 | **CROCC** | Rootletin | **-1.49** | 0.0132 | 1 |
| Q9NQC3 | **RTN4** | Reticulon-4 | **-1.48** | 0.0076 | 1,3,4 |
| Q8NBN7 | **RDH13** | Retinol dehydrogenase 13 | **-1.46** | 0.0072 | 4 |
| O95159 | **ZFPL1** | Zinc finger protein-like 1 | **-1.44** | 0.0054 | 4 |
| P61254 | **RPL26** | 60S ribosomal protein L26 | **-1.41** | 0.0324 | 3 |
| Q9Y6A9 | **SPCS1** | Signal peptidase complex subunit 1 | **-1.41** | 0.0052 | 1 |
| Q96AG4 | **LRRC59** | Leucine-rich repeat-containing protein 59 | **-1.41** | 0.0015 | 1,2,4 |
| O94826 | **TOMM70** | Mitochondrial import receptor subunit TOM70 | **-1.40** | 0.0057 | 4 |
| P03928 | **MT-ATP8** | ATP synthase protein 8 | **-1.38** | 0.0401 | 1 |
| Q9Y5A9 | **YTHDF2** | YTH domain-containing family protein 2 | **-1.36** | 0.0140 | 1 |
| P24539 | **ATP5PB** | ATP synthase F(0) complex subunit B1, mitochondrial | **-1.35** | 0.0219 | 4 |
| Q8N5G0 | **SMIM20** | Small integral membrane protein 20 | **-1.35** | 0.0099 | 4 |
| P56182 | **RRP1** | Ribosomal RNA processing protein 1 homolog A | **-1.34** | 0.0045 | 4 |
| O95573 | **ACSL3** | Long-chain-fatty-acid--CoA ligase 3 | **-1.32** | 0.0092 | 1 |
| Q6ZNB6 | **NFXL1** | NF-X1-type zinc finger protein NFXL1 | **-1.32** | 0.0080 | 1 |
| B4DR61 | **SEC61A1** | Protein transport protein Sec61 subunit alpha isoform 1 | **-1.32** | 0.0169 | 1 |
| P35610 | **SOAT1** | Sterol O-acyltransferase 1 | **-1.30** | 0.0051 | 4 |
| P42167 | **TMPO** | Lamina-associated polypeptide 2, isoforms beta/gamma | **-1.30** | 0.0139 | 3 |
| P28331 | **NDUFS1** | NADH-ubiquinone oxidoreductase 75 kDa subunit, mitochondrial | **-1.30** | 0.0249 | 4 |
| Q9NP73 | **ALG13** | Putative bifunctional UDP-N-acetylglucosamine transferase and deubiquitinase ALG13 | **-1.30** | 0.0136 | 4 |
| P61026 | **RAB10** | Ras-related protein Rab-10 | **-1.29** | 0.0048 | 1 |
| O94874 | **UFL1** | E3 UFM1-protein ligase 1 | **-1.29** | 0.0020 | 4 |
| Q9BT22 | **ALG1** | Chitobiosyldiphosphodolichol beta-mannosyltransferase | **-1.28** | 0.0033 | 1,4 |
| Q8TC12 | **RDH11** | Retinol dehydrogenase 11 | **-1.27** | 0.0182 | 1 |
| Q8TCJ2 | **STT3B** | Dolichyl-diphosphooligosaccharide--protein glycosyltransferase subunit | **-1.27** | 0.0100 | 1,4 |
| Q9NVH1 | **DNAJC11** | DnaJ homolog subfamily C member 11 | **-1.27** | 0.0179 | 4 |
| H0YI09 | **TMT1A** | Thiol Methyltransferase 1A | **-1.27** | 0.0484 | 4 |
| Q15154 | **PCM1** | Pericentriolar material 1 protein | **-1.25** | 0.0143 | 1 |
| P60763 | **RAC3** | Ras-related C3 botulinum toxin substrate 3 | **-1.24** | 0.0500 | 5 |
| P63000 | **RAC1** | Ras-related C3 botulinum toxin substrate 1 | **-1.24** | 0.0253 | 1,3,4 |

| | | | | | |
|---|---|---|---|---|---|
| O94905 | **ERLIN2** | Erlin-2 | **-1.24** | 0.0363 | 1 |
| P49755 | **TMED10** | Transmembrane emp24 domain-containing protein 10 | **-1.24** | 0.0396 | 1 |
| O75396 | **SEC22B** | Vesicle-trafficking protein SEC22b | **-1.24** | 0.0062 | 1 |
| Q99567 | **NUP88** | Nuclear pore complex protein Nup88 | **-1.23** | 0.0022 | 3 |
| P49792 | **RANBP2** | E3 SUMO-protein ligase RanBP2 | **-1.23** | 0.0059 | 3 |
| O00592 | **PODXL** | Podocalyxin | **-1.23** | 0.0072 | 1 |
| Q9BWL3 | **C1orf43** | Uncharacterized protein C1orf43 | **-1.23** | 0.0135 | 4 |
| Q9NX20 | **MRPL16** | 39S ribosomal protein L16, mitochondrial | **-1.22** | 0.0286 | 4 |
| Q96CW1 | **AP2M1** | AP-2 complex subunit mu | **-1.22** | 0.0227 | 4 |
| Q96EY7 | **PTCD3** | Pentatricopeptide repeat domain-containing protein 3, mitochondrial | **-1.22** | 0.0193 | 4 |
| O15027 | **SEC16A** | Protein transport protein Sec16A | **-1.22** | 0.0004 | 3 |
| Q9Y4P3 | **TBL2** | Transducin beta-like protein 2 | **-1.21** | 0.0154 | 4 |
| Q9Y5M8 | **SRPRB** | Signal recognition particle receptor subunit beta | **-1.20** | 0.0163 | 1 |
| Q9NZ01 | **TECR** | Very-long-chain enoyl-CoA reductase | **-1.20** | 0.0093 | 1 |
| Q7Z7H5 | **TMED4** | Transmembrane emp24 domain-containing protein 4 | **-1.20** | 0.0453 | 1 |
| Q9H0U4 | **RAB1B** | Ras-related protein Rab-1B | **-1.20** | 0.0476 | 4 |
| Q9H0P0 | **NT5C3A** | Cytosolic 5'-nucleotidase 3A | **-1.20** | 0.0127 | 4 |
| Q9Y2U8 | **LEMD3** | Inner nuclear membrane protein Man1 | **-1.20** | 0.0015 | 4 |
| P51153 | **RAB13** | Ras-related protein Rab-13 | **-1.20** | 0.0364 | 1 |
| O94973 | **AP2A2** | AP-2 complex subunit alpha-2 | **-1.20** | 0.0002 | 4 |

FC, fold change. |FC| > 1.2, $p<0.05$.

**Supplementary Table 11**: Pearson's correlation analyses of markers for retinal inflammasome, cell degeneration, and gliosis with retinal AD pathology markers

<table>
<tr><th></th><th></th><th colspan="3">Retinal inflammasome</th><th colspan="2">Retinal degeneration</th><th colspan="3">Retinal gliosis</th></tr>
<tr><th></th><th></th><th>NLRP3</th><th>ASC</th><th>Caspase-1</th><th>NGSDMD</th><th>CCasp3</th><th>Iba1</th><th>GFAP</th><th>Vimentin</th></tr>
<tr><td rowspan="2">Retinal Amyloidosis</td><td>$A\beta_{42}$</td><td>0.81****<br>n = 19</td><td>0.40<br>n = 19</td><td>0.70***<br>n = 19</td><td>0.64**<br>n = 19</td><td>0.77***<br>n = 17</td><td>0.85****<br>n = 20</td><td>0.65**<br>n = 20</td><td>0.56*<br>n = 15</td></tr>
<tr><td>Aβ oligomer</td><td>0.41<br>n = 9</td><td>-0.40<br>n = 9</td><td>0.39<br>n = 9</td><td>-0.22<br>n = 8</td><td>0.32<br>n = 7</td><td>0.46<br>n = 9</td><td>-0.15<br>n = 10</td><td>0.05<br>n = 8</td></tr>
<tr><td rowspan="4">Retinal Tauopathy</td><td>PHF-1</td><td>0.30<br>n = 13</td><td>0.31<br>n = 13</td><td>0.19<br>n = 13</td><td>0.28<br>n = 13</td><td>0.30<br>n = 12</td><td>0.18<br>n = 15</td><td>0.60*<br>n = 13</td><td>0.21<br>n = 7</td></tr>
<tr><td>pS396</td><td>0.40*<br>n = 25</td><td>0.44*<br>n = 25</td><td>0.32<br>n = 25</td><td>0.38<br>n = 24</td><td>0.53**<br>n = 25</td><td>0.31<br>n = 27</td><td>0.43*<br>n = 24</td><td>0.19<br>n = 12</td></tr>
<tr><td>MC-1</td><td>0.23<br>n = 24</td><td>-0.21<br>n = 24</td><td>-0.12<br>n = 24</td><td>-0.12<br>n = 20</td><td>0.06<br>n = 21</td><td>0.26<br>n = 26</td><td>0.14<br>n = 24</td><td>-0.06<br>n = 13</td></tr>
<tr><td>Oligo-tau</td><td>0.70***<br>n = 24</td><td>0.44*<br>n = 24</td><td>0.60**<br>n = 24</td><td>0.77****<br>n = 23</td><td>0.80****<br>n = 23</td><td>0.69****<br>n = 27</td><td>0.60**<br>n = 24</td><td>0.60*<br>n = 14</td></tr>
<tr><td rowspan="3">Retinal Gliosis</td><td>Iba1</td><td>0.77****<br>n = 25</td><td>0.42*<br>n = 25</td><td>0.65***<br>n = 25</td><td>0.39<br>n = 24</td><td>0.69***<br>n = 23</td><td>-</td><td>0.65***<br>n = 24</td><td>0.81**<br>n = 9</td></tr>
<tr><td>GFAP</td><td>0.91****<br>n = 23</td><td>0.71***<br>n = 23</td><td>0.77****<br>n = 23</td><td>0.68***<br>n = 26</td><td>0.85****<br>n = 21</td><td>0.65***<br>n = 24</td><td>-</td><td>0.42<br>n = 11</td></tr>
<tr><td>Vimentin</td><td>0.84**<br>n=10</td><td>0.16<br>n = 10</td><td>0.78**<br>n = 10</td><td>0.54<br>n = 11</td><td>0.58<br>n = 8</td><td>0.81**<br>n = 9</td><td>0.42<br>n = 11</td><td>-</td></tr>
<tr><td rowspan="3">Retinal Inflammasome</td><td>NLRP3</td><td>-</td><td>0.59**<br>n = 27</td><td>0.83****<br>n = 27</td><td>0.74****<br>n = 23</td><td>0.80****<br>n = 23</td><td>0.77****<br>n = 25</td><td>0.91****<br>n = 23</td><td>0.84**<br>n = 10</td></tr>
<tr><td>ASC</td><td>0.59**<br>n = 27</td><td>-</td><td>0.70****<br>n = 27</td><td>0.58**<br>n = 23</td><td>0.76****<br>n = 23</td><td>0.42*<br>n = 25</td><td>0.71***<br>n = 23</td><td>0.16<br>n = 10</td></tr>
<tr><td>Caspase-1</td><td>0.83****<br>n = 27</td><td>0.70****<br>n = 27</td><td>-</td><td>0.57**<br>n = 23</td><td>0.80****<br>n = 23</td><td>0.65***<br>n = 25</td><td>0.77****<br>n = 23</td><td>0.78**<br>n = 10</td></tr>
<tr><td rowspan="3">Retinal Degeneration</td><td>NGSDMD</td><td>0.74****<br>n = 23</td><td>0.58**<br>n = 23</td><td>0.57**<br>n = 23</td><td>-</td><td>0.75****<br>n = 21</td><td>0.39<br>n = 24</td><td>0.68***<br>n = 26</td><td>0.54<br>n = 11</td></tr>
<tr><td>CCasp3</td><td>0.80****<br>n = 23</td><td>0.76****<br>n = 23</td><td>0.80****<br>n = 23</td><td>0.75****<br>n = 21</td><td>-</td><td>0.69***<br>n = 23</td><td>0.85****<br>n = 21</td><td>0.58<br>n = 8</td></tr>
<tr><td>Atrophy</td><td>0.86****<br>n = 14</td><td>0.62*<br>n = 14</td><td>0.82***<br>n=14</td><td>0.60*<br>n = 14</td><td>0.86***<br>n = 11</td><td>0.72**<br>n = 15</td><td>0.72**<br>n = 15</td><td>0.87*<br>n = 6</td></tr>
</table>

Pearson's correlation analyses: *p* and *r* values determine the statistical significance and strength of each pairwise association between markers of retinal inflammasome, degeneration, and gliosis versus retinal AD-associated pathologies, inflammasomes, gliosis, and degeneration markers. *p* and *r* values presented in bold type with asterisk(s) depicts strong to very strong correlation.

**Supplementary Table 12**: Spearman's correlation analyses of markers for retinal inflammasome, cell degeneration, and gliosis with brain AD pathology and cognition

| | **Retinal inflammasome** | | | **Retinal degeneration** | | **Retinal gliosis** | | |
|---|---|---|---|---|---|---|---|---|
| | NLRP3 | ASC | Caspase-1 | NGSDMD | CCasp3 | Iba1 | GFAP | Vimentin |
| **Brain Aβ plaque** | 0.34<br>n = 27 | 0.25<br>n = 27 | 0.34<br>n = 27 | 0.21<br>n = 25 | 0.42*<br>n = 25 | 0.28<br>n = 29 | 0.39*<br>n = 26 | 0.38<br>n = 18 |
| **ABC** | **0.67*****<br>**n = 27** | 0.35<br>n = 27 | 0.56**<br>n = 27 | 0.43*<br>n = 25 | **0.72******<br>**n = 25** | 0.49**<br>n = 29 | **0.71******<br>**n = 26** | **0.62****<br>**n = 18** |
| **NFTs** | **0.64*****<br>**n = 27** | 0.53**<br>n = 27 | **0.65*****<br>**n = 27** | 0.49*<br>n = 25 | **0.78******<br>**n = 25** | 0.56**<br>n = 29 | 0.59**<br>n = 26 | 0.43<br>n = 18 |
| **Braak** | **0.68*****<br>**n = 27** | **0.60*****<br>**n = 27** | **0.63****<br>**n = 27** | 0.55**<br>n = 25 | **0.72******<br>**n = 25** | 0.49**<br>n = 30 | **0.78******<br>**n = 26** | 0.46<br>n = 18 |
| **Brain atrophy** | 0.47*<br>n = 27 | 0.27<br>n = 27 | 0.41*<br>n = 27 | 0.23<br>n = 25 | 0.45*<br>n = 25 | 0.36<br>n = 29 | 0.45*<br>n = 26 | 0.16<br>n = 18 |
| **CAA** | 0.54**<br>n = 27 | 0.31<br>n = 27 | 0.47*<br>n = 27 | 0.29<br>n = 25 | 0.59**<br>n = 25 | 0.49**<br>n = 29 | 0.39<br>n = 26 | 0.35<br>n = 17 |
| **CDR** | **0.66*****<br>**n = 27** | 0.26<br>n = 27 | 0.55**<br>n = 27 | 0.37<br>n = 25 | **0.62****<br>**n = 25** | **0.66*****<br>**n = 28** | 0.51**<br>n = 26 | **0.68****<br>**n = 16** |
| **MMSE** | **-0.66*****<br>**n = 25** | -0.49*<br>n = 25 | -0.58*<br>n = 25 | -0.50*<br>n = 25 | **-0.69*****<br>**n = 22** | -0.58**<br>n = 26 | -0.48*<br>n = 26 | -0.59**<br>n = 18 |
| **MoCA** | -0.76<br>n = 7 | -0.33<br>n = 7 | -0.60<br>n = 7 | -0.41<br>n = 6 | -0.45<br>n = 8 | -0.31<br>n = 7 | -0.49<br>n = 6 | -0.80<br>n = 4 |

Spearman's rank correlation analyses: *p* and *r* values determine the statistical significance and strength of each pairwise association between markers of retinal inflammasome, degeneration, and gliosis versus brain AD-pathology markers. *p* and *r* values presented in bold type with asterisk(s) depicts strong to very strong correlation.

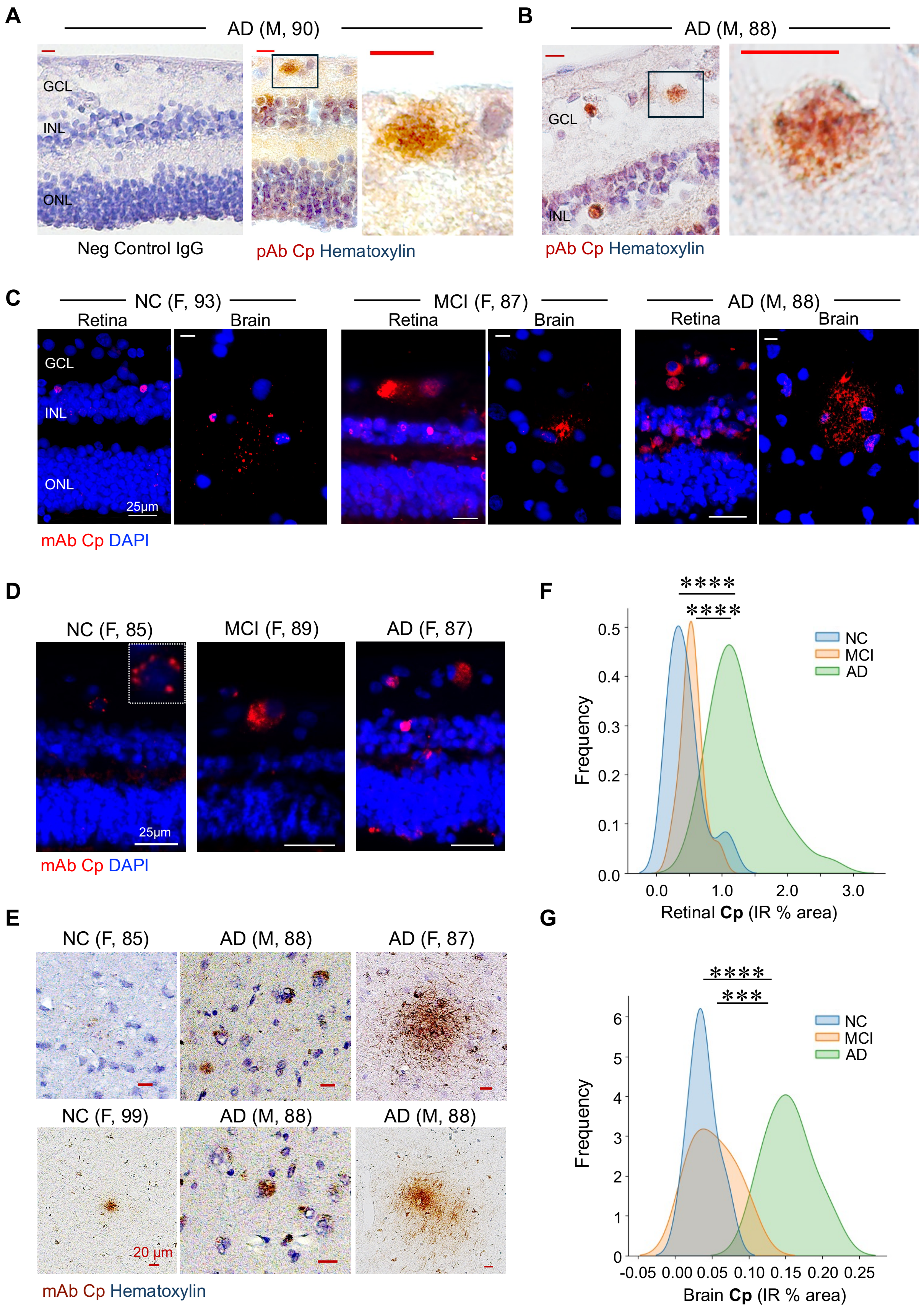
A
AD (M, 90)
GCL
INL
ONL
Neg Control IgG
pAb Cp Hematoxylin
B
AD (M, 88)
GCL
INL
pAb Cp Hematoxylin
C
NC (F, 93)
MCI (F, 87)
AD (M, 88)
Retina
Brain
25µm
mAb Cp DAPI
D
NC (F, 85)
MCI (F, 89)
AD (F, 87)
25µm
mAb Cp DAPI
E
NC (F, 85)
AD (M, 88)
AD (F, 87)
NC (F, 99)
AD (M, 88)
AD (M, 88)
20 µm
mAb Cp Hematoxylin
F
****
****
NC
MCI
AD
Frequency
Retinal Cp (IR % area)
G
****
***
NC
MCI
AD
Frequency
Brain Cp (IR % area)

**Supplementary Figure 1. Cp inclusions in AD retina and paired-brain tissues.**

**A-B.** Representative peroxidase-based images (DAB brown, hematoxylin nuclei blue) of retinal cross-sections from AD patients. **A.** Left image exhibit no immunoreaction when used negative (Neg) control IgG, whereas staining with Cp polyclonal antibody (pAb) depicts the presence of Cp inclusions in GCL and INL (middle image). Cp inclusions in higher magnification is shown (right image). **B.** AD retinal cross sections depict the presence of Cp inclusions in the GCL and INL. Higher magnification, right image. Scale bars: 10 μm. **C.** Representative fluorescence images of retinal and paired-brain (Area 9 located in the dorsolateral prefrontal cortex) cross-sections from MCI and AD patients versus NC individuals, depicting the presence of Cp inclusions (red), with specific Cp monoclonal antibody (mAb). DAPI (blue) stained nuclei. Scale bars: 25 μm (retina) and 10 μm (brain cross-sections). **D.** Representative fluorescence images of retinal cross-sections from MCI and AD patients versus NC individuals (the same subjects which were stained using peroxidase-based (DAB) method, see **Fig. 1E)**, depicting the presence of Cp inclusions (red), stained with specific Cp mAb. DAPI (blue) stained nuclei. Scale bars: 25 μm. **E.** Representative peroxidase-based (DAB) images of brain cross-sections from AD patients versus NC individuals depicting the presence of Cp inclusions (brown) stained with Cp mAb and hematoxylin. Scale bars: 20 μm**. F-G.** Gaussian distribution curves display the frequency of **(F)** Retinal and **(G)** Brain Cp % IR area in individuals with premortem clinical diagnoses of NC, MCI, and AD. ***$p < 0.001$ and ****$p < 0.0001$, by one-way ANOVA and Tukey's post hoc multiple comparison test. Ganglion cell layer (GCL); Inner nuclear layer (INL); Outer nuclear layer (ONL).

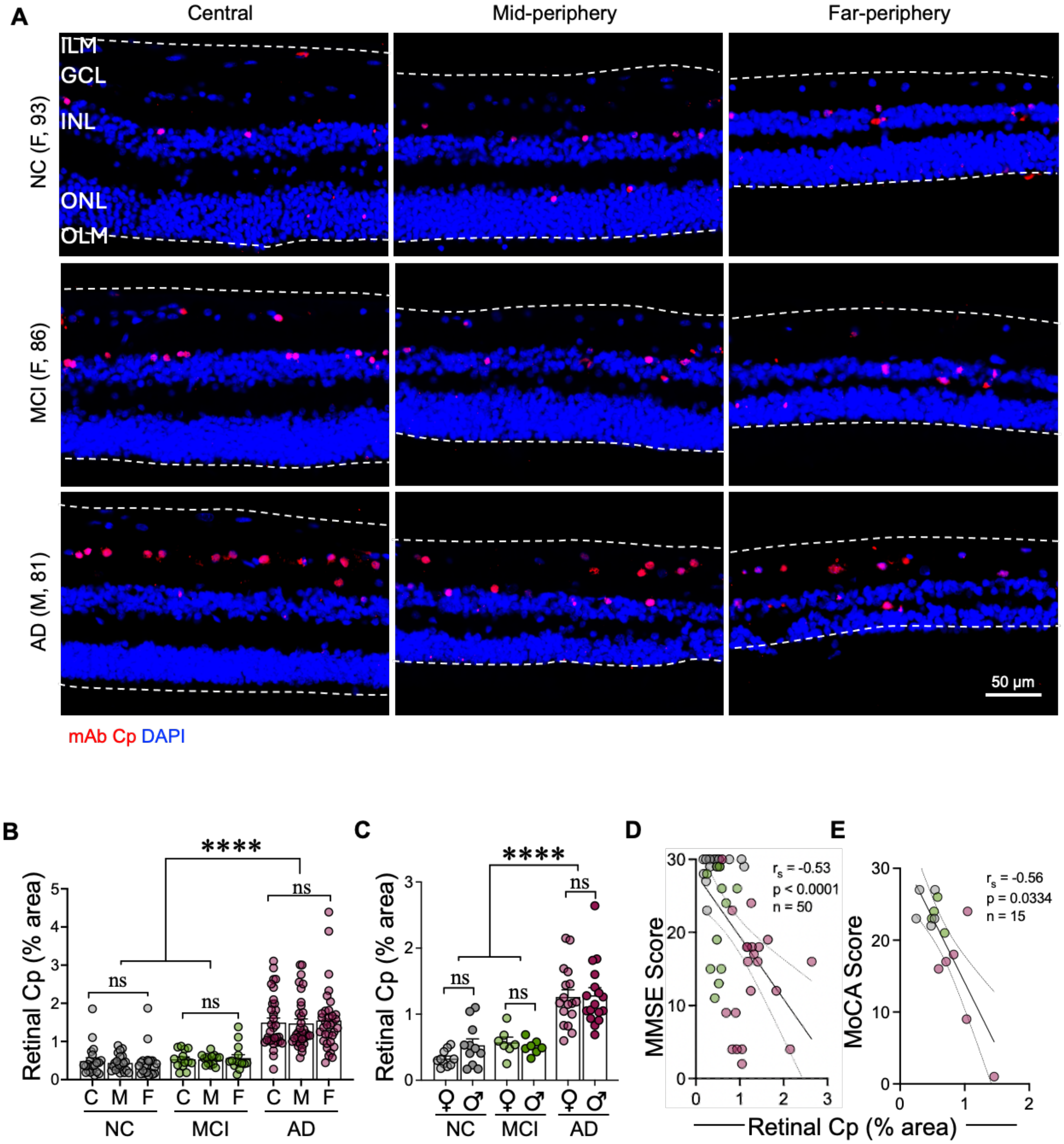
A
Central
Mid-periphery
Far-periphery
ILM
GCL
INL
ONL
OLM
NC (F, 93)
MCI (F, 86)
AD (M, 81)
50 µm
mAb Cp DAPI
B
Retinal Cp (% area)
****
ns
C M F
NC
MCI
AD
C
D
MMSE Score
r<sub>s</sub> = -0.53
p < 0.0001
n = 50
E
MoCA Score
r<sub>s</sub> = -0.56
p = 0.0334
n = 15
Retinal Cp (% area)

**Supplementary Figure 2. Distribution of Cp inclusions across retinal subregions and sex.**

**A.** Representative fluorescence images of retinal cross-sections from MCI and AD patients versus NC individuals depicting the distribution of Cp inclusions (red) across three retinal subregions (central, mid-periphery, and far-periphery). DAPI (blue) stained nuclei. White dotted lines display the analyzed area, between the inner limiting membrane (ILM) and outer limiting membrane (OLM). Scale bar: 50 µm. **B-C.** Scatter plots display quantitative-IHC analysis of retinal Cp % IR area in (**B**) three retinal subregions (C - Central; M - Mid periphery and F - Far periphery) among NC (N=21), MCI (N=14), and AD (N=34) subjects, and (**C**) male and female subjects of same cohort (NC=10M, 11F, MCI=7M, 7F, AD=17M, 17F). **D, E.** Pearson correlation ($r_p$) analyses between retinal Cp burden and (**D**) MMSE, and (**E**) MoCA cognitive score. Statistics: Data from individual subjects (circles) as well as group means ± SEMs are shown. ♀ = female; ♂ = Male. ****$p < 0.0001$, by one-way ANOVA and Tukey's post hoc multiple comparison test. ns = non-significant. Ganglion cell layer (GCL); Inner nuclear layer (INL); Outer nuclear layer (ONL).

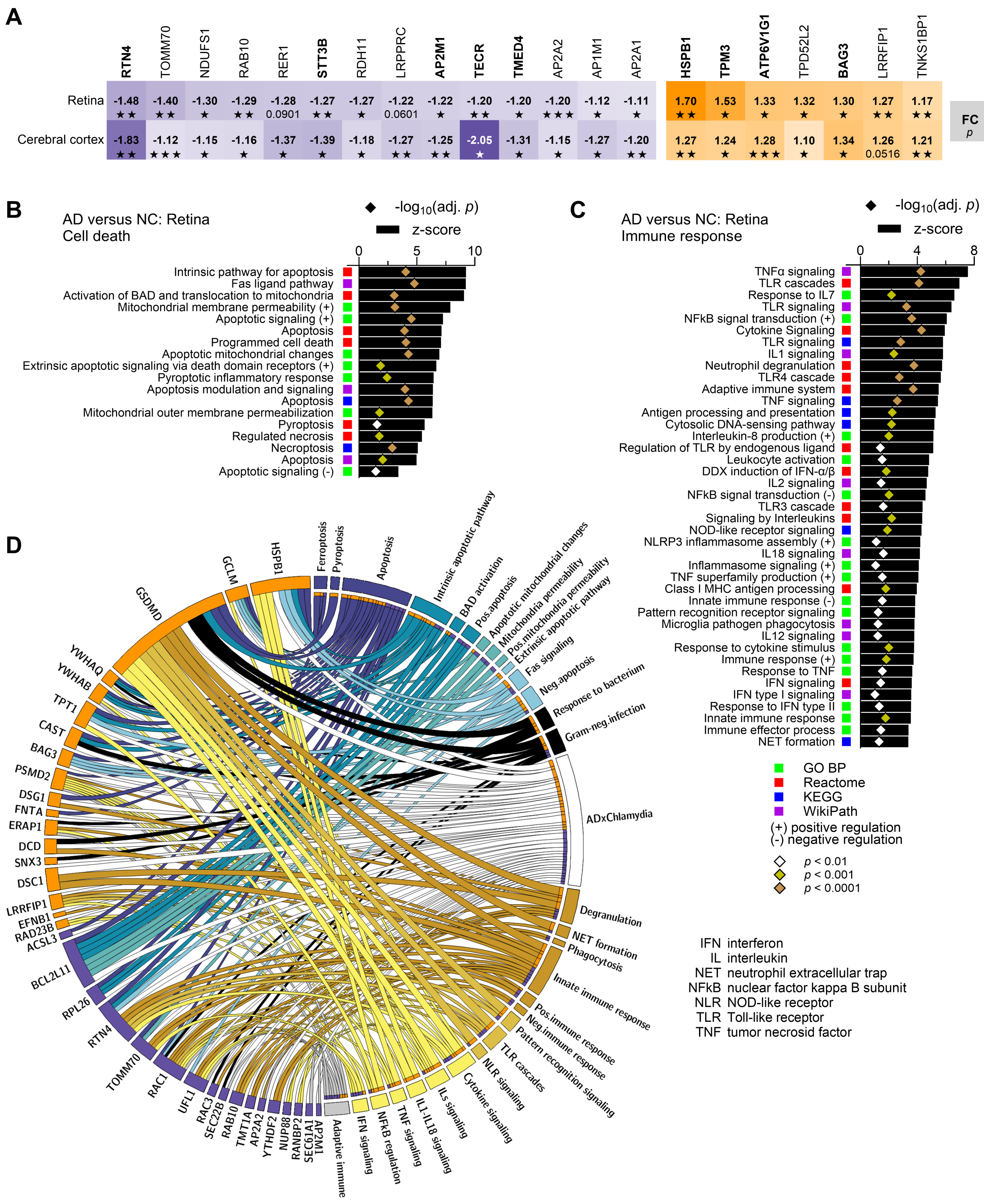


**Supplementary Figure 3. Cell death and immune response pathways, and association with *Chlamydia* infection in the human retina and cerebral cortex.**

**A.** Heatmaps of shared *Chlamydia*-interactome differentially expressed proteins (DEPs) in AD retina and cerebral cortex [Fold Change (FC) and *p*]. The expression of 5 downregulated (RTN4, STT3B, AP2M1, TECR, TMED4) and 5 upregulated (HSPB1, TPM3, LRRFIP1 [t-test], BAG3, ATP6V1G1) proteins were within cutoffs for DEPs ($|FC| > 1.2$ and $p<0.05$) in both the retina and the cortex. Additional proteins, whose expression was within cutoffs for DEPs ($|FC| > 1.2$ and $p<0.05$) for either the retina or the cortex, were found. **B-C.** Gene ontology (GO) analysis of DEPs related to (**B)** cell death and (**C**) immune response in human AD (n = 6) versus NC retina (n = 6). The analysis was carried out in Metascape and included the GO Biological Processes (BP), Reactome, Kyoto Encyclopedia of Genes and Genomes (KEGG) and WikiPathways databases. Bar and symbol graphs represent z-scores and Benjamini-Hochberg adjusted *p*-values from Metascape analysis, respectively. Range of *p*-values are presented as color-coded symbols. **D.** Chord diagram displays the association of *Chlamydia*-interacting proteins with pathways related to cell death (blue gradient outer segments and ribbons), immune response (yellow gradient and grey outer segments and ribbons), as well as Gram (-) bacterial infection (white outer segment and ribbons). For proteins, orange and purple outer segments indicates upregulation and downregulation, respectively.

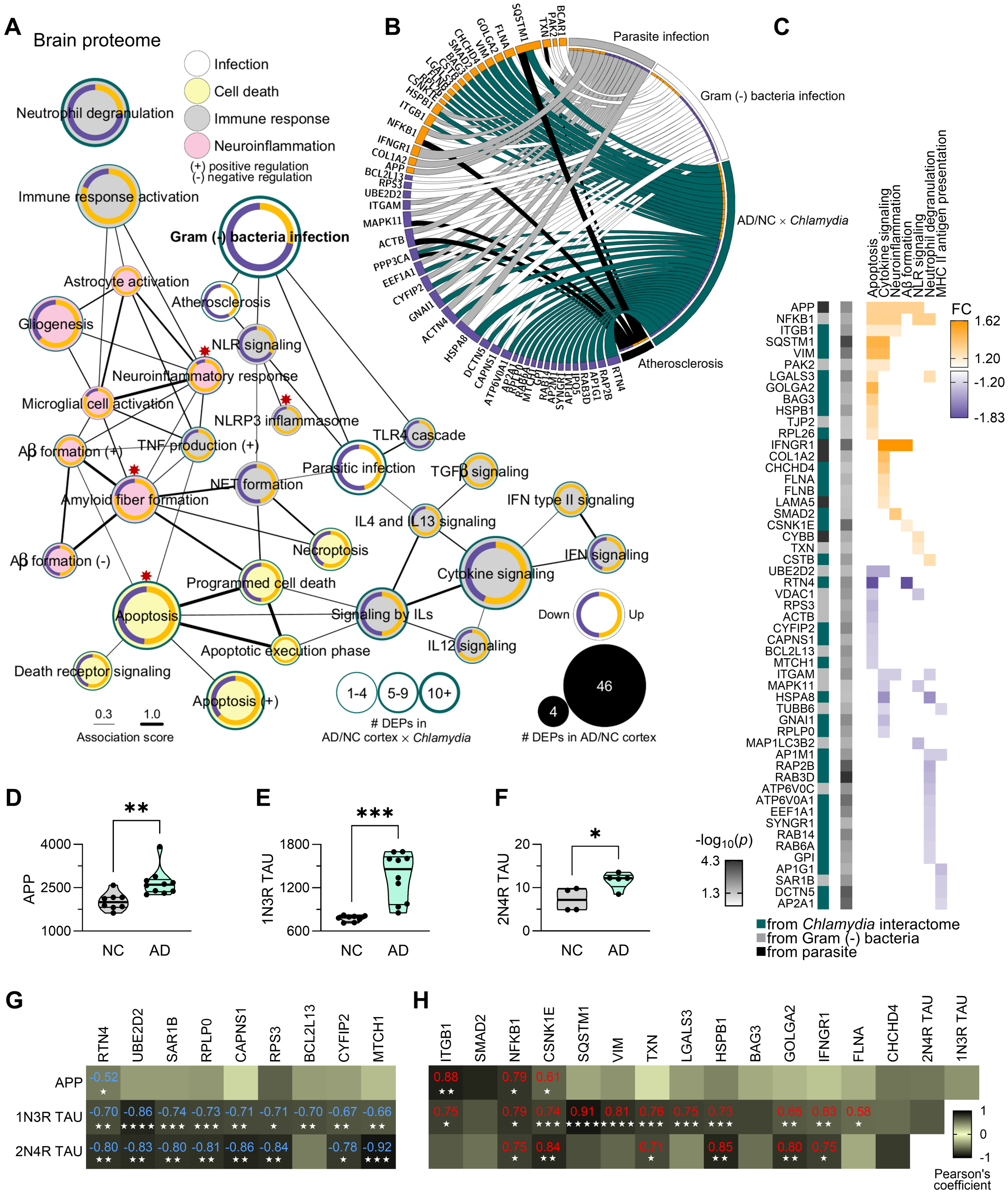
A
Brain proteome
Infection
Cell death
Immune response
Neuroinflammation
(+) positive regulation
(-) negative regulation
Neutrophil degranulation
Immune response activation
Gram (-) bacteria infection
Astrocyte activation
Atherosclerosis
Gliogenesis
NLR signaling
Neuroinflammatory response
Microglial cell activation
NLRP3 inflammasome
TLR4 cascade
Aβ formation (+)
TNF production (+)
Parasitic infection
TGFβ signaling
NET formation
Amyloid fiber formation
IFN type II signaling
IL4 and IL13 signaling
Necroptosis
IFN signaling
Aβ formation (-)
Cytokine signaling
Programmed cell death
Apoptosis
Signaling by ILs
Down
Up
Apoptotic execution phase
IL12 signaling
Death receptor signaling
Apoptosis (+)
0.3
1.0
Association score
1-4
5-9
10+
### DEPs in AD/NC cortex × Chlamydia
46
4
### DEPs in AD/NC cortex
B
Parasite infection
Gram (-) bacteria infection
AD/NC × Chlamydia
Atherosclerosis
C
FC
1.62
1.20
-1.20
-1.83
Apoptosis
Cytokine signaling
Neuroinflammation
Aβ formation
NLR signaling
Neutrophil degranulation
MHC II antigen presentation
APP
NFKB1
ITGB1
SQSTM1
VIM
PAK2
LGALS3
GOLGA2
BAG3
HSPB1
TJP2
RPL26
IFNGR1
COL1A2
CHCHD4
FLNA
FLNB
LAMA5
SMAD2
CSNK1E
CYBB
TXN
CSTB
UBE2D2
RTN4
VDAC1
RPS3
ACTB
CYFIP2
CAPNS1
BCL2L13
MTCH1
ITGAM
MAPK11
HSPA8
TUBB6
GNAI1
RPLP0
MAP1LC3B2
AP1M1
RAP2B
RAB3D
ATP6V0C
ATP6V0A1
EEF1A1
SYNGR1
RAB14
RAB6A
GPI
AP1G1
SAR1B
DCTN5
AP2A1
-log10(p)
4.3
1.3
from Chlamydia interactome
from Gram (-) bacteria
from parasite
D
APP
4000
2500
1000
NC
AD
**
E
1N3R TAU
1800
1200
600
NC
AD
***
F
2N4R TAU
20
10
0
NC
AD
*
G
RTN4
UBE2D2
SAR1B
RPLP0
CAPNS1
RPS3
BCL2L13
CYFIP2
MTCH1
APP
-0.52
1N3R TAU
-0.70 -0.86 -0.74 -0.73 -0.71 -0.71 -0.70 -0.67 -0.66
2N4R TAU
-0.80 -0.83 -0.80 -0.81 -0.86 -0.84 -0.78 -0.92
H
ITGB1
SMAD2
NFKB1
CSNK1E
SQSTM1
VIM
TXN
LGALS3
HSPB1
BAG3
GOLGA2
IFNGR1
FLNA
CHCHD4
2N4R TAU
1N3R TAU
0.88 0.79 0.61
0.75 0.79 0.74 0.91 0.81 0.76 0.75 0.73 0.65 0.63 0.58
0.75 0.84 0.71 0.85 0.80 0.75
1
0
-1
Pearson's coefficient

**Supplementary Figure 4. Cell death and immune response pathways, and association with *Chlamydia* infection in the cerebral cortex.**

**A.** GO network (Metascape) of pathways related to infection, cell death, immune response and neuroinflammation. The size of the nodes represents the number of DEPs in AD versus NC cortex, with the inner ring showing the proportion of these DEPs that are downregulated (purple) or upregulated (orange) in AD. The green border and its thickness represent the number of DEPs that interact with *Chlamydia* inclusion in each pathway. The thickness of edges represents the shared DEPs (association score) between pathways. Red asterisks indicate pathways (NLRP3 inflammasome, apoptosis, neuroinflammatory response, and amyloid fiber formation) that were further explored and validated. **B.** Chord diagram displays the association between cell death, immune response-related DEPs in AD cortex and *Chlamydia* (interactome), Gram (-) bacteria and parasitic infections, and atherosclerosis. For DEPs, orange and purple outer segments indicate upregulation and downregulation, respectively. **C.** Heatmaps of DEPs [FC and -$\log_{10}(p)$] in AD cortex for selected pathways. Only proteins connected to Gram (-) bacteria and parasitic infections (Metascape analysis) and *Chlamydia* infection (*Chlamydia* interactome) are shown for each pathway. **D-F.** Mass spectrometry (MS) quantitation of (**D)** Amyloid beta precursor protein (APP), (**E**) one N-terminal domain and three microtubule-binding repeat domains (1N3R) and **(F)** two N-terminal domains and four microtubule-binding repeat domains (2N4R) isoforms of TAU in cerebral cortex of human AD (n = 10) and NC retina (n = 8). **G-H.** Heatmaps of (**G)** negative and (**H**) positive Pearson's correlation coefficients analysis (*r*) between *Chlamydia* interactors and APP, 1N3R TAU, and 2N4R TAU, quantified by MS. Correlation statistics: *$p$<0.05, **$p$<0.01, ***$p$<0.001, and ****$p$<0.0001.

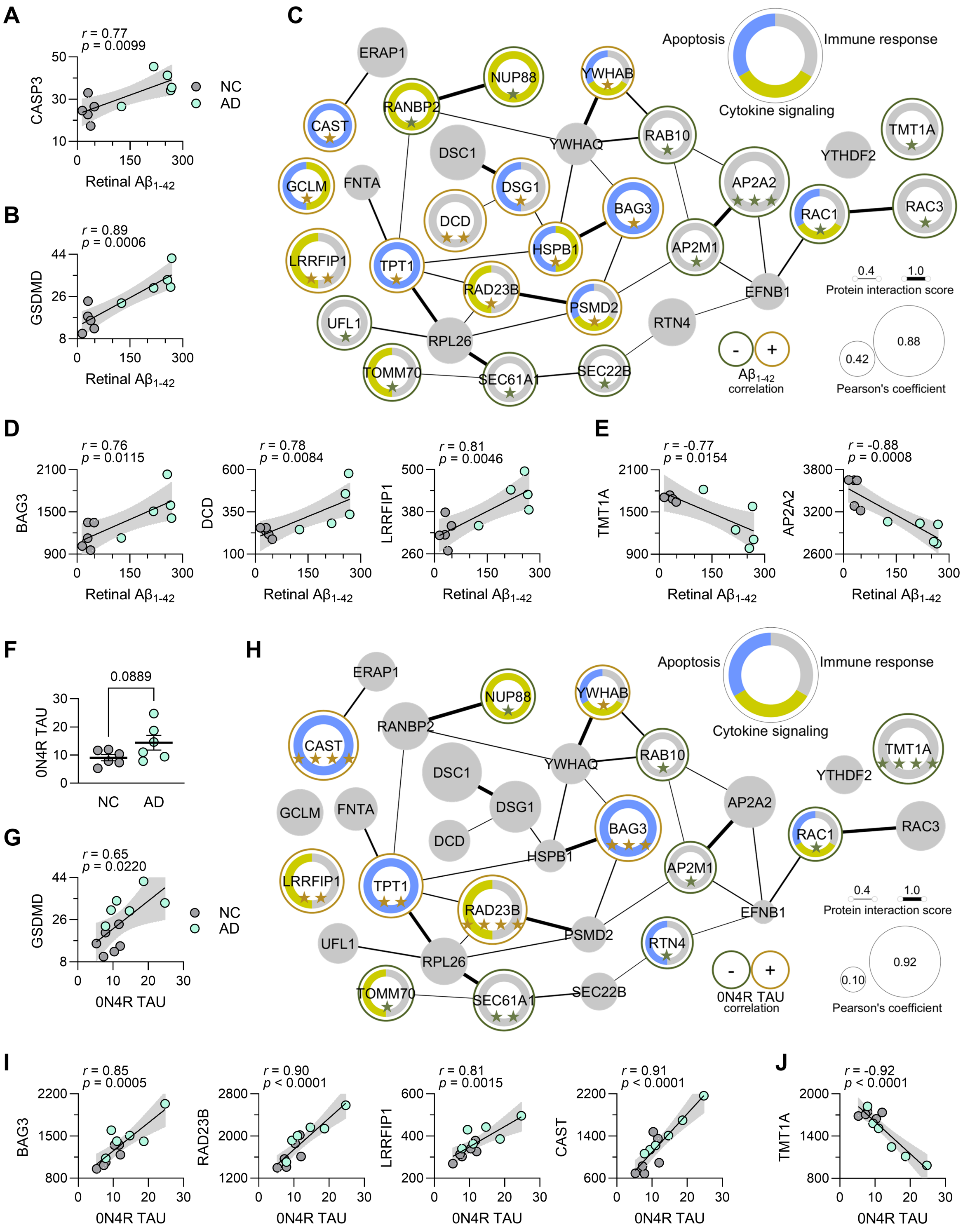

**Supplementary Figure 5. Correlation of *Chlamydia* inclusion interactors with amyloid deposits and neurofibrillary tangle burden in the AD retina.**

**A-B.** Pearson's correlation coefficient analysis of A$\beta_{1\text{-}42}$ levels with (**A**) Caspase-3 (CASP3) and (**B**) Gasdermin D (GSDMD). Retinal A$\beta_{1\text{-}42}$ was quantified by using sandwich enzyme-linked immunosorbent assay (ELISA) from human AD (n = 5) and NC retina (n = 5). (**C**) Protein interaction network (String v12.0) of *Chlamydia* interactors and their correlation with A$\beta_{1\text{-}42}$ levels. Node size represents the Pearson's correlation coefficient (*r*). Dark green and orange node borders indicate negative and positive correlation with A$\beta_{1\text{-}42}$ levels, respectively. Inner ring categorizes the protein's role into apoptosis (blue), immune response (grey) and/or cytokine signaling (olive). The thickness of edges between nodes represents the protein interaction score from String v12.0. Correlation statistics: ⋆$p$<0.05, ⋆⋆$p$<0.01, and ⋆⋆⋆$p$<0.001. **D-E.** Pearson's correlation coefficients analysis between several *Chlamydia* interactors and retinal A$\beta_{1\text{-}42}$ levels. Interactors with strongest (**D**) positive and (**E**) negative correlations are shown. **F.** Mass spectrometry quantitation of 0N4R isoform of TAU [absence of N-terminal insert (0N) and presence of four microtubule-binding repeat domains (4R)] in human AD (n = 6) and NC retina (n = 6). **G.** Pearson's correlation coefficient analysis of 0N4R TAU with GSDMD. **H.** The same protein interaction network of *Chlamydia* interactors showing their correlation to 0N4R TAU. Node size represents the Pearson's correlation coefficient (*r*). Dark green and orange node borders indicate negative and positive correlation with 0N4R TAU, respectively. Inner ring categorizes the protein's role in apoptosis (blue), immune response (grey), and/or cytokine signaling (olive). The thickness of edges between nodes represents the protein interaction score from String v12.0. Correlation statistics: ⋆$p$<0.05, ⋆⋆$p$<0.01, ⋆⋆⋆$p$<0.001, and ⋆⋆⋆⋆$p$<0.0001. **I-J.** Pearson's correlation coefficients analysis between several *Chlamydia* interactors and 0N4R TAU quantitated in the retina by MS. Interactors with strongest (**I**) positive and (**J**) negative correlations are shown.

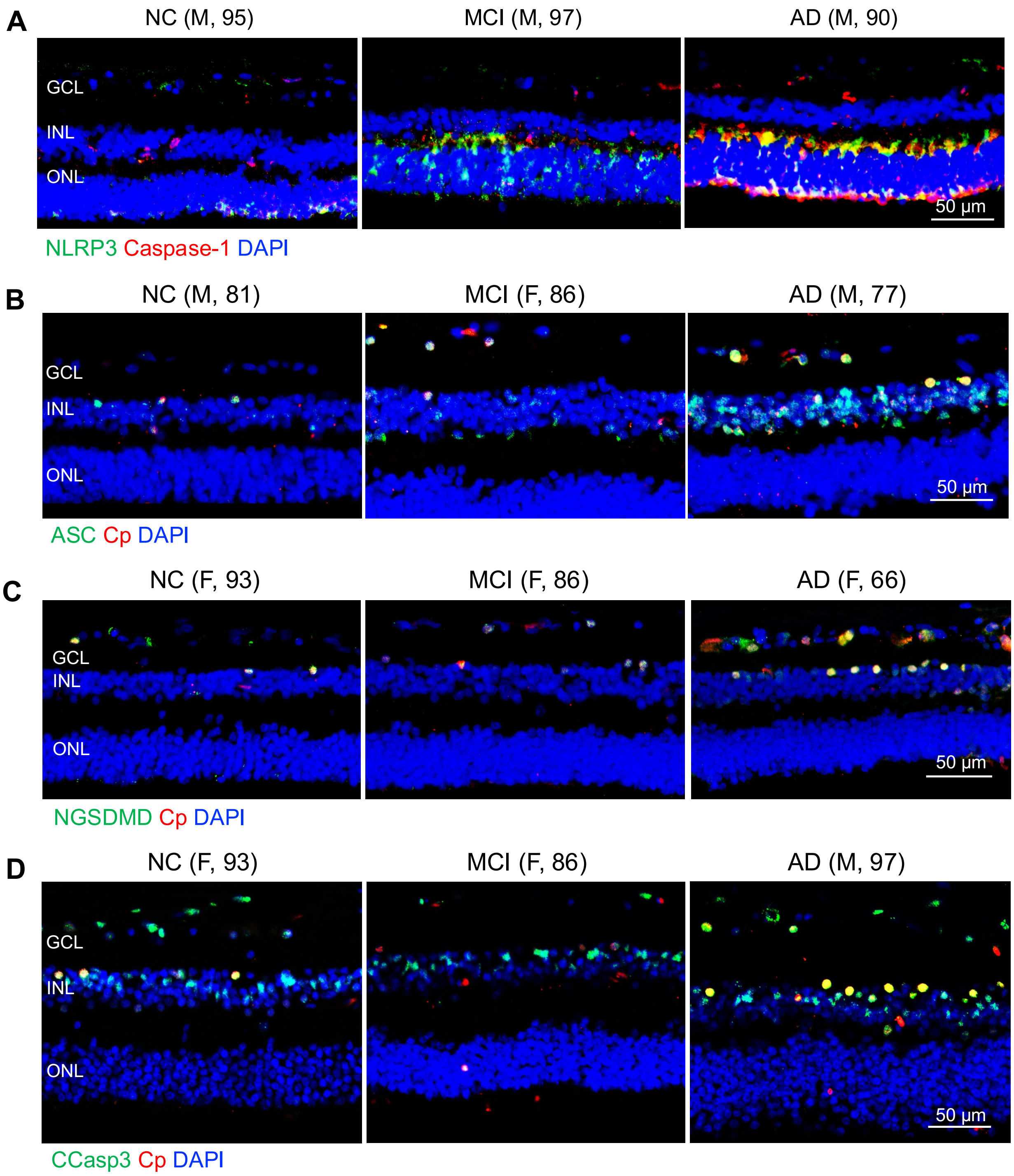
A
NC (M, 95)
MCI (M, 97)
AD (M, 90)
GCL
INL
ONL
50 µm
NLRP3 Caspase-1 DAPI
B
NC (M, 81)
MCI (F, 86)
AD (M, 77)
GCL
INL
ONL
50 µm
ASC Cp DAPI
C
NC (F, 93)
MCI (F, 86)
AD (F, 66)
GCL
INL
ONL
50 µm
NGSDMD Cp DAPI
D
NC (F, 93)
MCI (F, 86)
AD (M, 97)
GCL
INL
ONL
50 µm
CCasp3 Cp DAPI

**Supplementary Figure 6. Retinal Cp Colocalized with retinal NLRP3 inflammasome components, early apoptosis, and cellular pyroptosis markers.**

**A-D.** Representative images of retinal cross-sections from MCI and AD patients versus NC controls stained with **(A)** NLRP3 (green) and caspase-1, (red), (**B**) ASC (green) and Cp (red), **(C)** NGSDMD (green) and Cp (red), (**D**) CCasp3 (green) and Cp (red) with DAPI (blue) nuclear staining. Colocalization are shown in yellow. Scale bars: 50 µm.

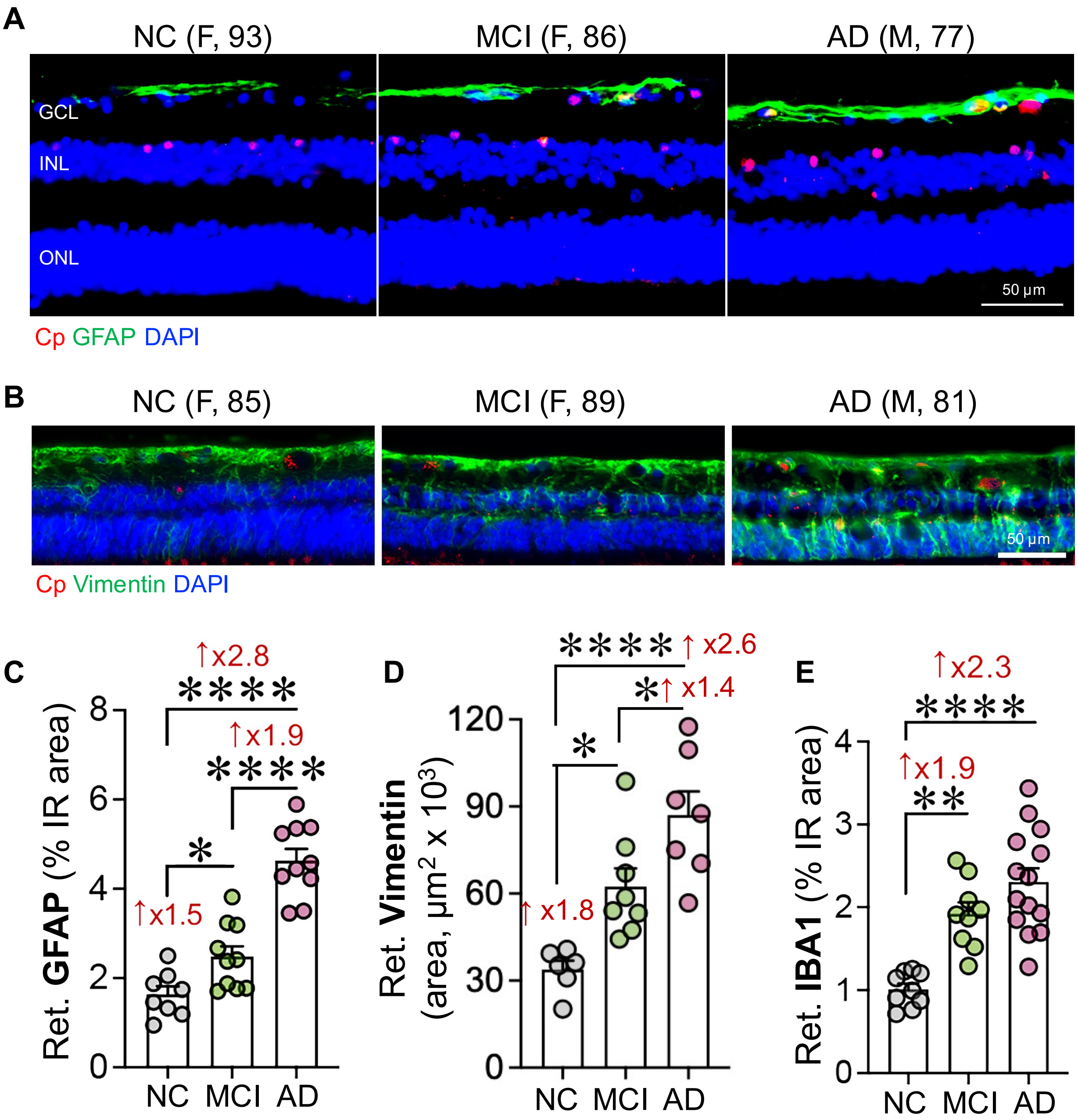


**Supplementary Figure 7. Association of retinal Cp with retinal gliosis.**

**A-B.** Representative images of retinal cross-sections from MCI and AD patients versus NC controls stained with (**A**) Cp (red) and GFAP (green) and (**B**) Cp (red) and vimentin (green) with DAPI (blue) nuclear staining. **C-E.** Quantitative IHC analysis of retinal macrogliosis (**C**) astroglia (GFAP) and (**D**) Müller Glia (vimentin), (**E**) microglia (IBA1) % IR area in individuals with premortem clinical diagnoses of NC (n = 6-9), MCI (n = 8-10), and AD (n = 7-14). Scale bar: 50 µm. Statistics: Data from individual subjects (circles) as well as group means ± SEMs are shown. Fold changes are shown in red. **p* < 0.05, ***p* < 0.01, and *****p* < 0.0001, by one-way ANOVA and Tukey's post hoc multiple comparison test.

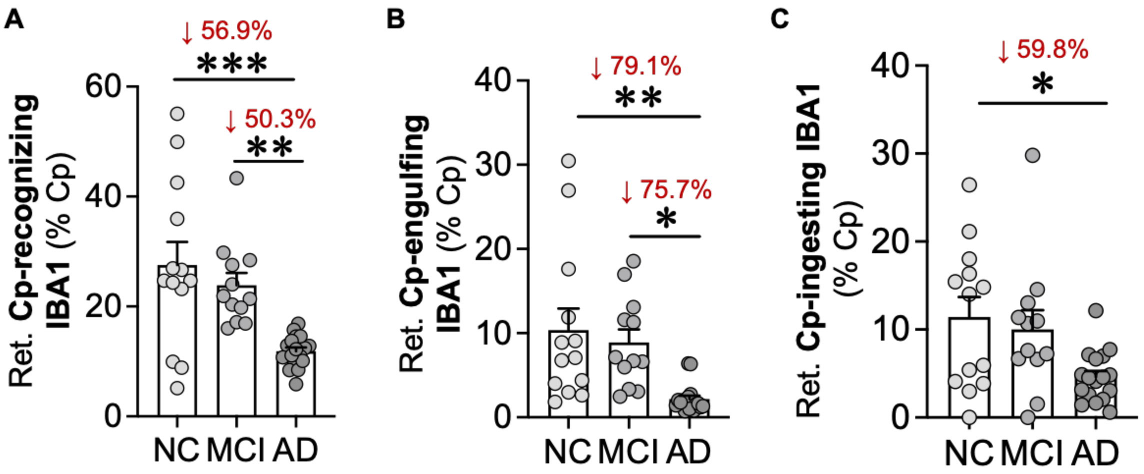


**Supplementary Figure 8. Analysis of retinal microglia in association with retinal Cp.**

**A-C.** Scatter plot of quantitative IHC analysis of retinal microgliosis (IBA1) per retinal Cp % IR area in three microglial stages for Cp phagocytosis (**A**) Cp-recognizing microglia, (**B**) Cp-engulfing microglia, and (**C**) Cp-ingesting microglia in individuals with premortem clinical diagnoses of NC (n = 13), MCI (n = 12), and AD (n = 17). Statistics: Data from individual subjects (circles) as well as group means ± SEMs are shown. Percentage (%) changes are shown in red. *$p < 0.05$, **$p < 0.01$, and ***$p < 0.001$, by one-way ANOVA and Tukey's post hoc multiple comparison test.

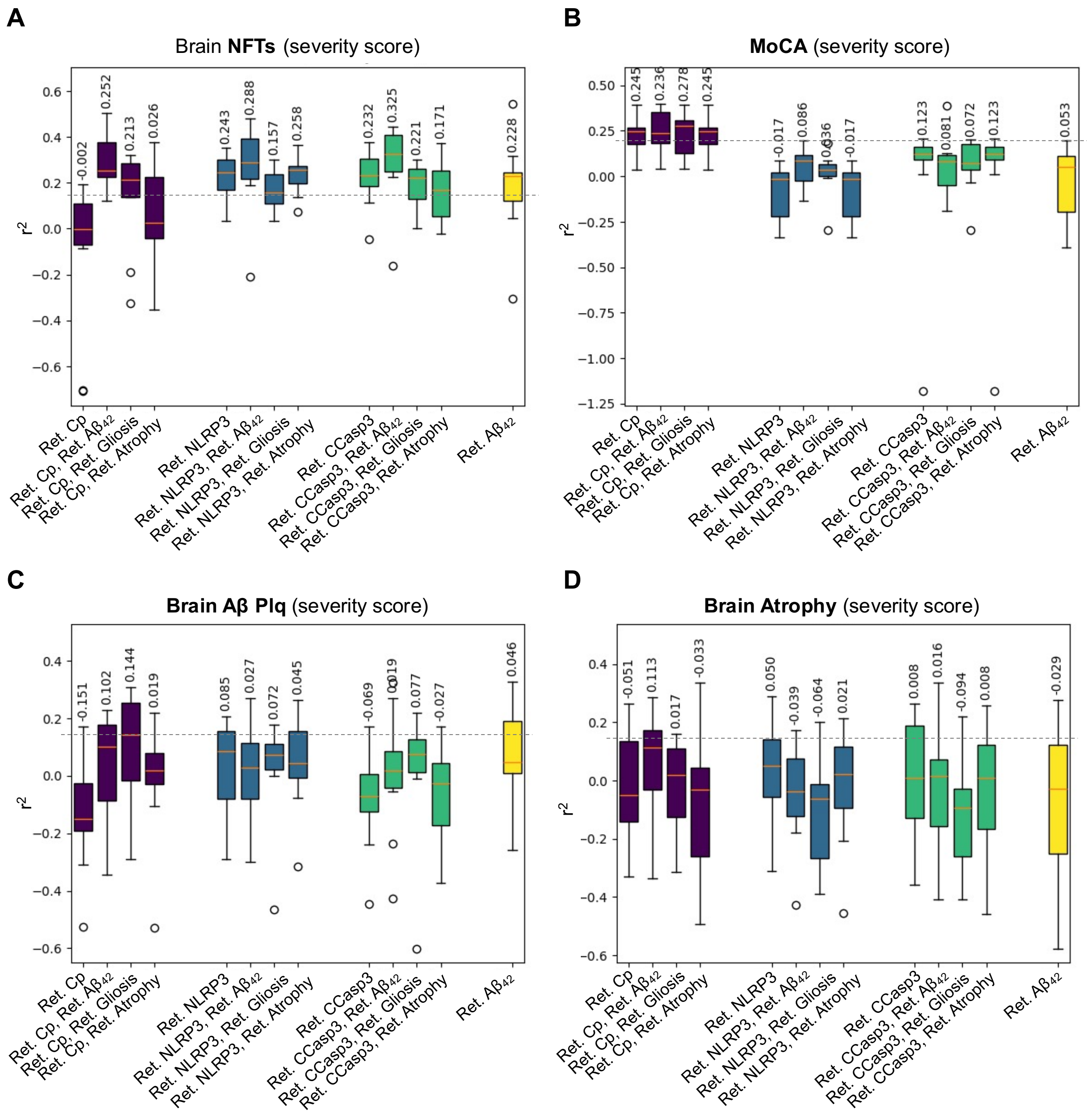


**Supplementary Figure 9. Prediction of brain AD pathologies by retinal Cp, NLRP3, cleaved caspase-3, and A$\beta_{42}$.**

**A-D.** Machine learning algorithm with a Random Forest regressor using 80 estimators was trained on the data to predict several brain pathologies and cognitive status, including (**A**) brain NFT severity score, (**B**) Montreal cognitive assessment (MoCA) score, (**C**) total brain Aβ plaques severity score, and (**D**) brain atrophy severity score. The distributions show the spread of models trained on different folds of the 5x2 cross-validation. The mean $r^2$ for each model is shown at the top of each box plot. Only models performing with variance coefficient $r^2 > 0.15$ were retained.

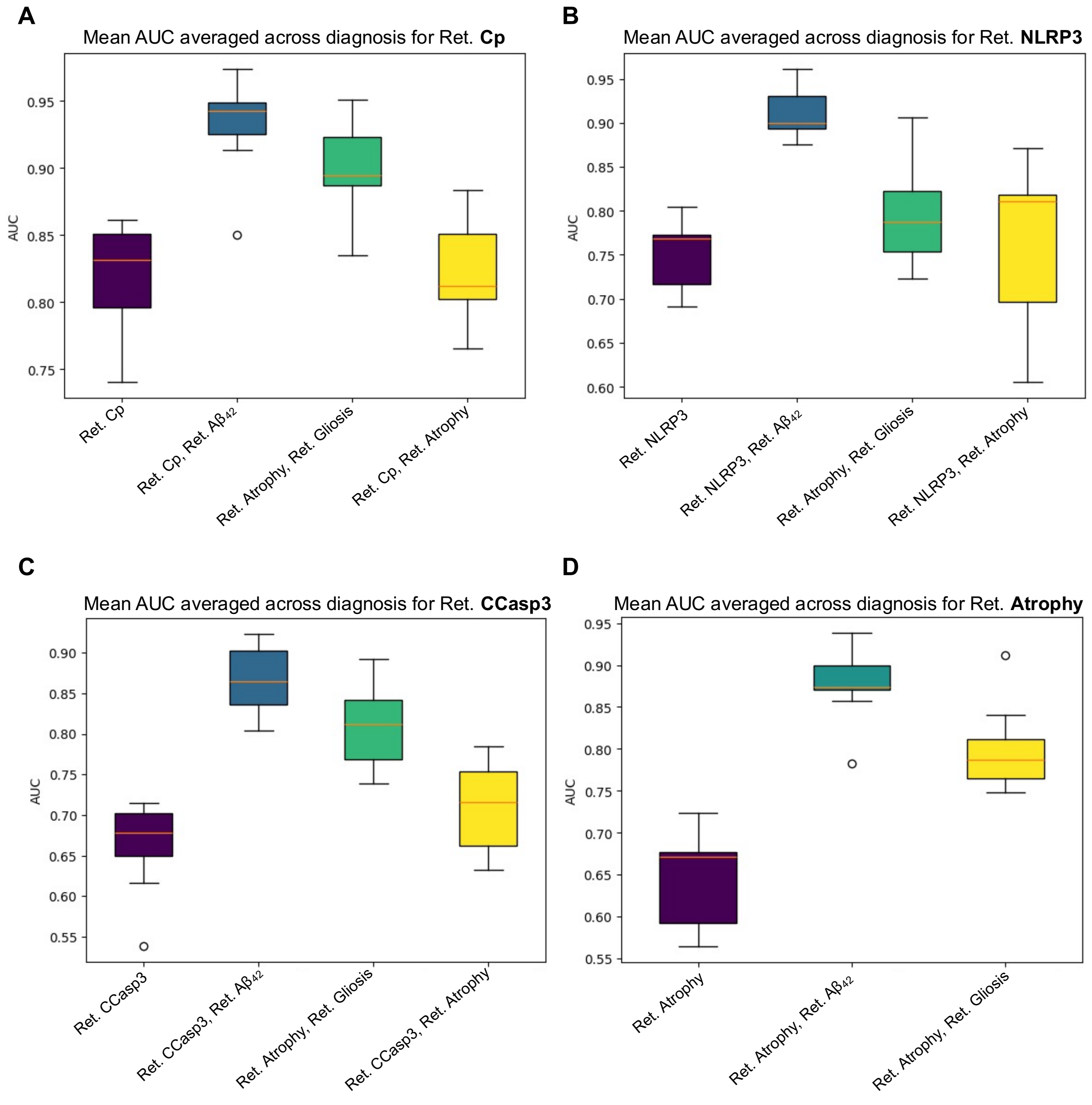


**Supplementary Figure 10. The AUC box plots across all diagnostic groups**.

**A-D.** Box plots representing the AUC measure for (**A**) retinal Cp, (**B**) retinal NLRP3, (**C**) retinal CCaspase3, and (**D**) retinal atrophy for all diagnostic groups combined. For each model, AUC was measured either individually or combined with retinal $A\beta_{42}$, or retinal gliosis (IBA1, GFAP, and Vimentin), or retinal atrophy.

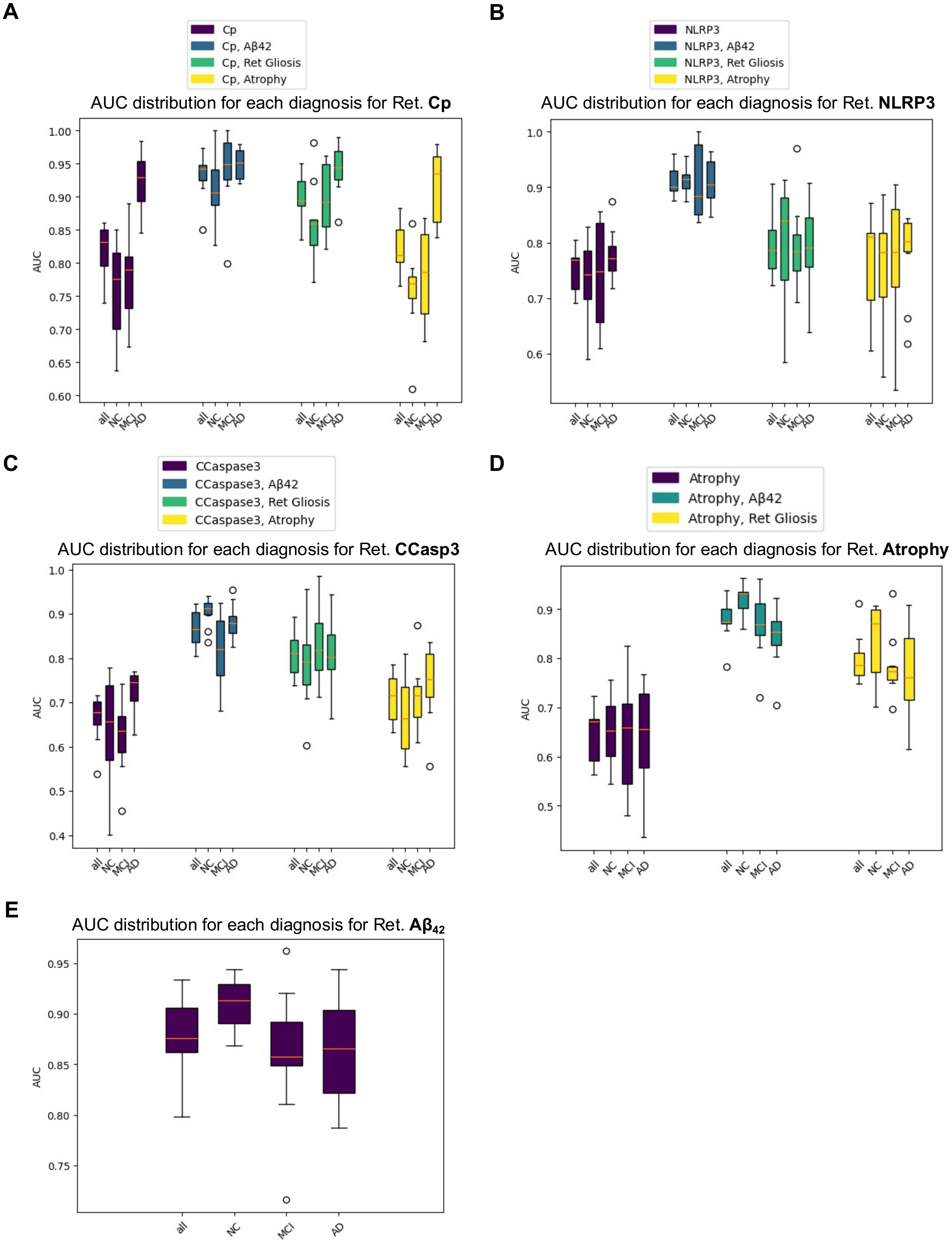
A
Cp
Cp, Aβ42
Cp, Ret Gliosis
Cp, Atrophy
AUC distribution for each diagnosis for Ret. Cp
AUC
all NC MCI AD
B
NLRP3
NLRP3, Aβ42
NLRP3, Ret Gliosis
NLRP3, Atrophy
AUC distribution for each diagnosis for Ret. NLRP3
AUC
all NC MCI AD
C
CCaspase3
CCaspase3, Aβ42
CCaspase3, Ret Gliosis
CCaspase3, Atrophy
AUC distribution for each diagnosis for Ret. CCasp3
AUC
all NC MCI AD
D
Atrophy
Atrophy, Aβ42
Atrophy, Ret Gliosis
AUC distribution for each diagnosis for Ret. Atrophy
AUC
all NC MCI AD
E
AUC distribution for each diagnosis for Ret. Aβ42
AUC
all NC MCI AD

**Supplementary Figure 11. The AUC box plots across each diagnostic groups**.

Box plots representing the AUC measure for (**A**) retinal Cp, (**B**) retinal NLRP3, (**C**) retinal CCasp3, (**D**) retinal atrophy, and (**E**) retinal $A\beta_{42}$ for each diagnostic groups NC, MCI and AD. For each model, AUC was measured either individually or combined with retinal $A\beta_{42}$, or retinal gliosis (IBA1, GFAP, and Vimentin), or retinal atrophy.
